# Rad51 determines pathway usage in post-replication repair

**DOI:** 10.1101/2024.06.14.599120

**Authors:** Damon Meyer, Shannon J. Ceballos, Steven Gore, Jie Liu, Giordano Reginato, Maria I. Cano-Linares, Katarzyna H. Maslowska, Florencia Villafañez, Christopher Ede, Vincent Pagès, Felix Prado, Petr Cejka, Wolf-Dietrich Heyer

## Abstract

Stalled replication forks can be processed by several distinct mechanisms collectively called post-replication repair which includes homologous recombination, fork regression, and translesion DNA synthesis. However, the regulation of the usage between these pathways is not fully understood. The Rad51 protein plays a pivotal role in maintaining genomic stability through its roles in HR and in protecting stalled replication forks from degradation. We report the isolation of separation-of-function mutations in *Saccharomyces cerevisiae* Rad51 that retain their recombination function but display a defect in fork protection leading to a shift in post-replication repair pathway usage from HR to alternate pathways including mutagenic translesion synthesis. Rad51-E135D and Rad51-K305N show normal *in vivo* and *in vitro* recombination despite changes in their DNA binding profiles, in particular to dsDNA, with a resulting effect on their ATPase activities. The mutants lead to a defect in Rad51 recruitment to stalled forks *in vivo* as well as a defect in the protection of dsDNA from degradation by Dna2-Sgs1 and Exo1 *in vitro*. A high-resolution cryo-electron microscopy structure of the Rad51-ssDNA filament at 2.4 Å resolution provides a structural basis for a mechanistic understanding of the mutant phenotypes. Together, the evidence suggests a model in which Rad51 binding to duplex DNA is critical to control pathway usage at stalled replication forks.

## Introduction

Homologous recombination (HR) provides a central pathway to maintain genomic stability (Kowalczykowski et al., 2016). In the context of double-strand break (DSB) repair in mitotically growing cells, HR uses the intact sister chromatid as a template for DNA repair synthesis (Li and Heyer, 2008). Initially, the DSB is resected in a 5’-3’ direction to generate 3’-OH ending single-stranded DNA tails (Cejka and Symington, 2021). Resection is initiated by the Mre11 nuclease in the Mre11-Rad50-Xrs2 complex, which, in conjunction with Sae2, can also process chemically complex DSB including those covalently bound to proteins. After an initial endonucleolytic incision and 3’-5’ resection by the Mre11 nuclease, two long-range nuclease pathways operate in the 5’-3’ direction including the dsDNA-specific exonuclease Exo1 and the ssDNA nuclease Dna2, which functions in conjunction with the helicase-topoisomerase complex Sgs1-Top3-Rmi1 that generates the ssDNA substrate for Dna2. Resection involves 100s if not 1,000s of nucleotides of ssDNA which is bound by the ssDNA-binding protein RPA. Nucleation of Rad51 filaments on RPA-coated ssDNA is achieved by mediator proteins, mainly Rad52 in budding yeast, a role that is usurped by the BRCA2 tumor suppressor protein in mammalian cells (Kowalczykowski, 2015). The resulting Rad51-ssDNA filament catalyzes the signature HR steps of homology search and DNA strand invasion to generate displacement loops (D-loops) and prime repair DNA synthesis in conjunction with the dsDNA motor protein Rad54 (Kowalczykowski, 2015). In budding yeast, Rad54 is essential to D-loop formation *in vivo* and *in vitro* (Petukhova et al., 1998; Piazza et al., 2019). The HR pathway, including Rad51, Rad52, and Rad54, also supports DNA replication by recovering broken replication forks in a process called break-induced replication and through the repair of replication-associated single-stranded gaps that occur after repriming (Arbel et al., 2021; Branzei and Szakal, 2016; Deem et al., 2011; Kockler et al., 2021).

Rad51 is the eukaryotic homolog of bacterial RecA with highly similar functions in HR (Aboussekhra et al., 1992; Kowalczykowski, 2015; Shinohara et al., 1992). RecA, however, has at least two additional functions in bacteria. It acts as a cofactor during translesion synthesis by DNA PolV and it triggers the SOS response, a bacterial DNA damage response signaling pathway initiated by RecA-bound ssDNA (Maslowska, 2019; Patel et al., 2010). Currently, no data suggests that Rad51 plays a similar role in TLS in eukaryotes. A function of Rad51 bound to ssDNA in adaption to DNA damage has been postulated, but the mechanism of this potential role in DNA damage signaling has not been defined (Lee et al., 2003). Eukaryotic Rad51 proteins bind ssDNA and dsDNA with similar affinity and only a slight preference for ssDNA (Bianco et al., 1998; Chi et al., 2006; Tombline et al., 2002; Zaitseva et al., 1999). This is in strong contrast to bacterial RecA, which strongly prefers binding to ssDNA (Kowalczykowski, 1991; Kowalczykowski et al., 1987). Binding to ssDNA by Rad51 is essential for HR, whereas Rad51 binding to duplex DNA inhibits HR, providing the rationale for proteins that mediate the assembly of Rad51 filaments on ssDNA and the dissociation of Rad51 from dsDNA (Daley et al., 2014; Mason et al., 2015; Shah et al., 2010; Solinger et al., 2002; Sung and Robberson, 1995; Zaitseva et al., 1999). Rad51 binding to chromosomal duplex DNA may also be limited by chromatin and accessibility. This poses the question, why did high-affinity dsDNA binding evolve in eukaryotic Rad51 and what is its function?

An independent role of some HR proteins in response to replication stress has been identified in vertebrate cells in protecting nascent DNA at replication forks from pathological degradation by MRE11, DNA2 or EXO1 (Berti et al., 2020; Hashimoto et al., 2010; Schlacher et al., 2011). Blockage of the replicative helicase or DNA polymerases and regressed forks generate substrates for these nucleases in the form of ssDNA/dsDNA junctions in gaps and the dsDNA ends in the regressed replication forks. The HR proteins required for fork protection are BRCA2 and RAD51 but not RAD54 (Berti et al., 2020; Schlacher et al., 2011), suggesting that RAD51 filament formation but not the entire HR process is required for fork protection. This is consistent with separation-of-functions mutants in human RAD51 and BRCA2 that affect differentially the HR and fork protection functions (Mason et al., 2019; Schlacher et al., 2011). The mechanisms involved and the exact binding sites for BRCA2 and RAD51 to protect stalled replication forks are not fully understood and recent work suggested an involvement of RAD51 binding to dsDNA (Halder et al., 2022; Liu et al., 2023). It is unclear how fork protection relates to the response to replication stress in yeast.

Replication stress and fork stalling can be elicited by unusual DNA structures, protein-DNA complexes, limiting nucleotide pools, interference from transcription as well as endogenously and exogenously induced DNA damage (Berti et al., 2020). Stalling at DNA damage sites on the leading and lagging DNA strands can be overcome by direct translesion synthesis (TLS) or downstream repriming leaving a post-replicative gap (Arbel et al., 2021; Berti et al., 2020; Branzei and Psakhye, 2016). Such a gap can either be closed by TLS or a homologous recombination (HR)-based mechanism of template switching (TS) using the sister chromatid as a donor. These processes may be spatially and temporally uncoupled from the replication fork (Daigaku et al., 2010; Gonzalez-Prieto et al., 2013; Karras and Jentsch, 2010; Lettier et al., 2006; Wong et al., 2020). TLS skips the lesion through the engagement of specialized DNA polymerases (Goodman and Woodgate, 2013). In yeast, a key TLS polymerase is Pol zeta (Pol ζ), a four subunit assembly of Pol31, Pol32, Rev7 and the catalytic subunit Rev3 (Makarova and Burgers, 2015). TLS is supported by monoubiquitylation of PCNA at lysine 164 by the Rad6 E2 ubiquitin conjugase in concert with the Rad18 E3 ubiquitin ligase (Hoege et al., 2002; Stelter and Ulrich, 2003). HR is critical for TS and involves the formation of Rad51-ssDNA filaments that perform homology search and DNA strand exchange in conjunction with its co-factors the Rad51 paralogs Rad55-Rad57 and the dsDNA motor protein Rad54 (Arbel et al., 2021; Branzei and Szakal, 2016; Giannattasio et al., 2014).

Unique to fork stalling at leading strand lesion is the process of fork regression, another mechanism of TS, where the fork regresses into a four-stranded structure called a chicken foot. Yeast Rad5 was shown to promote fork regression *in vitro* (Blastyak et al., 2007). However, *in vivo*, DNA damage checkpoint signaling suppresses fork regression in yeast and direct electron microscopic analysis detected few regressed forks in wild type cells after the addition of hydroxy urea (HU), an inhibitor of ribonucleotide reductase that leads to fork stalling (Sogo et al., 2002).

In contrast, in mammalian cells, fork regression is frequently observed in response to various fork stalling agents including HU, methyl methanesulfonate (MMS) and UV, and this process is dependent on the central HR protein RAD51 (Zellweger et al., 2015) as well as the human RAD5 homolog HLTF (Blastyak et al., 2010). Contra-directional migration of the regressed fork can reestablish a replication fork with the original lesion being avoided. In human cells, fork regression and restart of regressed forks are controlled by a complex ensemble of dsDNA translocases (ZRANB3, SMARCAL1, HLTF) and DNA helicases (FBH1, RECQ1) regulated by poly-ubiquitylation of PCNA (Berti et al., 2020). Alternatively, the chicken foot structure could be processed by 5’-3’ exonucleases to generate a 3’-ending ssDNA strand available for Rad51 filament formation and DNA strand invasion in front of the fork stall point to restart DNA synthesis (Carr and Lambert, 2021). In sum, TS comprises several mechanistically diverse processes, including fork regression and several forms of gap repair, which can occur at the stalled fork or behind the stalled fork (Branzei, 2011). How the usage of these different pathways is managed is currently not fully understood. One aspect is PCNA poly-ubiquitylation by the ubiquitin-conjugating complex Ubc13/Mms2 and Rad5 (Branzei, 2011), but it is not clear whether all TS pathways are affected. Consistently, inactivation of PCNA poly-ubiquitylation by a *ubc13* mutation inhibits TS leading to a strong increase in the use of the TLS pathway (Maslowska et al., 2019). Interestingly, Rad5 also provides an additional pathway to recruit TLS polymerases to stalled forks (Fan et al., 2018; Gallo et al., 2019; Gangavarapu et al., 2006; Maslowska and Pagès, 2022; Xu et al., 2016). Another regulator in budding yeast appears to be the Srs2 helicase, which has the ability to strip Rad51 from ssDNA, an activity that is counteracted by Rad55-Rad57 (Krejci et al., 2003; Liu et al., 2011a; Veaute et al., 2003). Interestingly, Srs2 is recruited to stalled forks by sumoylation of PCNA on lysines 127 and 164 thanks to its C-terminal SUMO binding motif (Papouli et al., 2005; Pfander et al., 2005). The genetic observations that sensitivity to fork stalling agents caused by mutations in Rad5, Rad6 and Rad18 that abolish TLS and TS are largely suppressed by mutations in Srs2 are consistent with a model that Srs2 suppresses activation of an HR-dependent TS pathway independent of PCNA ubiquitylation (also termed salvage pathway) when recruited by SUMO-PCNA (Aboussekhra et al., 1989; Lawrence and Christensen, 1979; Schiestl et al., 1990). In sum, HR-dependent and independent TS as well as TLS pathways operate at regressed forks in yeast, but it remains unclear what regulates their usage (Broomfield et al., 2001; Smirnova and Klein, 2003).

To define what controls pathway usage at stalled replication forks in the budding yeast *Saccharomyces cerevisiae*, we isolated the separation of function mutants in *RAD51*, which are largely intact for their HR function but severely affect pathway usage at replication forks stalled by the alkylating agent MMS. The mutants display sensitivity to MMS and their survival strongly depends on other post-replication repair pathways, namely TLS and Rad51-independent template switching (fork regression). We directly show that pathway usage at fork stalling lesions is dramatically changed. The Rad51 mutant proteins are poorly recruited to stalled forks *in vivo* and cannot protect dsDNA from exonucleolytic attack *in vitro*. Biochemical analysis identified a defect in dsDNA binding as the major characteristic that correlated between the *in vitro* and *in vivo* phenotypes of the mutants. We suggest that Rad51 binding to dsDNA is a major determinant in post-replication repair and that the properties of the Rad51 filament are critical for pathway usage at stalled forks.

## Results

### *RAD51* mutant isolation

Rad51 protein functions as the central homology search and DNA strand invasion factor during HR, which involves filament formation on ssDNA. In an independent role, Rad51 protects stalled replication forks, but the mechanisms involved are less well defined. Rad54 is a dsDNA motor protein that is critical for HR but not required for fork protection in mammalian cells (Schlacher et al., 2011). Using sensitivity to the fork stalling agent MMS, we isolated two mutations in *RAD51*, each with a single amino acids change, *rad51-E135D* (*referred* to as *rad51-ED*) and *rad51-K305N* (referred to as *rad51-KN*), that suppress the MMS-sensitivity of *rad51 rad54* double mutants (**Fig. 1A**). In high copy when borne on a 2µ plasmid, both *rad51* mutants substantially suppressed the MMS sensitivity of a *rad51Δ rad54^ts^* double mutant at the restrictive temperature. There was also significant but less extensive suppression of the MMS sensitivity of a *rad51Δ rad54Δ* double mutant (**Fig. 1B**).

**Figure 1:**
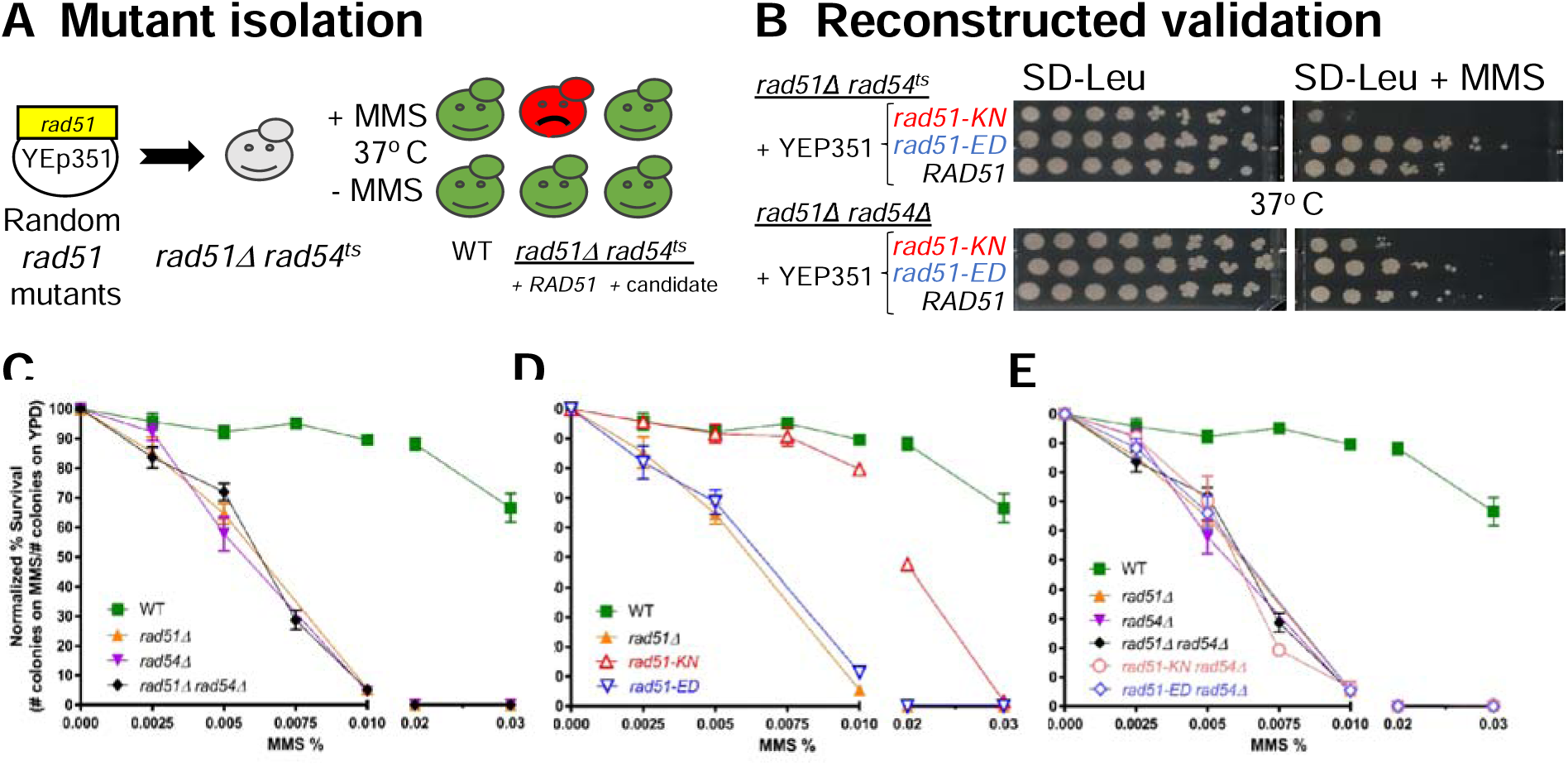
Isolation, validation, and MMS sensitivity of *rad51-ED* and *rad51-KN*. **A.** Scheme to isolate *rad51* mutants with predicted phenotypes for wild type cells (WT), *rad51Δ rad54ts* cells with wild type *RAD51* (*RAD51*) and a candidate mutant (candidate). **B.** Validation of mutant phenotype with plasmid-borne gene by transformation into fresh tester strains. Serial dilution *assays* of *rad51* mutations in the presence and absence of 0.002% MMS (*rad51Δ rad54^ts^*: WDHY2546 with pWDH957 (*RAD51*), pWDH954 (*rad51-ED*), pWDH953 (*rad51-KN*); *rad51Δ rad54Δ*: WDHY2544 with pWDH957 (*RAD51*), pWDH954 (*rad51-ED*), pWDH953 (*rad51-KN*)). **C-E.** MMS sensitivity of chromosomally integrated mutations. Survival of cells following exposure to varying levels of MMS (0.0025%, 0.005%, 0.0075%, 0.01%, 0.02%, and 0.03%) was measured as the number of colonies formed on YPD relative to unexposed cells (0% MMS) in the following genotypes: wild type (WT) (WDHY3960), *rad51Δ* (WDHY3898), *rad54Δ* (WDHY3961), *rad51Δ rad54Δ* (WDHY3959), *rad51-KN* (WDHY3962), *rad51-ED* (WDHY3548), *rad51-KN rad54Δ* (WDHY3963), and *rad51-ED rad54Δ* (WDHY3547). The mean and SEM was calculated from a minimum of three independent cultures.

### *rad51-ED* and *rad51-KN cause s*ensitivity to fork stalling by methyl methanesulfonate

To study these mutations in a more controlled fashion, we introduced them into the chromosomal *RAD51* gene and conducted survival assays through colony formation (**Fig. 1C-E**). We found the expected high sensitivity of *rad51Δ* and *rad54Δ* mutant cells to MMS and the expected epistasis between both mutations (**Fig 1C**). The *rad51-ED* mutant was as sensitive to MMS as the *rad51Δ* strain, whereas the *rad51-KN* was only mildly more sensitive than wild type cells, notable only at higher MMS doses (**Fig. 1D**). Congruent results were noted in the serial dilution plate assays (**Fig. S1A**; compare row 5 wild type and row 12 *rad51Δ* with row 2 *rad51-ED* and row 9 *rad51-KN*). Neither chromosomal mutant was able to suppress the MMS sensitivity of a *rad54Δ* mutation in the colony formation assay (**Fig. 1E**). There was some suppression in the serial dilution plate assay (**Fig. S1A**, compare row 6 *RAD51 rad54Δ* with row 4 *rad51-ED rad54Δ* and row 11 *rad51-KN rad54Δ*). In the serial dilution assays we consistently used a rather low MMS concentration (0.0033%) to compare the widely varying sensitivity of all strains on a single plate. This discrepancy between both MMS sensitivity assays is explained by the fact that serial dilution assays measure a mixture of growth and survival, whereas the colony formation assay measures survival only. The steady-state levels of the Rad51-ED and Rad51-KN mutant proteins were the same as the wild type Rad51 protein (**Fig. S1C**), eliminating the possibility that differences in protein levels are the cause of the observed phenotypes. We conclude that Rad51-ED and Rad51-KN are regularly expressed mutant proteins with altered functions in the cellular response to MMS.

### *rad51-ED* and *rad51-KN* are recombination-proficient and do not suppress the recombination defect caused by *rad54Δ*

To assess the recombination properties of the newly isolated *rad51* mutants, we employed a well-established sister chromatid recombination assay that allows monitoring of spontaneous and DSB-induced recombination between two directly repeated copies of mutant *leu2* genes that can give rise to Leu^+^ recombinants (**Fig. 2A**). The system uses *URA3* as a marker between the repeated *leu2* genes to distinguish gene conversion events (Leu^+^ Ura^+^) from other events that include single-strand annealing, intrachromatid crossing over, unequal sister-chromatid exchange, unequal sister-chromatid conversion, and replication slippage and are summarily called pop-out events (Leu^+^, Ura^-^). Spontaneous pop-out events in this system are largely mediated by Rad52-dependent strand annealing and are independent of Rad51 ((Smith and Rothstein, 1999) and **Fig. 2B left**). Neither *rad51Δ* and *rad54Δ* nor *rad51-ED* and *rad51-KN* mutations affected these events. The Leu^+^ Ura^+^ gene conversion events strongly depended on Rad51, Rad52, and Rad54, as expected (**Fig. 2B right**). The rates of Leu^+^ Ura^+^ gene conversion events in the *rad51-ED* and *rad51-KN* mutants were indistinguishable from the wild type rate. The mutants did not suppress the recombination defect caused by the *rad54Δ* mutation (**Fig. 2B right**). The frequency of both events (Leu^+^ Ura^+^, Leu^+^ Ura^-^) can be dramatically stimulated by the induction of a DNA double-stranded break using a galactose-controlled version of the HO-endonuclease which cleaves a HO recognition site engineered between the repeated mutant *leu2* genes (**Fig. 2A**). In determining the frequencies of DSB-induced pop-out (Leu^+^ Ura^-^) and gene conversion (Leu^+^ Ura^+^) events, the results showed that the *rad51-ED* and *rad51-KN* mutants exhibited no interference with pop-out events, no defect in gene conversion, and no suppression of the *rad54Δ* gene conversion defect (**Fig. 2C**). This exactly mirrors the results for spontaneous HR events. We conclude that *rad51-ED* and *rad51-KN* are proficient for mitotic homologous recombination.

**Figure 2:**
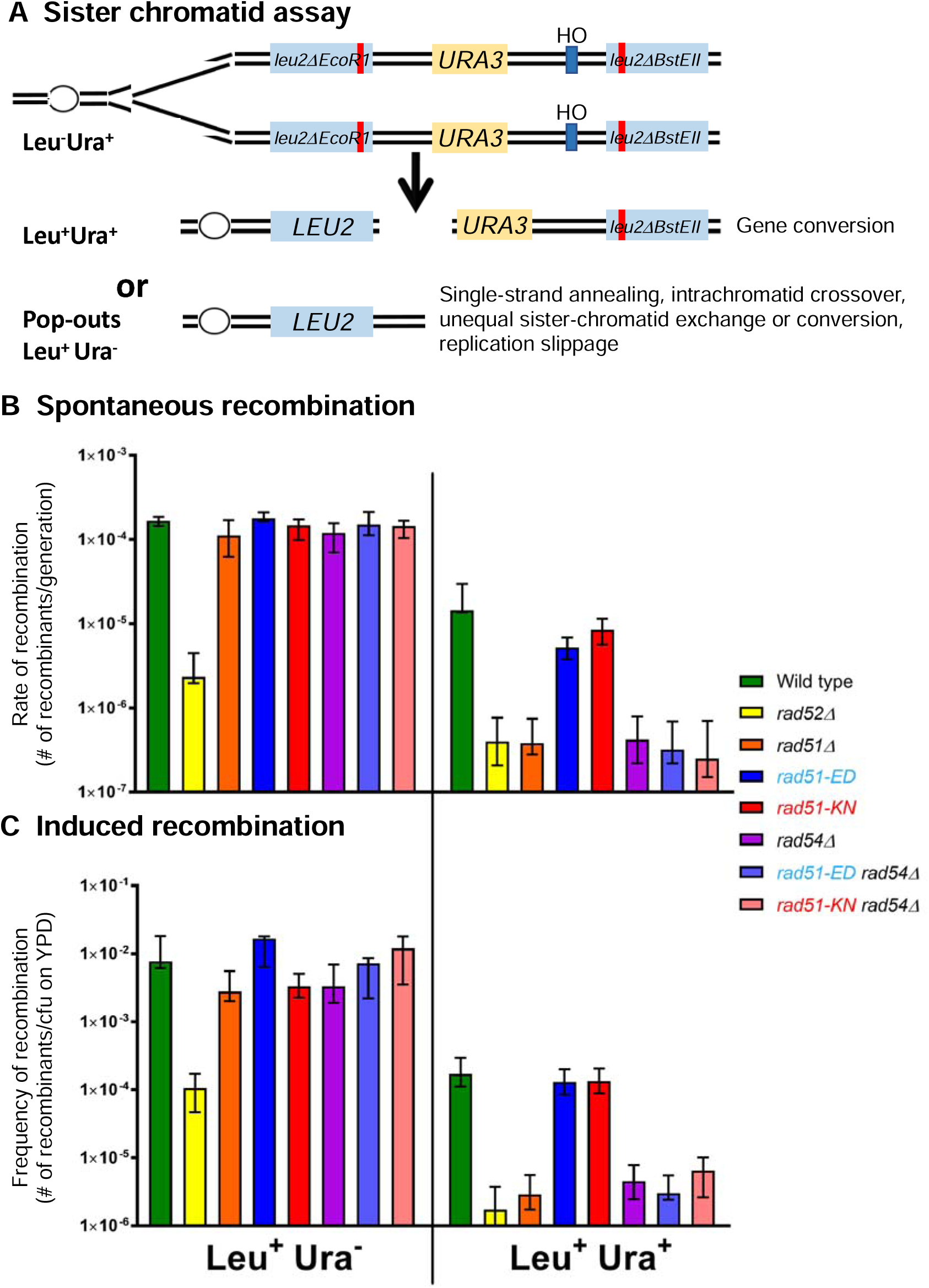
*rad51-ED* and *rad51-KN* are recombination-proficient and cannot suppress the recombination defect of *rad54Δ*. **A.** Direct repeat recombination assay (Smith and Rothstein, 1999). HO: HO endonuclease cleavage site**. B.** Spontaneous recombination rate data. **C.** Recombination frequency data after DSB induction. Strains were freshly dissected from diploid strains and single spore clones were assayed, a minimum of 12 single clones were assayed per genotype. The median recombination rates/frequencies are given with the 95 % confidence interval (Lea and Coulson, 1949). Diploids allowed for multiple strains to be created by dissection. Wildtype (WDHY3383), *rad52*Δ (WDHY3348), *rad51*Δ (WDHY3915)*, rad51-KN* (WDHY3385), *rad51-KN rad54*Δ (WDHY3463), *rad51-ED* (WDHY3777), *rad51-ED rad54*Δ (WDHY3462), *rad54*Δ (WDHY3349). The viability data are shown in **Figure S2**. The data are all relative to the number of colony forming units (cfu) on YPD.

While *rad52Δ* showed a significant survival defect after DSB induction by HO, *rad51Δ* and *rad54Δ* showed a smaller survival defect (**Fig. S2A**), as expected. This reflects the involvement of Rad52 in both pop-out and gene conversion events, whereas pop-out events are unaffected in *rad51Δ* and *rad54Δ* cells, allowing substantial DSB repair. The *rad51-ED* mutant shows slightly better survival after DSB induction than *rad51Δ* cells which was within the error of the experiment, whereas *rad51-KN* cells survived significantly better than *rad51Δ* cells exceeding the survival of wild type cells (**Fig. S2A**). This difference in response to a single DSB between both mutants was also recapitulated by their differential sensitivity to 100 Gy of ionizing radiation in serial dilution plate assays, where *rad51-ED* was IR sensitive but somewhat less so than *rad51Δ* (**Fig. S1B**)*. rad51-KN* showed resistance to IR that was equivalent to wild type cells (**Fig. S1B**). We conclude that *rad51-ED* has a survival defect in response to a single DSB or IR despite being apparently fully proficient in HR. This may be related to the signaling role of DNA bound Rad51 affecting adaptation from DNA damage (Lee et al., 2003). The improved survival of the *rad51-KN* mutant after a single DSB may also be related to this signaling role. IR also induces base damages that may elicit replication fork stalling leading to a possible alternate interpretation that the IR-sensitivity of the mutants is related to fork protection (see below), although we did not further explore the IR phenotype.

### *rad51-ED* and *rad51-KN* display a strong mutator phenotype similar to a complete HR defect

HR-defective cells such as *rad51Δ* or *rad54Δ* mutants display a considerable mutator phenotype likely caused by spontaneous DNA damage that, instead of being repaired or tolerated by HR, is processed by the error-prone TLS pathway, causing an increase in spontaneous mutagenesis. We used the forward mutagenesis *CAN1* assay to determine the rate of spontaneous mutations and saw the expected 9-10-fold increase in *rad51Δ, rad54Δ* single and double mutants (**Fig. 3**). It was previously reported that this increase largely depends on the central TLS polymerase component Rev3 (Loeillet et al., 2020). To our surprise, *rad51-ED* and *rad51-KN* showed a significant increase of 9-fold and 6-fold, respectively, in their mutation rate. These values are not significantly different from those of the *rad51Δ, rad54Δ* single and double mutants. Both mutants did not suppress the mutator phenotype of the *rad54Δ* mutant (**Fig. 3**). Instead, *rad51ED* slightly elevated the mutation rate in *rad54Δ* cells from 9-fold in the single mutants to 24-fold in the *rad51-ED rad54Δ* double mutant, which suggests a non-epistatic relationship. We conclude that *rad51-ED* and *rad51-KN* show a mutator phenotype equivalent to a complete HR defect despite being fully HR proficient.

**Figure 3.**
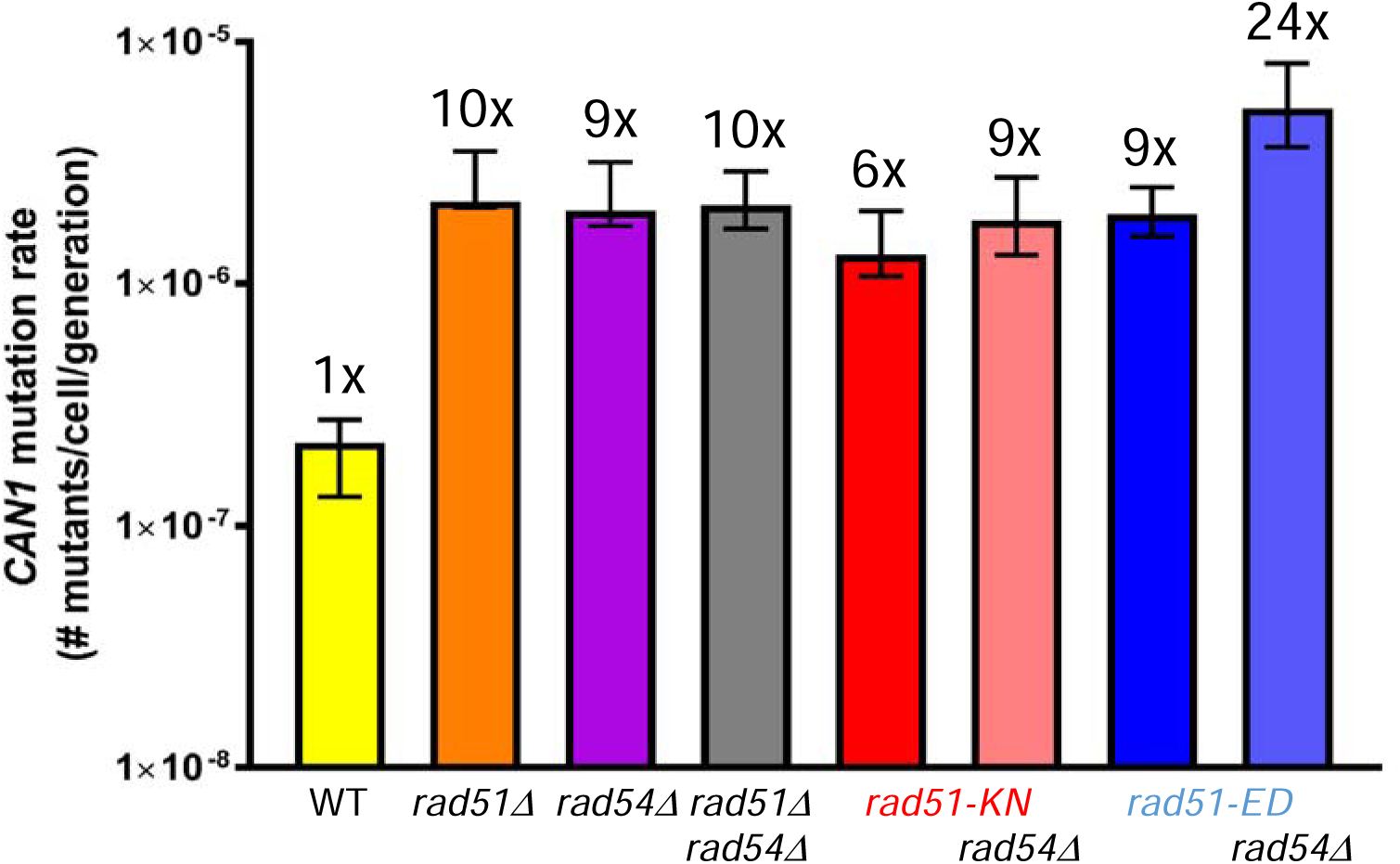
*rad51-ED* and *rad51-KN* display a strong mutator phenotype similar to a complete HR defect. The spontaneous *CAN1* mutation rate from a minimum of 12 single clones was assayed. Freshly dissected diploid strains allowed for multiple strains of the following genotypes to be created: WT and *rad51*Δ (WDHY4204)*, rad54*Δ (WDHY4205), *rad51Δ rad54*Δ (WDHY3162), *rad51-KN* (WDHY4208), *rad51-KN rad54*Δ (WDHY4210), *rad51-ED* (WDHY4207), and *rad51-ED rad54*Δ (WDHY4209). The median rate and ± 95% confidence interval were calculated for each genotype. Fold increase compared to WT is given.

### *rad51-ED* and *rad51-KN* display strong synergy with defects in post-replication repair

The significant increase in spontaneous mutagenesis could signal a shift from high-fidelity damage tolerance by HR to error-prone bypass *via* TLS at stalled replication forks. We explored the genetic interactions of *rad51-ED* and *rad51-KN* with mutations affecting various aspects of post-replication repair. We first tested a deletion mutation of *REV3*, which by itself shows no sensitivity to MMS at the low levels (0.0033%) used for the serial dilution assays in this experiment (**Fig. 4A** row 3). This contrasts with the significant sensitivity of HR-defective mutants (*rad54Δ* row 2, *rad51Δ* row 5, *rad51Δ rad54Δ* row 6). An HR defect displayed strong synergy with a defect in *REV3* leading to extreme MMS sensitivity and also affecting growth/survival on YPD growth media without MMS (*rad54Δ rev3Δ* row 4, *rad51Δ rev3Δ* row 7, *rad51Δ rad54Δ rev3Δ* row 8). These results were expected and show that HR and TLS represent parallel competing pathways to process MMS-induced DNA damage, and the single mutant phenotype suggests that HR plays a more prominent role in wild type cells than TLS. *rad51-ED* showed the MMS sensitivity expected from the previous experiments (**Fig. 1D, S1A**) that rivals that of a complete HR defect (**Fig. 4A** compare *rad54Δ* row 2, *rad51Δ* row 5, *rad51Δ rad54Δ* row 6 with *rad51-ED* row 9). The *rad51-ED rev3Δ* double mutant showed extreme MMS sensitivity and a growth/survival defect on YPD, like a complete HR defect (**Fig. 4A** compare *rad51-ED* row 9 with *rad51-ED rev3Δ* row 11). The addition of a *rad54Δ* mutation to these strains (rows 10, 12) did not further increase MMS sensitivity or the growth/survival defect on YPD. The low MMS concentration (0.0033%) was needed to compare the widely varying sensitivity of all strains on a single plate. *rad51-KN* showed little to no MMS sensitivity in the serial dilution assays (**Fig. 4B** row 9), as expected from **Figure 1**, where this mutant showed MMS sensitivity only at high MMS concentrations. Remarkably, *rad51-KN* showed synergy with *rev3Δ* (**Fig. 4** row 11), although none of the single mutant was measurably sensitive at the concentration used (rows 3, 9). These results show that MMS-stalled forks in *rad51-ED* cells are preferentially processed by TLS similar to the situation in cells with a complete HR defect, although this mutant is capable of normal HR. The *rad51-KN* mutant shows similar but less pronounced phenotypes.

**Figure 4.**
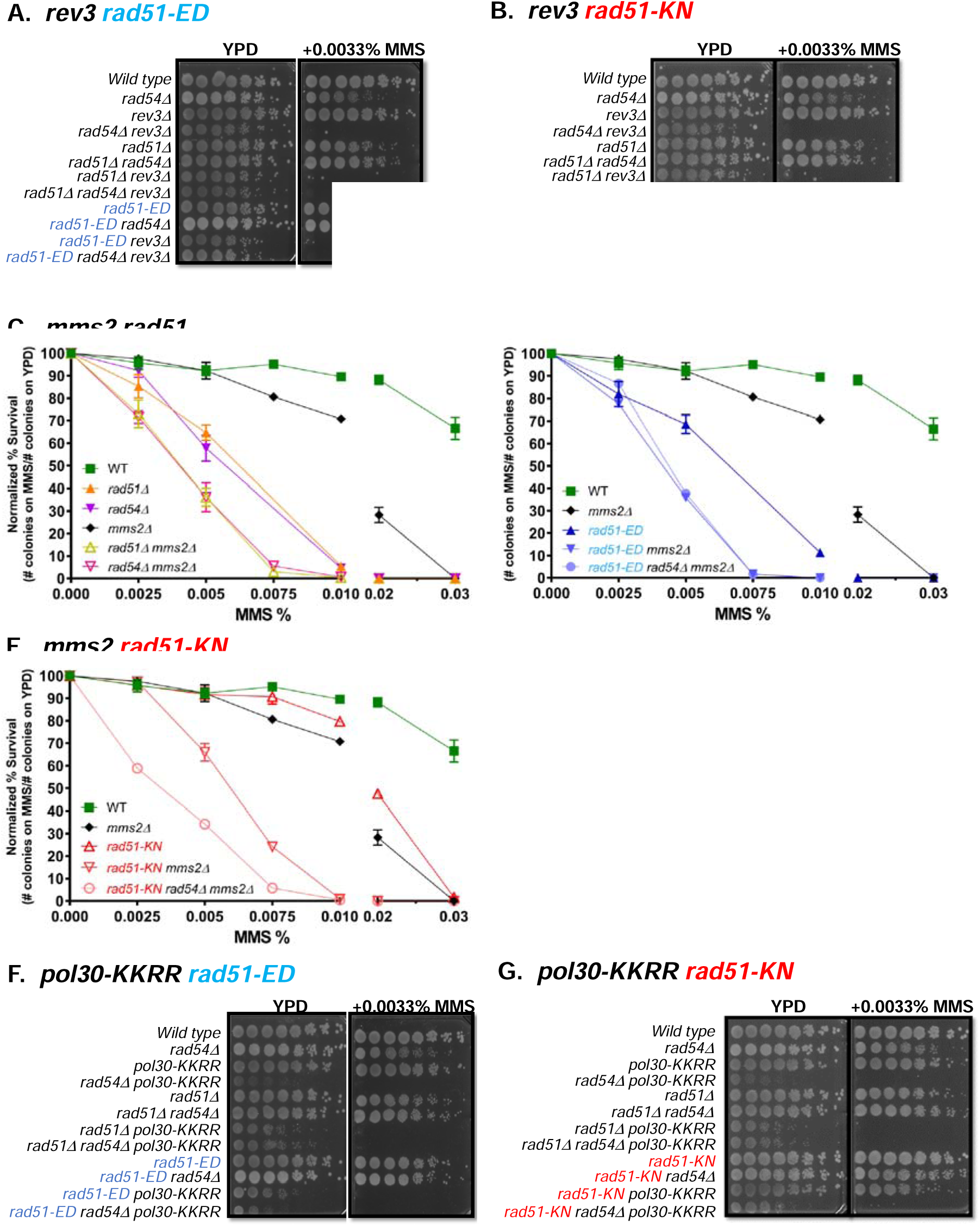
*rad51-ED* and *rad51-KN* display strong synergy with defects in post-replication repair. **A, B.** Analysis with *rev3.* Serial dilutions of wild type (WDHY1636), *rad54Δ* (WDHY1275), *rev3Δ* (WDHY3584), *rad54Δ rev3Δ* (WDHY3588), *rad51Δ* (WHDY2542), *rad51Δ rad54Δ* (WDHY2544), *rad51Δ rev3Δ* (WDHY3787), *rad51Δ rad54Δ rev3* (WDHY3788), *rad51-ED* (WDHY3548), *rad51-ED rad54Δ* (WDHY3546), *rad51-ED rev3Δ* (WDHY3784), *rad51-ED rad54Δ rev3Δ* (WDHY3785), *rad51-KN* (WDHY3962), *rad51-KN rad54Δ* (WDHY3572), *rad51-KN rev3Δ* (WDHY3589), *rad51-KN rad54Δ rev3Δ* (WDHY3789). **C-E.** Analysis of survival following exposure to MMS in the following genotypes: wildtype (WT) (WDHY3960), *rad51Δ* (WDHY3898), *rad54Δ* (WDHY3961), *mms2Δ* (WDHY3855), *rad51Δ mms2Δ* (WDHY3952), *rad54Δ mms2Δ* (WDHY3956), *rad51-KN* (WDHY3962), *rad51-KN mms2Δ* (WDHY3958), *rad51-KN rad54Δ mms2Δ* (WDHY4564), *rad51-ED* (WDHY3548), *rad51-ED mms2Δ* (WDHY5169), *rad51-ED rad54Δ mms2Δ* (WDHY6123). The mean and SEM were calculated from a minimum of three independent cultures. **F, G.** Analysis with *pol30-KKRR.* Serial dilutions of wild type (WDHY1636), *rad54Δ* (WDHY1275), *pol30-KKRR* (WDHY3555), *rad54Δ pol30-KKRR* (WDHY3563), *rad51Δ* (WHDY2542), *rad51Δ rad54Δ* (WDHY2544), *rad51Δ pol30-KKRR* (WDHY3783), *rad51Δ rad54Δ pol30-KKRR* (WDHY3780), *rad51-ED* (WDHY3548), *rad51-ED rad54Δ* (WDHY3546), *rad51-ED pol30-KKRR* (WDHY3561), *rad51-ED rad54Δ pol30-KKRR* (WDHY3570), *rad51-KN* (WDHY3962), *rad51-KN rad54Δ* (WDHY3572), *rad51-KN pol30-KKRR* (WDHY3557), *rad51-KN rad54Δ pol30-KKRR* (WDHY3568).

Next, we tested the role of Mms2, which is required for polyubiquitylation of PCNA at K164 (Hoege et al., 2002) and template switching. In quantitative survival assays, we confirmed the relative mild MMS survival defect of *mms2Δ* cells compared to HR-defective cells (*rad51Δ, rad54Δ*). Analysis of the *rad51Δ mms2Δ* and *rad54Δ mms2Δ* double mutants revealed a non-epistatic relationship between an HR defect and a loss of Mms2 function (**Fig. 4C**). These results show that in wild type cells HR makes a stronger contribution to MMS survival than pathway(s) dependent on PCNA poly-ubiquitylation. Moreover, these results suggest that Mms2 controls a strand exchange-independent template switching mechanism, in analogy to mammalian cells (Ciccia et al., 2012). This mechanism is unlikely to be HR-mediated gap repair that is Rad51 dependent; alternatively, it could be fork regression operating in yeast in a Rad51-independent manner. *rad51-ED* showed the MMS sensitivity expected from the previous experiments (**Fig. 1D)** that rivals that of a complete HR defect. *rad51-ED* showed the same increase in MMS sensitivity in the double mutant with *mms2Δ* as the *rad51Δ* strain that remained unchanged by adding a *rad54Δ* mutation (**Fig. 4D**). *rad51-KN* showed little MMS sensitivity as expected from **Figure 1**, which was similar to the MMS sensitivity of the *mms2Δ* single mutant (**Fig. 4E**). Remarkably, *rad51-KN* showed strong synergy with *mms2Δ* that was further enhanced by fully eliminating HR through a *rad54Δ* mutation (**Fig. 4F**). We conclude that MMS-stalled forks in *rad51-ED* cells have a higher propensity to be processed by a Mms2-dependent pathway (hypothetically fork regression) similar to the situation in cells with a complete HR defect. The *rad51-KN* mutant shows a less accentuated phenotype.

Furthermore, we tested a PCNA mutant (encoded by the *POL30* gene in budding yeast), which eliminated the K127 and K164 acceptor sites (*pol30-K127R,K169R* abbreviated as *pol30-KKRR*) for ubiquitylation and sumoylation to obstruct Srs2-mediated anti-recombination, TLS, and TS (likely fork regression) pathways. This double mutant displayed only very mild MMS sensitivity at the low (0.0033%) concentration used (**Fig. 4F** row 3), as expected (Hoege et al., 2002). This result suggests that under these conditions the salvage HR is the predominant pathway. This notion is confirmed by the extreme synergy in MMS sensitivity with an HR defect as well as the strong growth/survival defect on YPD (**Fig. 4F**, compare *pol30-KKRR* row 3 with *rad54Δ pol30-KKRR* row 4 or *rad51Δ pol30-KKRR* row 7 or *rad51Δ rad54Δ pol30-KKRR* row 8). *rad51-ED* mimicked again the behavior of a complete HR defect (**Fig. 4F**; compare *rad51-ED* row 9 with *rad51-ED pol30-KKRR* row 11) including the remarkable growth/survival defect on YPD lacking MMS. Consistent with the genetic interaction analysis involving *rev3Δ* and *mms2Δ,* the *rad51-KN* mutant shows a less pronounced phenotype which was enhanced by a complete HR defect (**Fig. 4G**, compare *rad51-KN* row 9 with *rad51-KN pol30-KKRR* row 11 and with *rad51-KN rad54Δ pol30-KKRR* row 12). We conclude that in *rad51-ED* and *rad51-KN* cells, HR-independent template switching and TLS process a larger fraction of MMS-stalled forks than in wild type cells.

Finally, we used a subtle mutation in *RAD5*, *rad5-G535R,* which by itself has little to no phenotype (Fan et al., 1996). Rad5 promotes PCNA polyubiquitylation and template-switching and also plays a role in recruiting TLS polymerases, affecting two major post-replication repair pathways. In consequence, the deletion of the *RAD5* gene causes one of the most pronounced MMS sensitivities caused by a single mutant in budding yeast, requiring a *rad5* allele with a more subtle phenotype. In serial dilution assays, we noticed that the suppression of the MMS sensitivity of *rad54Δ* by *rad51-ED* and *rad51-KN* depended on the wild type *RAD5* gene (**Fig. S1A** compare *rad51-ED rad54Δ RAD5* row 4 with *rad51-ED rad54Δ rad5-535* row 3 and *rad51-KN rad54Δ RAD5* row 10 with *rad51-KN rad54Δ rad5-535* row 11). These results reinforce the conclusion that *rad51-ED* and *rad51-KN* rely more heavily on TLS and HR-independent template switching to process stalled replication forks.

From this genetic analysis with four genes affecting single or multiple post-replication repair pathways, we conclude that the balance between post-replication repair pathways is significantly altered in the *rad51-ED* and *rad51-KN* mutants. We suggest that both mutants represent a separation of Rad51 function in HR, which is not measurably affected in spontaneous and induced recombination assays, but the mutants alter Rad51 function at stalled replication forks, which is eliminated (*rad51-ED*) or diminished (*rad51-KN*).

### *rad51-ED* dramatically shifts the balance from template switching to translesion synthesis

To investigate the effect of *rad51-ED* and *rad51-KN* mutants on lesion bypass, we took advantage of the iDamage approach (**Fig. 5A**) (Maslowska et al., 2019). In short, a plasmid containing the single lesion of interest is stably inserted at a specific locus of the yeast genome by means of the Cre recombinase and modified lox sites. Lesion bypass is monitored by counting blue and white colonies, as the lesion is inserted in the *lacZ* reporter gene (**Fig. 5A**). While this system does not allow to study MMS-induced DNA damage, we investigated the bypass of two common lesions: the thymine-thymine pyrimidine(6-4)pyrimidone photoproduct [TT(6-4)], and the N2-dG-Acetylaminofluorene (N2dG-AAF). All experiments were performed in a parental strain where NER and mismatch repair are inactivated to prevent the repair of the lesion and the DNA loop used as a strand marker, respectively. We measured a level of TLS of ∼5% and ∼17% for the TT(6-4) and the G-AAF lesions in this parental strain (**Fig. 5B**). The inactivation of *rad51* led to a decrease in the level of the template-switching pathways (HR-mediated gap repair, fork regression) and a concomitant increase in the level of TLS (**Fig. 5B**). In *rad51Δ* cells about half of the template switching events remained, suggesting a significant contribution of Rad51-independent template switching to the total number of tolerance events. This reflects the competition between template switching and TLS: when a template switching pathway is inhibited, TLS is favored. Upon introduction of the same lesion in the *rad51-ED* and *rad51-KN* mutants, we observed the same decrease in the use of template switching and the same increase in TLS (**Fig. 5B**).

**Figure 5.**
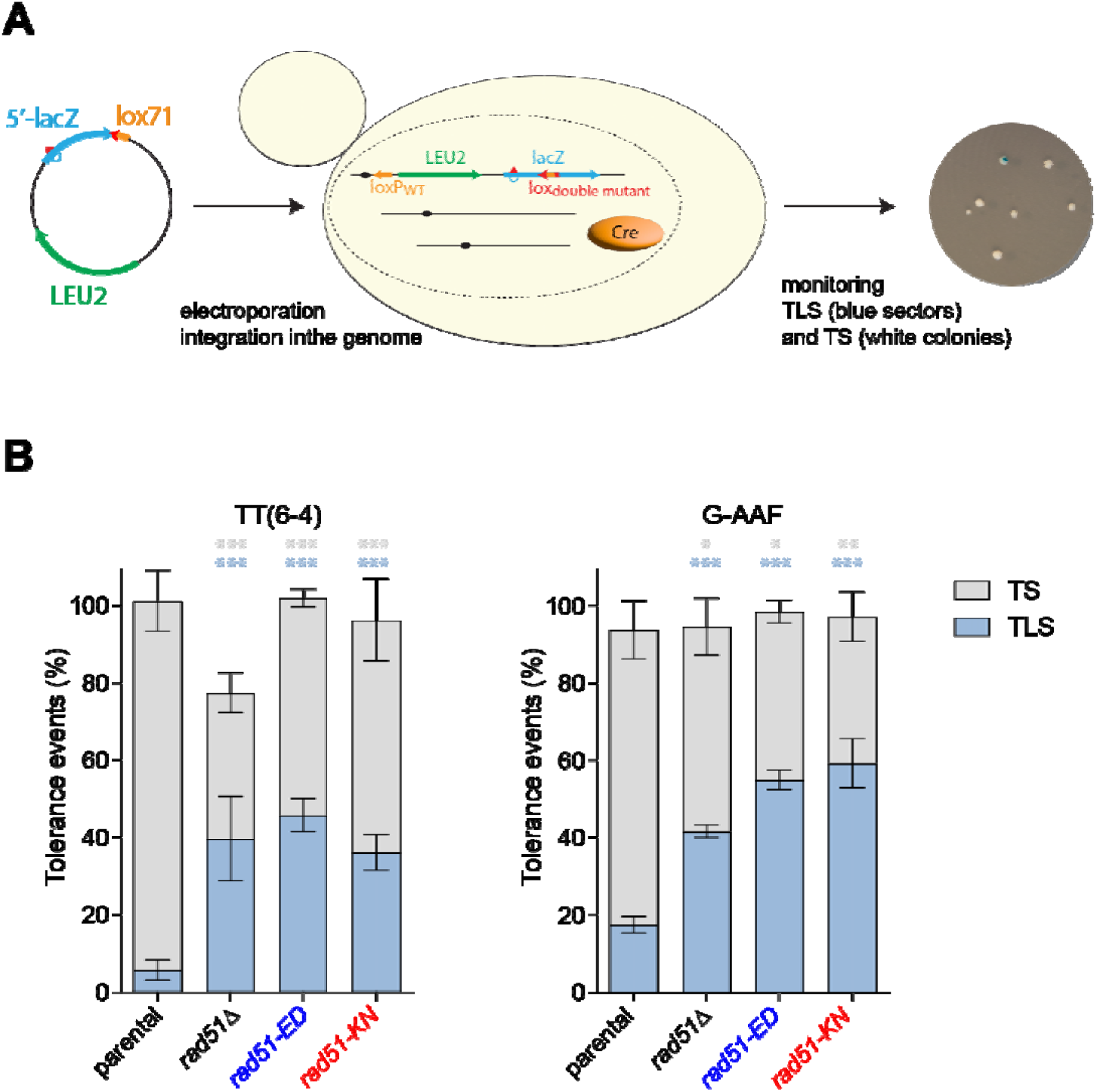
*rad51-ED* and *rad51-KN* shifts the balance from damage avoidance to translesion synthesis in post-replication repair. **A.** Outline of the iDamage approach: a non-replicative plasmid containing a single lesion is integrated into one of the yeast chromosomes using Cre/lox site-specific recombination. The integrative vector carrying a selection marker (*LEU2*) and the 5′-end of the *lacZ* reporter gene containing a single lesion is introduced into a specific locus of the chromosome with the 3′-end of *lacZ*. The precise integration of the plasmid DNA into the chromosome restores a functional *lacZ* gene, enabling the phenotypical detection of TLS and DA events (as blue and white colonies on X-gal indicator media). **B.** B. Partitioning of DNA damage tolerance pathways at a TT(6-4) photoproduct and at a G-AAF lesion. Tolerance events represent the percentage of cells able to survive in the presence of the integrated lesion compared to the lesion-free control in wild type (SC53, SC55), *rad51Δ* (SC253, SC255), *rad51-ED* (SC844, SC845) and *rad51-KN* (SC868, SC869) strains. The data represent the average and standard deviation of at least three independent experiments. Unpaired t-test was performed to compare TLS and DA values from the different mutants to the parental strain. *P < 0.05; **P < 0.005; ***P < 0.0005

### *rad51-ED* and *rad51-KN* display defects in binding to stalled forks and sister chromatid junction formation

The recombinational processing of MMS-induced DNA lesions is associated with the formation of Rad51-dependent sister-chromatid junctions (SCJs). The previous genetic results suggested that there is a defect in *rad51-ED* and *rad51-KN* cells to process MMS-stalled replication forks. The accumulation of these recombination intermediates can be detected as X-shaped structures by 2D electrophoresis in *sgs1*Δ cells, which are defective in their dissolution (Liberi et al., 2005). To address the functionality of Rad51-ED and Rad51-KN in the recombinational response to MMS, we measured the accumulation of SCJs in cells synchronized in G1 and released into S phase in the presence of MMS (**Fig. 6A**). Rad51-ED and Rad51-KN are affected in their ability to promote SCJs as inferred from the behavior of the mutants. *sgs1*Δ *rad51-KN* cells displayed a 1.7-fold reduction in the amount of SCJs along the analyzed times that was mostly due to a delay in the accumulation of SCJs as compared to the wild type. Remarkably, *sgs1*Δ *rad51-ED* was severely affected in the accumulation of SCJs and reduced to ∼20% of the wild type levels. We conclude that the *rad51-ED* and, to a lesser degree, *rad51-KN* mutants are defective in HR at MMS-stalled replication forks leading to increased usage of alternate post-replication repair pathways. Alternatively, it is possible that in these mutants such structures are less stable or more likely to be degraded.

**Figure 6.**
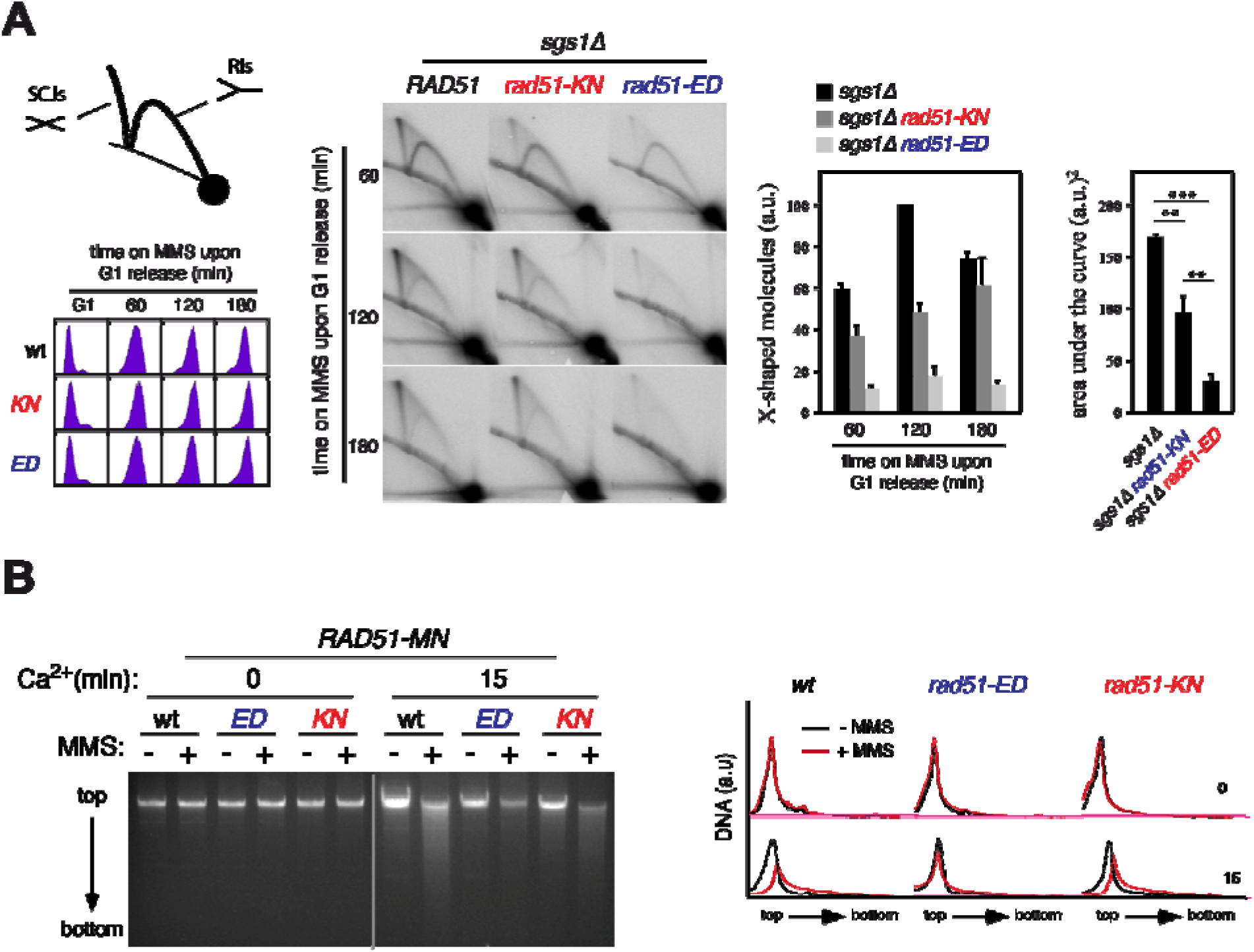
Rad51-ED is severely affected in binding to damaged DNA and Sister Chromatid Junction formation. **A.** *rad51-KN* and *rad51-ED* are defective in the accumulation of Sister Chromatid Junctions (SCJ), as determined by 2D gel analysis of X-shaped molecules in *sgs1*Δ (W303sgs1), *sgs1*Δ *rad51-KN* (W303sr305) and *sgs1*Δ *rad51-ED* (W303sr135) cells synchronized in G1 and released in the presence of 0.033% MMS. A schematic representation of the migration pattern of X-shaped (SCJs) and Y-shaped (replication intermediates) molecules, as well as the DNA content analysis of the different cultures along the time course, is shown on the left. The panel on the right shows the amounts of X-shaped molecules relative to the total amount of molecules at the indicated times. The highest value is set at 100, and the mean and SEM from four independent time courses are given. The area under the curve (AUC) is also plotted. Two and three asterisks indicate statistically significant differences according to a One-Anova (Bonferroni) test (*P* < 0.001 and 0.0001, respectively). **(B)** Rad51 binding to replicative DNA lesions as determined by ChEC analysis of exponentially growing *RAD51-MN* (wt) (wR51MN-2), *rad51-ED-MN* (wR51-135MN) *and rad51-KN-MN* (wR51-305MN) cells incubated with and without 0.05% MMS for 2 h. Total DNA from cells permeabilized and treated or not with Ca^2+^ for 15 minutes is shown, as well as the DNA digestion profiles. ChEC experiments were repeated three times with similar results.

The pronounced MMS sensitivity (**Fig. 1, S1**) and a severe defect in SCJs formation (**Fig. 6A**) of *rad51-ED* cells contrasts with its wild type behavior in direct genetic assays of HR (**Fig. 2**). A mechanistic difference between the roles of Rad51 in HR at DSBs and MMS-induced ssDNA is that the role of Rad51 at ssDNA lesions is coupled to replication (Gonzalez-Prieto et al., 2013), whereas the role of Rad51 at DSBs is replication-independent (Alabert et al., 2009; Barlow and Rothstein, 2009). Thus, Rad51-ED might be specifically affected in the binding to MMS-induced lesions during S-phase. To address this, we constructed chimeras of Rad51 with the MNaseI (Rad51-MN) and followed their binding to DNA lesions by Chromatin Endogenous Cleavage (ChEC) analysis. In this approach, the induction of the nuclease activity of the chimera by treating cells with Ca^2+^ will only generate a detectable cut in the DNA if the repair protein is targeted to a lesion that is not a DSB. Thus, we can infer Rad51 binding to DNA if DNA is digested upon Ca^2+^ treatment (Gonzalez-Prieto et al., 2013). In the absence of MMS, most DNA remained as a single, high molecular DNA band on top of the gel; in the presence of MMS both Rad51-MN and Rad51-KN-MN generated a smear below the top band, whereas Rad51-ED-MN hardly digested the top band (**Fig. 6B**). These results suggest that Rad51-ED is severely impaired in binding to MMS-stalled replication forks or that its DNA binding substrate is not formed or degraded.

### Rad51-ED and Rad51-KN mutant proteins show altered DNA binding and ATPase profiles but are proficient in D-loop formation which depends on Rad54

The genetic analysis suggests that the *rad51-ED* and *rad51-KN* mutants separate Rad51 function in HR from that at stalled forks. To elucidate the mechanisms involved, we purified the native mutant proteins to apparent homogeneity for biochemical analysis (**Fig. S3**). The mutant proteins purified like the wild type protein and were free of contaminating activities (endo/exonuclease, topoisomerase, phosphatase) that would interfere with the *in vitro* experiments and their interpretation. The key biochemical activities of Rad51 are DNA binding, DNA-stimulated ATP hydrolysis, homology search and DNA strand invasion, which underlie its functions in HR and at stalled forks. First, we measured the ATPase activity of the mutant proteins along with wild type Rad51 and found that the ATPase activity was diminished with both ssDNA and dsDNA co-factors. The defect of the Rad51-ED mutant protein was significantly more pronounced than that of the Rad51-KN mutant (**Fig. 7A, B**).

**Figure 7:**
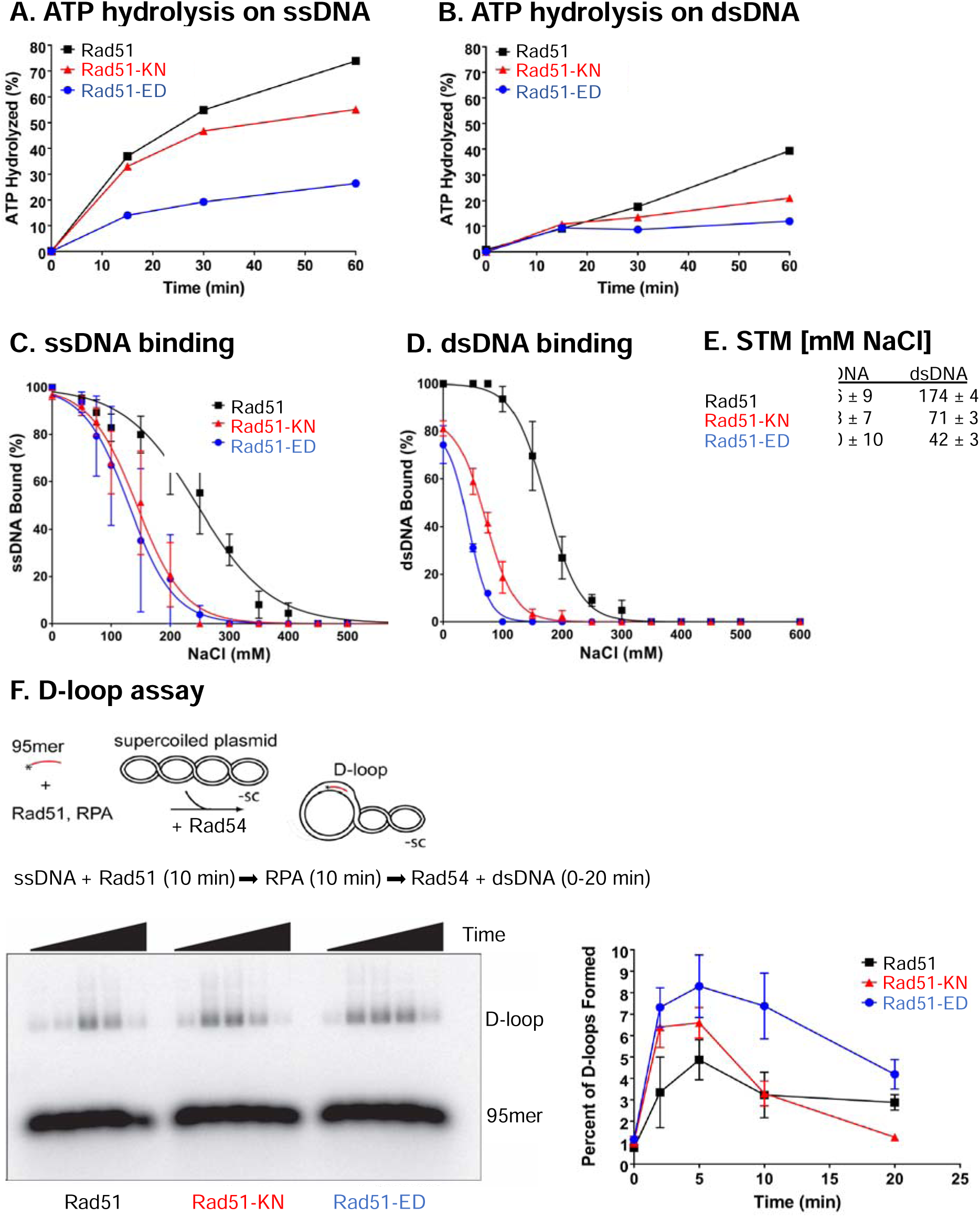
Rad51-ED and Rad51-KN mutant proteins show reduced ATPase activity, impaired DNA binding but are proficient in D-loop formation. **A, B.** ATPase rates were determined using 3 μM of the purified proteins wild type Rad51 (black rectangles), Rad51-ED (blue circles) and Rad51-KN (red triangles) in the presence of ssDNA (9 μM ΦX174 virion DNA in A) or dsDNA (9 μM ΦX174 RF1 in B). Shown are means of a minimum of n=3, error bars represent the standard error but are not visible as they were smaller than the plotting symbol. **C-E.** Salt titration midpoint (STM) DNA binding analysis by EMSA. The nucleoprotein complexes were assembled by incubating 3.33 μM of Rad51 proteins (Rad51, Rad51-ED, Rad51-KN) together with 10 μM nt of ssDNA (ΦX174 virion) or 10 µM of dsDNA ((ΦX174 RF1) in the presence of the indicated NaCl concentration for 15 min at 30 °C. All measurements were done a minimum of three times and the error bars represent the standard error. The NaCl concentrations of the calculated STM are shown in E. See **Figure S4** for representative gels. **F.** D-loop assays with 5’-end labelled 95mer (olWDH566) and pUC19 dsDNA conducted as time courses (0, 2, 5, 10, 20 min). Products were analyzed by gel electrophoresis and a representative gel is shown. Quantitation of D-loops from n=3, shown are means with error bars that represent the standard error.

The ATPase of Rad51 is strongly dependent on DNA binding, and an ATPase defect could be a reflection of an underlying DNA binding defect. As both mutants are HR proficient *in vivo* (**Fig. 2**), we expected a relatively subtle defect, as HR is dependent on Rad51 forming a filament on ssDNA. We assessed Rad51 DNA binding to ssDNA (ΦX174 viral DNA) and dsDNA (ΦX174 RF1 dsDNA) by electric mobility shift assays (EMSA) conducted as salt midpoint titrations to elucidate subtle DNA binding defects (see **Fig. S4** for representative EMSA gels). The salt titration curves (**Fig. 7C, D**) allowed the calculation of salt titration midpoints (STM; **Fig. 7E**) as the salt concentration, where 50% of the protein-DNA complexes are formed. A lower STM signals lower affinity for DNA. Under no salt conditions (0 mM NaCl), both mutant proteins showed no defect in binding to ssDNA and a small but significant defect for dsDNA (**Fig. 7C, D**). Upon increasing salt concentration, both mutant proteins showed a significant decrease in their STMs compared to wild type Rad51 for ssDNA binding. Importantly, both mutants showed a even more significant decrease in STM for dsDNA. The defect in Rad51-ED was significantly stronger than with Rad51-KN for both substrates. These results show that both mutants have lower affinity to DNA, and that the defect for dsDNA is stronger than the one for ssDNA. In fact, at physiological ionic strength (∼120 mM NaCl (Ke et al., 2013)), dsDNA binding was largely suppressed, especially by Rad51-ED. In contrast, the binding to ssDNA was reduced by only ∼30%. The severity of the dsDNA binding defect appears to correlate with the severity of the *in vivo* phenotypes with Rad51-ED showing the stronger defect *in vivo* and *in vitro* than Rad51-KN. This suggests that dsDNA binding might be a critical property of Rad51 for its function at stalled replication forks.

The *in vitro* recombination activity of the mutant proteins was assessed in the classic D-loop assay using an invading 95mer oligonucleotide and a 2,686 bp negatively supercoiled duplex target DNA in a reconstituted reaction with Rad51, Rad54 and RPA conducted as a time course (**Fig. 7F**). Rad51 wild type protein showed the expected behavior of forming D-loops peaking at 5 min that were processed over time (**Fig. 7F**). The mutant Rad51-ED and Rad51-KN proteins were at least as effective as wild type Rad51 in forming D-loop *in vitro* and the D-loops formed showed the same D-loop instability over time as in reactions with wild type Rad51 (**Fig. 7F**). D-loop formation by the Rad51-ED and Rad51-KN proteins was fully dependent on Rad54, identical to wild type Rad51 (**Fig. S5**), as expected from previous experiments. We conclude that the Rad51-ED and Rad51-KN mutant proteins are proficient in D-loop formation, consistent with the *in vivo* HR data (**Fig. 2**). Rad51-ED showed slightly more efficient D-loop formation than wild type Rad51, which could be the consequence of its lower dsDNA binding activity relieving the inhibitory effect in D-loop formation stemming from Rad51 binding to the donor. The *in vivo* and *in vitro* experiments collectively suggest that the moderate DNA binding and ATPase defects of the two mutants do not result in a measurable defect in HR. Rather, the results suggested that the mutants are defective in a process that requires a stable binding of RAD51 to DNA, and particularly to duplex DNA.

### Rad51-ED and Rad51-KN fail to protect dsDNA from degradation by Exo 1 or Dna2

Besides its key function in HR by forming the presynaptic filament on ssDNA for homology search and DNA strand invasion, Rad51 protects nascent DNA at stalled DNA replication forks from unscheduled degradation by nucleases including Exo1 and Dna2. The fork protection function of certain HR proteins was first noticed in human cells, particularly in BRCA1/2-deficient backgrounds (Berti et al., 2020; Hashimoto et al., 2010; Schlacher et al., 2011). The DNA protection function of RAD51 is distinct from its role in HR, as indicated by the absence of a requirement for RAD54 (Schlacher et al., 2011). However, the fork protection function of RAD51 involves its binding to DNA, particularly in its double-stranded form (Halder et al., 2022).

To define the capacity of the yeast Rad51 variants in DNA protection, we employed *in vitro* assays with 100 bp-long dsDNA (**Fig. 8A**). We first used yeast Exo1, one of the nucleases involved in long-range DNA end resection. Without Rad51, Exo1 degraded ∼90% of the DNA substrate. Wild type Rad51 inhibited DNA degradation by Rad51, while the two Rad51 mutants, Rad51-KN and Rad51-ED, did not protect DNA (**Fig. 8B, C**). Similar results were obtained when we used the Sgs1-Dna2 helicase-nuclease pair. While wild type Rad51 efficiently prevented DNA degradation, Rad51-KN and Rad51-ED were much less effective (**Fig. 8C, D**). Mechanistically, Rad51 prevents DNA unwinding by Sgs1, which prevents the generation of ssDNA for the Dna2 nuclease (**Fig. S6**). Rad51-KN and Rad51-ED did not prevent unwinding (**Fig. S6B, C**). Together, our reconstitution experiments revealed that Rad51-KN and Rad51-ED are impaired in DNA protection.

**Figure 8:**
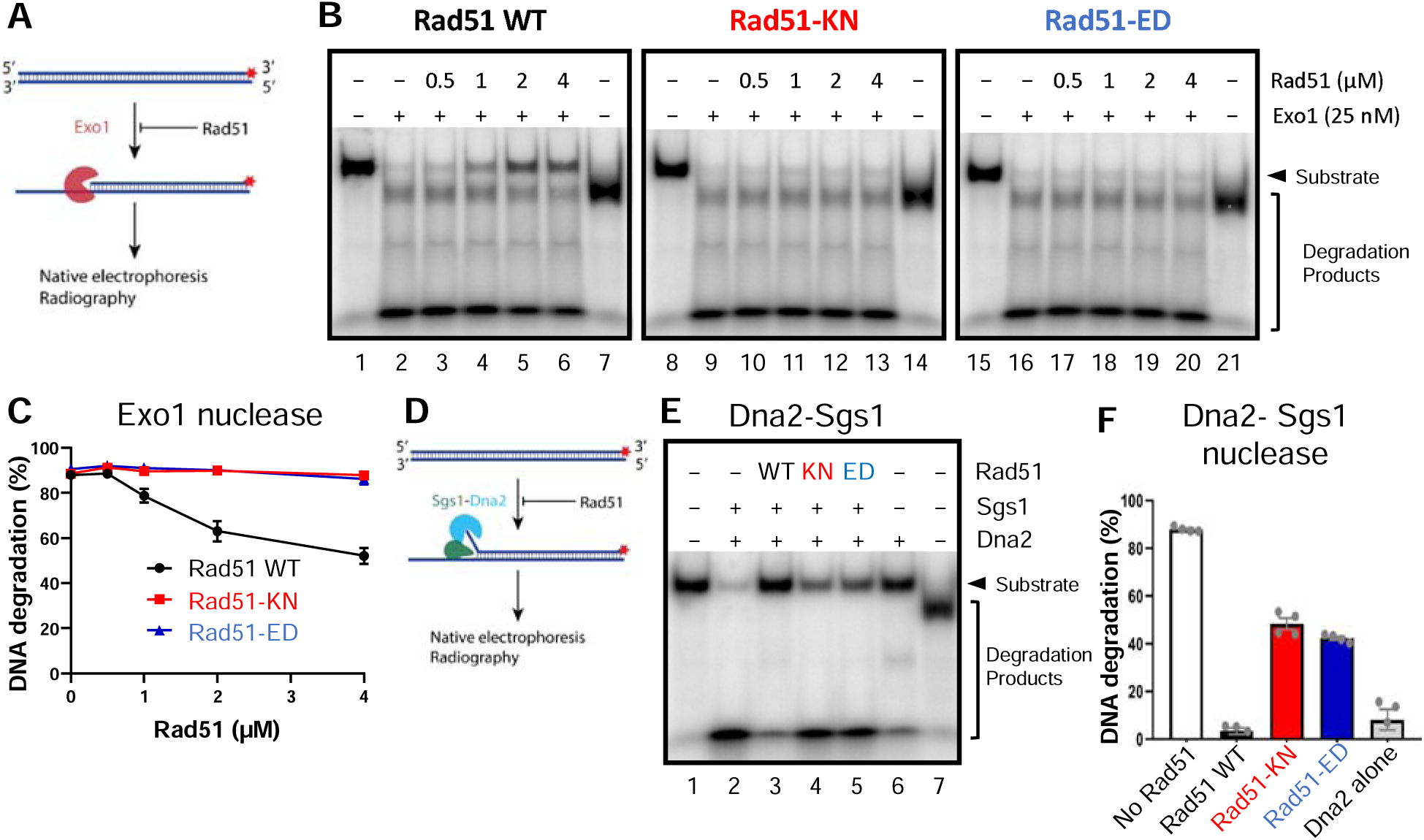
Rad51-KN and Rad51-ED are defective in protecting dsDNA from degradation by Exo1 or Dna2-Sgs1. **A.** Cartoon depicting the nuclease assay with Exo1 performed in panels B-C. The red asterisks represent the position of the radioactive label. **B.** Representative nuclease assays of Exo1 using a 100 bp DNA substrate in the presence of increasing concentrations of Rad51 wild type and variants, as indicated. In lanes 7, 14 and 21, the substrate was boiled to indicate the position of the ssDNA. **C.** Quantitation of experiments such as shown in B. Averages shown; error bars, SEM; n=3. **D.** Cartoon depicting the nuclease assay with Sgs1-Dna2 performed in panels E-F. The red asterisks represent the position of the radioactive label. **E.** Representative nuclease assay with Dna2-Sgs1 in the presence of 4 µM Rad51 wild type and variants. In lane 7, the substrate was boiled to indicate the position of the ssDNA. **F.** Quantitation of experiments such as shown in E. Averages shown; error bars, SEM; n=4.

### Molecular modeling of the Rad51-ED and Rad51-KN mutants on an experimentally derived high-resolution cryo-electron microscopic structure of the Rad51-ssDNA filament

Rad51-E135 is an invariable residue in a highly conserved region of the eukaryotic Rad51 proteins which is not conserved in the bacterial RecA homologues. Rad51-K305 is conserved in most Rad51 proteins in its charge (K or R), although some Rad51 proteins carry hydrophobic amino acids (L, A) at this position (**Fig. 9A**). To better understand the mechanistic basis of the defect of the Rad51-ED and Rad51-KN mutant proteins we modeled their structure bound to DNA based on a newly established high-resolution cryo-electron microscopic (cryo-EM) structure of the budding yeast wild type Rad51-ssDNA filament (Liu et al., 2024). In **Figure 9B**, we present the modeling of the Rad51-ED and Rad51-KN mutants on the overall Rad51-DNA filament based on the experimentally obtained structure.

**Figure 9.**
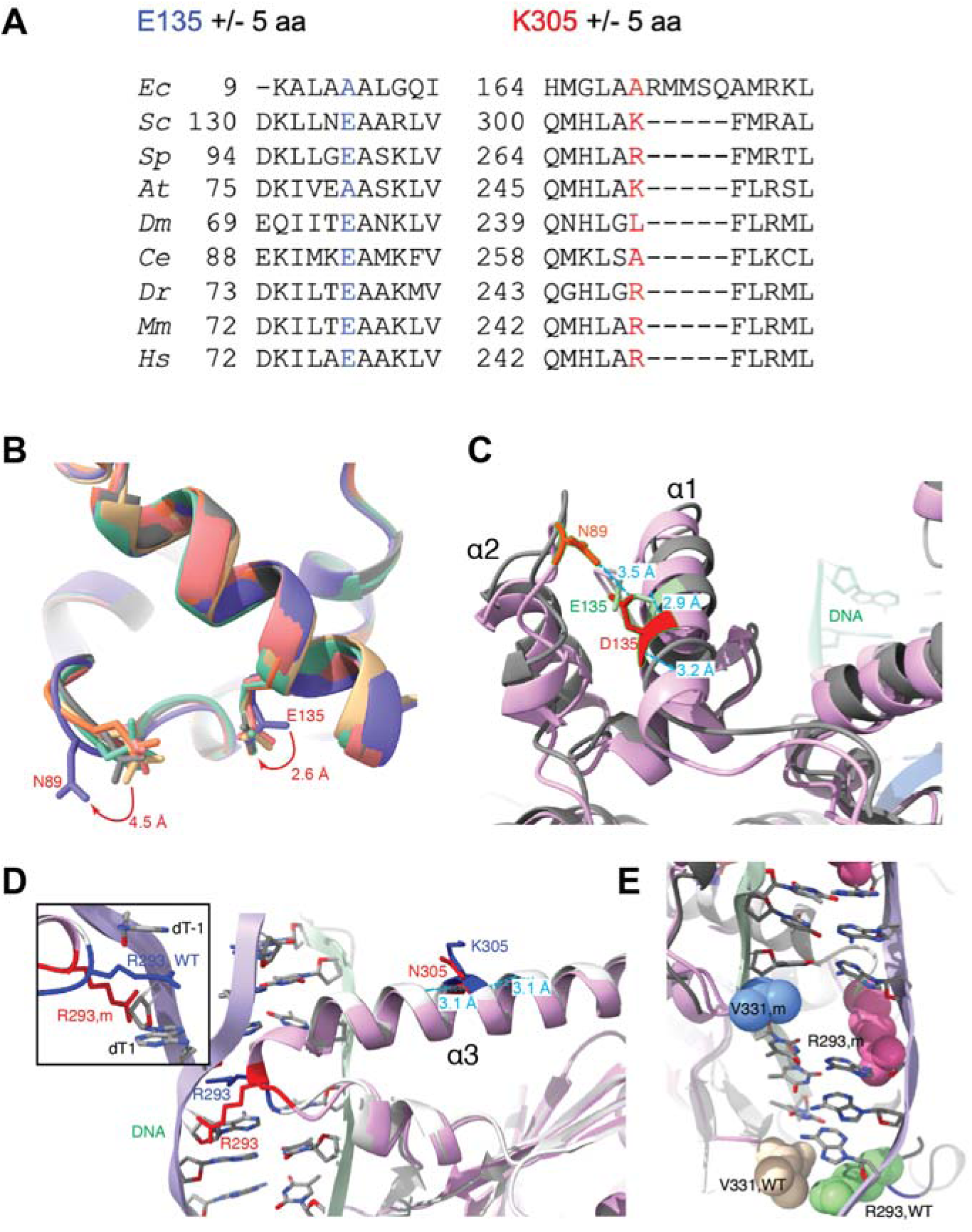
Modeling of Rad51-ED and Rad51-KN on an experimentally derived high-resolution structure of the Rad51-ssDNA filament. **A.** Sequence conservation of Rad51-ED and RAD51-KN in comparison with *Escherichia coli* RecA (*Ec*) and Rad51 from *Saccharomyces cerevisiae* (*Sc*), *Schizosaccharomyces pombe* (*Sp*), *Arabidopsis Thaliana* (*At*), *Drosophila melanogaster* (*Dm*), *Caenorhabditis elegans* (*Ce*), *Dario rerio* (*Dr*), *Mus musculus* (*Mm*), and *Homo sapiens* (*Hs*). **B.** Dynamic movements of E135 and N89 in the Rad51-ssDNA cryo-EM structure (PDB: 9B2D). **C**. Comparison of AlphaFold-predicted Rad51-ED structure (purple) with cryo-EM structure of Rad51 filament with modeled dsDNA (gray). The H-bonds formed with D135 (red) mutation were labeled with distances. N89 was labeled in orange and wild type E135 in green. **D**. Comparison of AlphaFold-predicted Rad51-KN structure (purple) with cryo-EM structure of Rad51 filament with modeled dsDNA (gray). The H-bonds formed with wild type K305 (blue) was labeled in cyan. Mutant D135 (red) shows a different orientation of R293 (red) against wild type R293 (blue). **E**. Altered contacts of R293 with V331 and dsDNA in wild type and Rad51-ED from **D**.

E135 is located in the N-terminal top region packed with α-helices, away from dsDNA and the bound ADP-AlF_3_, while K305 resides in the middle of a long α-helix which is in contact with DNA (**Fig. S7A**, **Suppl. Movie 1**). Recently, we developed a method to track the local dynamic amino acids by detecting the movement of amino acid side chains in the Rad51-ssDNA structure (Liu et al., 2024). As shown in **Figure 9B**, the side chain of E135 is dynamic and could move 2.6 Å within the two farthest conformations. We used AlphaFold to predict the structure of Rad51-E135D mutant and then compared it with the wild type Rad51 structure containing modeled dsDNA. Interestingly, in the mutant D135 has gained a new H-bond with N89 while maintaining the original two H-bonds with K131 and R138. The new H-bond between D135 and N89 pulled the α2 helix towards the α1 helix, freezing the movement of both D135 and N89 and locking the two helices together (**Fig. 9C, S7B**). Simultaneously, the flexible movement of N89, 4.5 Å within the two farthest conformations, has also been blocked (**Fig. 9B**). We predict that the gained rigidity of the Rad51-ED mutant at this flexible N-terminus would affect its ability to interact with its neighboring protomer in the filament to weaken DNA-binding cooperativity and potentially with other protein partners. K305 has limited movement within the α-helix but R283 at the end loop of this helix is very flexible with direct contacts with DNA (**Fig. S7C**). The Rad51-KN mutant has strengthened the interaction with M301 and A309, shortening the H-bond length from 3.1 Å to 3.0 Å, and 3.1 Å to 2.9 Å, respectively (**Fig. 9D**). This increased rigidity is transferred to the end of helix and limited R293 to a position to avoid collision with DNA and to lose contacts with V331 (**Fig. 9E)**. The K305N mutant is expected to have local impact on DNA-binding and maintain normal self- and hetero-protein interactions.

### Rad51-ED and RAD51-KN do not affect the interaction with the Rad55-Rad57 paralog complex

The Rad51 paralog complex Rad55-Rad57 functions in the nucleation of Rad51 filaments on RPA-coated ssDNA (Sung, 1997). It also stabilizes the Rad51-ssDNA filament against dissociation by the Srs2 helicase that has the ability to strip Rad51 from ssDNA, therefore acting as an anti-recombinase (Krejci et al., 2003; Liu et al., 2011a; Veaute et al., 2003). Detailed genetic analysis showed a strong similarity between the phenotypes of cells lacking Rad55 and the *rad51-ED* or *rad51-KN* mutants described here in their dependence on the TLS polymerase Rev3 (Maloisel et al., 2023). This opened the possible interpretation that the *rad51-ED* and *rad51-KN* mutants affected the interaction of Rad51 with the Rad55-Rad57 complex. Using purified wild-type Rad51 as well as Rad51-ED and RAD51-KN proteins in pull-down experiments with the purified Rad55-Rad57 complex, we could demonstrate that both mutants interacted normally with the Rad51 paralog complex (**Fig. S8**). We conclude that the phenotype of the *rad51-ED* and *rad51-KN* mutants is not caused by a defect in interacting with Rad55-Rad57.

## Discussion

We have isolated separation-of-function mutants in *RAD51, rad51-ED and rad51-KN*, that maintain the ability to perform HR *in vivo* and *in vitro* but severely shift the balance in the usage of post-replication repair pathways to TLS and potentially fork regression leading to a strong mutator phenotype (**Fig. S9A**). The mutations cause an impairment in the Rad51 ATPase and DNA binding activities (**Fig. 7**). The structural basis for these changes was analyzed by modeling of the mutants on an experimental high-resolution cryo-EM structure of the Rad51 filament showing that the mutants render Rad51 less dynamic and more rigid (**Fig.9**). These changes are relatively subtle and allow normal HR in cells and no significant defect in reconstituted *in vitro* D-loop reactions (**Figs. 2, 7)**. However, these changes lead to a defect in targeting Rad51 to stalled replication forks *in vivo* and in protecting dsDNA from exonucleases *in vitro* (**Figs. 6, 8**). Direct analysis of post-replication pathway usage in a competitive system with readout for TLS and TS by HR, independently demonstrated the shift of post-replication repair pathway usage (**Fig. 5**), consistent with the increased mutagenesis (**Fig. 3**) and genetic interactions with genes involved in postreplication repair (**Fig. 4**). We conclude that properties of Rad51 affect post-replication repair independent of its role in HR. This conclusion is further supported by previous studies showing that cells lacking Rad55 show elevated mutation rates and strong dependence on alternative post-replication repair pathway controlled by *REV3* and *MMS2* (Ball et al., 2009; Maloisel et al., 2023; Rattray et al., 2002; Xu et al., 2013), similar to the Rad51 mutants isolated here. The Rad55-Rad57 complex stabilizes Rad51 filaments (Liu et al., 2011a), but the mechanisms involved are likely different as the Rad51 mutants showed no defect in Rad55 interaction. In sum, the combined data show that Rad51 filament properties affect pathway usage in post-replication repair.

### What properties of Rad51 determine the phenotype of the ED and KN mutants?

The key difference emanating from the biochemical analysis was the defect of the mutants to bind dsDNA. Both mutants were significantly affected in binding dsDNA, notably more than for ssDNA (**Fig. 7**, **S4**). The much stronger defect of the Rad51-ED mutant protein than the Rad51-KN mutant in binding to dsDNA correlated with their *in vivo* phenotypes, where the *rad51-ED* mutant consistently showed a stronger phenotype than the *rad51-KN* mutant. Hence, we suggest that the relevant determinant of the ED and KN mutant phenotypes is the deficit in binding dsDNA. This is consistent with the fact that the mutant proteins are HR proficient *in vivo* and *in vitro*, as for HR, the ability of Rad51 to form filaments on ssDNA is the most critical function, whereas dsDNA binding is not required if not even detrimental (see Introduction). The ability to bind dsDNA with relatively high affinity is a newly acquired property in eukaryotic Rad51 proteins lacking in bacterial RecA (Bianco et al., 1998; Chi et al., 2006; Kowalczykowski, 1991; Tombline et al., 2002; Zaitseva et al., 1999). These observations suggest that the ability of Rad51 to bind dsDNA is unrelated to its HR function. Recent biochemical evidence with human RAD51 supports the model that dsDNA binding by RAD51 is critical for its second function in fork protection against nucleolytic degradation (Halder et al., 2022). Previous studies with a RAD51 mutant, RAD51-II3A, that is deficient in DNA strand exchange but proficient in DNA binding, showed that it retained its function in fork protection (Mason et al., 2019). RAD51-II3A represents the mirror-image separation of function mutant to *rad51-ED* and *rad51-KN*, showing that separation of function can be achieved in both directions.

The modeling shows that both mutants have increased structural rigidity and lost some conformational dynamics on dsDNA with bound nucleotides. The local structural dynamics of Rad51 protomers was illuminated in a high-resolution cryo-EM structure of the Rad51-ssDNA filament (Liu et al., 2024). Indeed, both Rad51-I345T and Rad51-H352Y display increased stability of the Rad51-ssDNA complex and exist as super-extended helical filaments. They show a shift of the N-terminal helical top when the structural alterations are transduced from the bottom ATPase core domain, compared to wild type Rad51 (Liu et al., 2024). A similar shift has also been observed for archaeal RadA, changing from the compressed to the extended filament on dsDNA (Galkin et al., 2006). The Rad51-ED mutant formed new H-bonds to lock the N-terminal helical top, impairing the transition between “active” extended and “inactive” compressed filaments. This explains the more severe phenotype of *rad51-ED*, as the increased rigidity of the Rad51-KN mutant filament only affects Rad51-KN DNA-binding but not its interaction with the neighboring Rad51 protomers, correlating with a less severe phenotype.

### Where is RAD51 binding dsDNA at stalled forks?

Fork stalling in budding yeast primarily results in non-regressed forks; in fact, extensive fork regression was only observed in cells defective for DNA damage checkpoint (Sogo et al., 2002). This contrasts with the situation in mammalian cells, where fork regression appears to be a primary response to fork stalling (Zellweger et al., 2015). Stalled forks have several ssDNA-dsDNA transitions that could be sensitive to nuclease degradation (**Fig. S9B** left). In addition, a recent study proposed a function of RAD51 in fork stabilization by binding to dsDNA to lock in the CMG helicase and prevent loss of the replicative helicase to enable direct restart (Liu et al., 2023). Interestingly, Rad51 was found to physically interact with the MCM complex (Cabello-Lobato et al., 2021), providing a potential mean to nucleate Rad51 filaments at stalled forks (**Fig. S9B**). It is unclear whether each and any of these potential dsDNA binding sites of RAD51 is relevant for fork protection, but the multitude of nucleases involved, including EXO1, DNA2, MRE11, and potentially others (Berti et al., 2020), may suggest that several sites may be relevant depending on context. We speculate that in the rad51-ED and rad51-KN mutants, DNA degradation would convert internal gaps (S9B, left) and regressed forks (S9B, right) into gapped forks that would be extended by TLS to restart replication.

Our results also provide indirect evidence for an HR (Rad51)-independent template switching mechanism in budding yeast that depends on PCNA-poly ubiquitylation (Mms2), which we suspect is fork regression. First, template switching is not eliminated in *rad51Δ* cells (**Fig. 5**). This suggests the existence of a Rad51-independent template switch mechanism, which could be fork regression driven by motor proteins such as Rad5. Second, *rad51Δ* cells show additive MMS-sensitivity with Mms2 (**Fig. 4 C-E**). Mms2 functions through K63 poly-ubiquitylation of PCNA on K164 (Hoege et al., 2002; Hofmann and Pickart, 1999). This result suggests that Mms2 controls an HR-independent template switch process in postreplication repair, likely fork regression as gap repair by TS is Rad51 dependent. Third, in *ubc13* cells where PCNA polyubiquitination is prevented, template switching is only partially eliminated as in *rad51Δ* cells (Maslowska et al., 2019; Maslowska and Pagès, 2023) (**Fig. 5**). Together these data strongly suggest the existence of two template switching pathways, one Rad51-dependent (gap repair) and one Rad51-independent (likely fork regression) (**Fig.S9A**). Fourth, in checkpoint-deficient yeast cells (*rad53*) regressed forks readily form (Sogo et al., 2002) and likely present a target for dsDNA-specific exonucleases like Exo1 with likely deleterious consequences (**Fig. S9B** right). Rad51 binding to dsDNA in these regions is expected to suppress nucleolytic attack. In mammalian cells, motor proteins implicated in fork regression (SMARCAL1, ZRANB3, and HLTF) facilitate DNA degradation, suggesting that regressed forks are substrates for DNA degradation (Kolinjivadi et al., 2017; Mijic et al., 2017; Taglialatela et al., 2017). In fact, it was found that eliminating Exo1 effectively suppresses the MMS sensitivity of yeast *rad53* mutants (Segurado and Diffley, 2008). While regressed forks might be rare or highly unstable in wild type cells, there are conditions, such as in checkpoint defective cells, where fork regression is readily documented in budding yeast (Sogo et al., 2002). This implies that the mechanisms for fork regression exist in budding yeast but are under negative control. Hence, Rad51 may also be binding to regressed forks to protect against dsDNA nucleases (**Fig. S9B** right).

### Why are Rad51-ED and Rad51-KN proficient for HR and defective in template switching by HR at stalled forks?

The *rad51-ED* and *rad51-KN* mutants are clearly HR-proficient *in vivo* and *in vitro* (**Fig. 2, 7**), while they are defective in generating HR-intermediates at stalled replication forks (**Fig. 6**). We suspect that the difference between both scenarios is the loading of Rad51, as especially Rad51-ED shows a defect in binding to stalled forks (**Fig. 6B**). While in recombination Rad51 is nucleated on ssDNA, we speculate that Rad51 is nucleated at stalled forks on duplex DNA or junction structures, possibly through its interaction with the replicative helicase (Cabello-Lobato et al., 2021). This speculation is consistent with models for RAD51 binding at stalled replication forks in mammalian cells to protect against exonucleolytic degradation (Halder et al., 2022) and locking in the CMG helicase (Liu et al., 2023) to allow direct restart. Such binding of Rad51 to dsDNA would be resistant against disruption by Srs2, as Srs2 only disrupts Rad51-ssDNA filaments, in addition to the local inhibition of Srs2 at stalled forks (Krejci et al., 2003; Urulangodi et al., 2015). Alternatively, it is possible that the Rad51-ED/KN loading site at stalled forks is degraded.

In sum, the ability of Rad51 to bind dsDNA with relatively high affinity appears to correlate with its acquired function to protect stalled replication forks against nucleolytic degradation and potentially retain the replicative helicase for direct restart. The genetic separation of the Rad51 in HR and fork protection in both directions (Rad51-ED/KN HR + fork protection -; RAD51-II3A HR - fork protection +) suggests that it might be possible to develop targeted inhibitors for each function.

## MATERIALS AND METHODS

### Yeast strains and growth conditions

Standard techniques for yeast growth and genetic manipulation were used (Burke et al., 2000). All yeast strains used for genetic experiments are in the W303-1A background and WT *RAD5* (**Table S1**), except for some strains used in **Figure S1C** which are *rad5-G535R* as noted. See **Table S2** for all plasmids used. See **Table S3** for all oligonucleotides and their sequences.

### Rad51 mutant isolation and genomic integration

To screen for *rad51* separation-of-function mutants, a random pool was created by mutagenic PCR using the GeneMorph II EZClone Domain Mutagenesis Kit that employs two error prone polymerases to achieve balanced mutations in all four bases. The *RAD51* open reading frame is 1.2 kb in length; the ideal pool of mutants would have one to a few mutations per gene. Mutagenic PCR conditions were empirically optimized to obtain a mutation rate of ∼1 mutation per kb, by varying the amount of template DNA and reducing the number of PCR cycles using olWDH632 and olWDH633 which are 400 nt upstream and downstream of the *RAD51* open reading frame, respectively, and plasmid pWDH957, which contains the wild type *RAD51* gene with 1,000 bp upstream and downstream of the *RAD51* open reading frame cloned as a 3.2 kb *Hind*III-*Sac*I fragment into Yep351 (pWDH958) using olWDH830 and olWDH1355. The pool used averaged two mutations per gene. The screen was conducted in a *rad51*Δ *rad54^ts^* strain (WDHY2546) to take advantage of the conditional phenotype. The *rad54^ts^* allele was determined to be caused by a single amino acid change (C692Y), which leads to loss of protein at the restrictive temperature. The mutagenized *RAD51* pool was co-transformed with pWDH957 digested with *Stu*I and *Nru*I to cut out almost the entire *RAD51* ORF from −22 to 47 bp upstream the stop codon for *in vivo* recombination to form functional circular YEp351 plasmids containing a mutagenized *RAD51* gene. Approximately 2,000 transformants were screened. Of these, 40 candidates exhibited improved viability when challenged with MMS compared to the *rad51*Δ *rad54^ts^* strain at 37° C. The plasmid dependence of the phenotype was confirmed. Plasmids were recovered in *E. coli,* the *RAD51* sequence was established, and the phenotype re-confirmed by transformation of a host strain that has never been exposed to MMS.

The mutations *rad51-ED* and *rad51-KN* were transferred to the native *RAD51* locus of WDHY2217 that contained a complete deletion of the *URA3* gene to preclude reversion events using the PCR-based allele replacement method of Erdeniz et al. (Erdeniz et al., 1997) with the *RAD51* primers olWDH1351 and 1353 as well as the *K. lactis URA3* primers olWDH1357 and 1358. Correct integration and the integrity of the integrated sequences were confirmed by amplifying the *RAD51* locus and sequencing the entire gene on both strands.

### Serial dilution assays

The serial dilution spot assays were performed on solid YPD or YPD with the addition of MMS at the dilutions specified. To assay for sensitivity to ionizing radiation (IR), YPD plates were spotted with cells and allowed to dry before being exposed to IR (Cesium-137 source). Cells were spotted at 5-fold dilutions starting at approximately 2 x 10^7^ cells/ml (OD600=1). The plates were incubated at 30 °C and photographed daily for up to 4 days.

### Immunoblots

Lysates were prepared by trichloroacetic acid (TCA) precipitations from cells grown to early log phase, washed, and re-suspended in 20 % TCA before cell disruption utilizing a FastPrep machine (FP120, Bio101, Savant) on setting “4”, three times for 15 seconds. Lysates were transferred to a new 1.5 mL tube and beads were washed three times with 5 % TCA. Precipitated protein was collected at 6,000 RPM in an Eppendorf centrifuge for 10 minutes, resuspended in 1.5 x Laemmli sample buffer (Laemmli, 1970), and neutralized with one-third volume of Tris-base, before analysis by gel electrophoresis. Immunoblots were conducted as described (Bashkirov et al., 2000). Lysate from 2.8 OD600 units of cells were loaded and separated on a 10 % SDS-PAGE gel before transfer to nitrocellulose. The membrane was blocked in 5 % skim milk dissolved in TBS (10 % Tris pH 7.5, 150 mM NaCl) overnight, the primary rabbit antibody (a generous gift from Dr. Akira Shinohara) was diluted 1:2,500 in 2 % skim milk and incubated for 6 hrs. Primary antibody was followed by five, 5 min washes in 2 % skim milk dissolved in TBS. Horse radish peroxidase (HRP)-conjugated anti-rabbit secondary antibody was applied (1:5,000) in 2 % skim milk in TBS for one hr at room temperature and subsequently washed three times (5 min each) with 2 % milk in TBS, followed by two, 5 min washes with TBS. HRP was visualized with Perkin Elmer Western Lightening Plus (Perkin Elmer, NEL103E001EA) and GE Life sciences hyperfilm (GE Healthcare and Life Sciences, 28-9068-38). All membranes were stripped with stripping buffer (62.5 mM Tris-HCl pH 6.7, 0.704 % β-mercaptoethanol, 2 % SDS) for 30 minutes at 50 °C prior to reprobing using mouse anti-3-phosphoglycerate kinase anti-serum (Invitrogen-Molecular probes, A6457).

### Direct-Repeat Sister-Chromatid Recombination Assay

Single colonies were grown in synthetic complete (SC) media lacking uracil with raffinose as a carbon source for 24 hrs at 30°C. To determine spontaneous recombination rates, appropriate dilutions were plated on YPD media to determine viability, and on SC-URA, and SC-URA-LEU to measure recombinants. 2 % galactose was added to the cultures to induce expression of the HO endonuclease, which was expressed for 2 hrs before the cells were collected and plated to determine DSB-induced recombination frequencies. After 3 days of growth at 30 °C, colonies were counted from at least 12 independent cultures per genotype by the method of the median (Lea and Coulson, 1949).

### Analysis of sister chromatid junctions (SCJs)

Yeast cells were grown at 30°C in a supplemented minimal medium (SMM). For G1 synchronization, cells were grown to mid-log phase and α-factor was added twice at 60 min intervals at 2 μg/ml (*BAR1* strains). Then, cells were washed three times and released into fresh medium with 50 μg/ml pronase in the absence or presence of MMS at the indicated concentrations. SCJs (X-shaped molecules) were analyzed by 2D-gel electrophoresis from cells arrested with sodium azide (0.1% final concentration) and cooled down on ice as reported (Clemente-Ruiz and Prado, 2009). Briefly, total DNA was isolated with the G2/CTAB protocol, digested with *Eco*RV and *Hind*III, resolved by neutral/neutral two-dimensional gel electrophoresis, blotted to Hybond^TM^-XL membranes, and analyzed by hybridization with the ^32^P-labelled A probe. Signals were quantified in a Fuji FLA5100 with the ImageGauge analysis program.

### *In vivo* ChEC analysis

Chromatin endogenous cleavage (ChEC) of *RAD51-MN*, *rad51-KN-MN* and *rad51-ED-MN* cells was performed as reported (Gonzalez-Prieto et al., 2013) from cultures grown at 30°C in SMM with or without 0.05% MMS for 2 hours and arrested with sodium azide (0.1% final concentration). For cleavage induction, digitonin-permeabilized cells were incubated with 2 mM CaCl_2_ at 30°C under gentle agitation. Total DNA was isolated and resolved into 0.8% TAE 1× agarose gels. Gels were scanned in a Fuji FLA5100, and the signal profile quantified using ImageGauge. The area of the DNA digestion profiles was equalized to eliminate DNA loading differences.

### Direct post-replication repair analysis in cells

All strains used for the iDamage assay are derivatives of the EMY74.7 strain (Johnson et al., 1998) In order to study tolerance events, all strains are deficient in repair mechanisms: nucleotide excision repair (*rad14*) and mismatch repair (*msh2*). Gene disruptions were achieved using PCR-mediated seamless gene deletion (Akada et al., 2006) or URAblaster (Alani and Kleckner, 1987) techniques. Rad51 mutations were introduced using CRISPR/Cas9 (Laughery et al., 2015). Integration of plasmids carrying the (6-4)TT and N2dG-AAF lesions (or control plasmids without lesion) was performed as previously described (Maslowska et al., 2019). Lesion tolerance rates were calculated as the relative integration efficiencies of damaged vs. non-damaged vectors normalized by the transformation efficiency of a control plasmid in the same experiment. DA events are calculated by subtracting TLS events from the total lesion tolerance events. All experiments were performed at least in triplicate. Graphs and statistical analysis were done using GraphPad Prism applying unpaired t-test. Bars represent the mean value ± s.d.

### Protein purifications

Rad51 mutant and wild type proteins were purified to homogeneity as described (New et al., 1998; Sung, 1994). The host *Saccharomyces cerevisiae* strain (WDHY1611; **Table S1**) had a chromosomal deletion of the *RAD51* gene to avoid contaminating mutant protein preparation by wild type Rad51 protein. In brief, the mutant and wild type proteins (native, untagged) were purified using a four-column purification strategy including: Q-Sepharose, Cibacron Blue, Hydroxyapatite, and Mono Q. Rad54 (Nimonkar et al., 2012) and RPA (Binz et al., 2006) were purified as described. GST-Rad55-His_6_-Rad57 were purified as described (Liu et al.,2011b) and dialyzed into storage buffer containing 20_mM Tris-HCl pH_7.5, 0.2_M NaCl, 0.1_mM EDTA, 1_mM DTT and 50% glycerol. Sgs1 was expressed in *Spodoptera frugiperda* (*Sf*9) cells using the pFB-MBP-Sgs1 vector and purified by amylose and NiNTA affinity purification as previously described (Cejka and Kowalczykowski, 2010). The N-terminal MBP tag was removed after the amylose purification step using PreScission Protease. Yeast Exo1 was purified from *Sf*9 cells with pFB-Exo1-FLAG by FLAG affinity and HiTrap SP HP (Cytiva) ion exchange chromatography as described previously (Cannavo et al., 2013). Dna2 was expressed in the *S. cerevisiae* strain WDH668 from the pGAL18 Flag-HA-Dna wt-His: vector and purified by NiNTA and FLAG affinity chromatography as described (Budd et al., 2000; Cejka et al., 2010). Yeast RPA was expressed using the p11d–tRPA vector (a kind gift from M. Wold, University of Iowa) in BL21 (DE3) pLysS cells and purified on a HiTrap Blue column (Cytiva) followed by desalting with a HiTrap Desalting column (Cytiva) and finally with a HiTrap Q column (Cytiva) (Anand et al., 2018).

### Biochemical assays

ATPase assay: The ATP analysis was carried out using Thin Layer Chromatography (TLC), as described (Carreira et al., 2009). 3 μM Rad51 was pre-incubated with 9 μM (nt or bp) DNA (phiX174 ss- or dsDNA), at 30 °C, in 20 mM Tris HCl (pH 7.5), 10 mM MgCl_2_, 0.1 mg/ml BSA, 1 mM DTT, and 200 mM NaCl, then 0.05 mM ATP (1:100 nt radioactive) was added to initiate the reaction and start the time course. One μl aliquots were then spotted onto cellulose polyethyleneimine TLC plates. The plates were air dried and developed in 1 M formic acid and 0.5 M LiCl. The amount of ATP hydrolyzed was analyzed by PhosphorImager and quantified using ImageQuant software (Molecular Dynamics, Inc., Sunnyvale CA, USA).

DNA binding assay: DNA binding assays were performed as described (Li et al., 2007) and conducted as salt titrations to assay the capability of Rad51 to bind DNA (ss- or dsDNA) in the presence of increasing concentrations of salt. DNA (10 μM nt/bp, ΦX174), salt (0-600 mM NaCl), and Rad51 (3.33 μM), were incubated for 15 min at 30 °C in the following buffer: 20 mM TEA, 1 mM DTT, 25 μg/ml BSA, 4 mM MgOAc, and 2.5 mM ATP. The reactions were stopped by the addition of glutaraldehyde (0.25 %) and incubated for an additional 15 min at 30 °C, before being analyzed on 0.8 % agarose gels in TAE buffer (40 mM Tris base, 20 mM acetic acid, 1 mM EDTA) 3.75 V/cm for 100 min. The dried gels were analyzed by PhosphoImager and quantified using ImageQuant software (Molecular Dynamics, Inc., Sunnyvale CA, USA). The data were plotted using Prism GraphPad software which also determined the salt-titration midpoint.

D-loop assay: D-loop assays were performed as described (Liu et al., 2011b). Briefly, a 95-mer primer (olWDH#566 (see Table S3), 20 nM molecules) and Rad51 (0.67 μM) were incubated for 10 min at 30 °C, in buffer (30 mM TrisOAc pH 7.5, 1 mM DTT, 50 mg/ml BSA, 5 mM MgOAc, 20 mM Phosphocreatine, 4 mM ATP, and 0.1 mg/ml creatine kinase). RPA (0.1 µM) was added and incubated for another 10 min at 30 °C, followed by addition of Rad54 (0.112 μM) and dsDNA plasmid (20 nM molecules pUC19) to initiate the reaction and start the time course at 30 °C. The amounts of D-loops formed were analyzed by PhosphorImager and quantified using ImageQuant software (Molecular Dynamics, Inc., Sunnyvale CA, USA).

Preparation of oligonucleotide-based substrates: The oligonucleotide-based DNA substrate used for the helicase and nuclease assays was generated by labeling the BIO100C oligonucleotide (see Table S1 for oligonucleotides sequence) at the 3’ end using terminal transferase (New England Biolabs) and [α-32P]dCTP (Hartmann Analytic) according to manufacturer’s instructions and purified on G25 columns (Cytiva). The purified oligonucleotide was mixed with a two-fold excess of BIO100 oligonucleotide (see Table S1 for oligonucleotides sequence) in annealing buffer (10 mM Tris-HCl pH 8, 50 mM NaCl, 10 mM MgCl2), heated to 95°C for 3 min and cooled down to room temperature overnight.

Helicase and nuclease assays: Helicase and nuclease assays were performed with a 100 bp substrate (1 nM, in molecules, oligonucleotides BIO100C and BIO100) in reaction buffer containing 25 mM Tris-acetate pH 7.5, 1 mM dithiothreitol, 2 mM magnesium acetate, 1 mM ATP, 80 U/ml pyruvate kinase (Sigma), 1 mM phosphoenolpyruvate and 0.15 mg/ml bovine serum albumin (New England Biolabs), in a final volume of 15 µl. The reactions were performed in the presence of 45 nM (final) *Saccharomyces cerevisiae* RPA. The indicated Rad51 variant was added to the reaction followed by the indicated resection factors, Sgs1 (100 nM) for the helicase assays, and Exo1 (25 nM) or Dna2-Sgs1 (1 nM each) for the nuclease assays. The reaction was incubated for 30 min at 30 °C and stopped by the addition of 5 µl of 2% STOP buffer (150 mM ethylenediaminetetraacetic acid [EDTA], 2% sodium dodecyl sulfate, 30% glycerol, bromophenol blue) supplemented with a 20-fold excess of unlabeled oligonucleotide with the same sequence as the labeled one (to prevent reannealing) and 1 µl of Proteinase K (18 mg/ml) (Roche). After deproteination for 30 min at 37 °C, the reactions were separated on a 10% TBE gel. The gel was dried on 17 Chr paper (Whatman), imaged on a Typhoon Imager (Cytiva) and quantitated using the ImageJ software.

Protein pulldown assay: 100 nM GST–Rad55–His_6_–Rad57 or GST (Pierce) were incubated with 100 nM Rad51 in buffer P containing 25_mM Tris-acetate (pH_7.5), 10_mM magnesium acetate, 50_mM NaCl, 1_mM DTT, 10% glycerol and 0.01% NP-40 in a 25 μl volume for 1_h at room temperature. Equilibrated and BSA-treated glutathione agarose beads (Pierce) were added to the mixture and incubated for 1_h. The beads and supernatant were separated by centrifugation and the beads were washed twice with binding buffer P. The pulled-down protein complexes were eluted by heating at 65_°C for 10_min in 12_μl 2x SDS–PAGE loading buffer and separated through a 10% SDS–PAGE gel. Anti-Rad51 (Santa Cruz Biotechnology) and anti-GST (Cytiva) antibodies were used for immunoblotting. Protein bands were visualized using Clarity Western ECL substrate (Bio-Rad) on immunoblots and quantified using Image Lab (Bio-Rad). 5% of the input and 100% of the elution for each reaction were loaded for comparison.

### Statistical analysis

The 95% or better confidence intervals for the median were calculated using the information provided in **Table S4.** This utilizes the formula: (n+1)/2 ± 1.96 (√n/2) to provide the upper and lower limit of the 95% or better confidence interval from the rank order of values. As such, this method will generate an uneven ± confidence interval around the median, making it easier to compare multiple bar graphs on a single figure. The p-values were always generated between two data sets using pairwise unpaired t-test with Welch’s correction that does not assume equal standard deviations between data sets.

## Acknowledgements

We thank Steve Kowalczykowski and Neil Hunter for sharing equipment, Rodney Rothstein for strains, and Akira Shinohara for anti-Rad51 serum. This work was supported by grants GM58015 and GM131037 from the US National Institutes of Health (WDH), Cancer Center Core Support Grant NCI P30CA093373 to UCD, grants BFU2015-63698-P and PID2021-127486NB-100 from MCIN/AEI/10.13039/501100011033 and “ERDF A way of making Europe” (FP), the Swiss National Science Foundation (SNSF) (Grants 310030_207588 and 310030_205199) and the European Research Council (ERC) (Grant 101018257) (PC), and Fondation pour la Recherche Médicale [FRM-EQU201903007797] (VP). DM was partially supported by the NCI T32 Oncogenic Signaling and Chromosome Biology training program T32 CA108459, SJC was partially supported by the NIH T32 Molecular & Cellular Biology training program T32 GM007377; CE was supported by a postdoctoral fellowship from the Deutsche Forschungsgemeinschaft (DFG); MIC-L was supported by a fellowship from Spanish government FPU13/00955.

## Author Contributions

DM, SJC, SG, CE, GR, MICL, KHM, FV conducted the experiments; JL and SG conducted the molecular modeling; DM, FP, PC, VP, and WDH designed experiments and interpreted results; WDH designed the study and wrote the manuscript with contributions from DM, FP, PC, VP, and JL.

## References

Aboussekhra, A., Chanet, R., Adjiri, A., and Fabre, F. (1992). Semi-dominant suppressors of Srs2 helicase mutations of *Saccharomyces cerevisiae* map in the *RAD51* gene, whose sequence predicts a protein with similarities to procaryotic RecA protein. Mol Cell Biol 12, 3224–3234.

Aboussekhra, A., Chanet, R., Zgaga, Z., Cassier Chauvat, C., Heude, M., and Fabre, F. (1989). *RADH*, a gene of *Saccharomyces cerevisiae* encoding a putative DNA helicase involved in DNA repair. Characteristics of *radH* mutants and sequence of the gene. Nucleic Acids Res 17, 7211–7219.

Akada, R., Kitagawa, T., Kaneko, S., Toyonaga, D., Ito, S., Kakihara, Y., Hoshida, H., Morimura, S., Kondo, A., and Kida, K. (2006). PCR-mediated seamless gene deletion and marker recycling in *Saccharomyces cerevisiae*. Yeast 23, 399–405.

Alabert, C., Bianco, J.N., and Pasero, P. (2009). Differential regulation of homologous recombination at DNA breaks and replication forks by the Mrc1 branch of the S-phase checkpoint. EMBO J 28, 1131–1141.

Alani, E., and Kleckner, N. (1987). A method for gene disruption that allows repeated use of URA3 selection in the construction of multiply disrupted yeast strains. Genetics 116, 541–555.

Anand, R., Pinto, C., and Cejka, P. (2018). Methods to Study DNA End Resection I: Recombinant Protein Purification. Methods Enzymol 600, 25–66.

Arbel, M., Liefshitz, B., and Kupiec, M. (2021). DNA damage bypass pathways and their effect on mutagenesis in yeast. FEMS microbiology reviews 45.

Ball, L.G., Zhang, K., Cobb, J.A., Boone, C., and Xiao, W. (2009). The yeast Shu complex couples error-free post-replication repair to homologous recombination. Mol Microbiol 73, 89–102.

Barlow, J.H., and Rothstein, R. (2009). Rad52 recruitment is DNA replication independent and regulated by Cdc28 and the Mec1 kinase. EMBO J 28, 1121–1130.

Bashkirov, V.I., King, J.S., Bashkirova, E.V., Schmuckli-Maurer, J., and Heyer, W.D. (2000). DNA repair protein Rad55 is a terminal substrate of the DNA damage checkpoints. Mol Cell Biol 20, 4393–4404.

Berti, M., Cortez, D., and Lopes, M. (2020). The plasticity of DNA replication forks in response to clinically relevant genotoxic stress. Nat Rev Mol Cell Biol.

Bianco, P.R., Tracy, R.B., and Kowalczykowski, S.C. (1998). DNA strand exchange proteins: a biochemical and physical comparison. Front Biosci 3, 570–603.

Binz, S.K., Dickson, A.M., Haring, S.J., and Wold, M.S. (2006). Functional assays for replication protein A (RPA). Methods Enzymol 409, 11–38.

Blastyak, A., Hajdu, I., Unk, I., and Haracska, L. (2010). Role of Double-Stranded DNA Translocase Activity of Human HLTF in Replication of Damaged DNA. Mol Cell Biol 30, 684–693.

Blastyak, A., Pinter, L., Unk, I., Prakash, L., Prakash, S., and Haracska, L. (2007). Yeast Rad5 protein required for postreplication repair has a DNA helicase activity specific for replication fork regression. Mol Cell 28, 167–175.

Branzei, D. (2011). Ubiquitin family modifications and template switching. FEBS Letters 585, 2810–2817.

Branzei, D., and Psakhye, I. (2016). DNA damage tolerance. Current Opinion in Cell Biology 40, 137–144.

Branzei, D., and Szakal, B. (2016). DNA damage tolerance by recombination: Molecular pathways and DNA structures. DNA Repair 44, 68–75.

Broomfield, S., Hryciw, T., and Xiao, W. (2001). DNA postreplication repair and mutagenesis in *Saccharomyces cerevisiae*. Mutat Res-DNA Repair 486, 167–184.

Budd, M.E., Choe, W., and Campbell, J.L. (2000). The nuclease activity of the yeast DNA2 protein, which is related to the RecB-like nucleases, is essential in vivo. J Biol Chem 275, 16518–16529.

Burke, D.J., Dawson, D., and Stearns, T. (2000). Methods in Yeast Genetics: A Cold Spring Harbor Laboratory Course Manual. (Cold Spring Habror Laboratory Press).

Cabello-Lobato, M.J., Gonzalez-Garrido, C., Cano-Linares, M.I., Wong, R.P., Yanez-Vilchez, A., Morillo-Huesca, M., Roldan-Romero, J.M., Vicioso, M., Gonzalez-Prieto, R., Ulrich, H.D., et al. (2021). Physical interactions between MCM and Rad51 facilitate replication fork lesion bypass and ssDNA gap filling by non-recombinogenic functions. Cell Rep 36, 109440.

Cannavo, E., Cejka, P., and Kowalczykowski, S.C. (2013). Relationship of DNA degradation by *Saccharomyces cerevisiae* Exonuclease 1 and its stimulation by RPA and Mre11-Rad50-Xrs2 to DNA end resection. Proc Natl Acad Sci USA 110, E1661–E1668.

Carr, A., and Lambert, S. (2021). Recombination-dependent replication: new perspectives from site-specific fork barriers. Current Opinion in Genetics & Development 71, 129–135.

Carreira, A., Hilario, J., Amitani, I., Baskin, R.J., Shivji, M.K.K., Venkitaraman, A.R., and Kowalczykowski, S.C. (2009). The BRC Repeats of BRCA2 Modulate the DNA-Binding Selectivity of RAD51. Cell 136, 1032–1043.

Cejka, P., Cannavo, E., Polaczek, P., Masuda-Sasa, T., Pokharel, S., Campbell, J.L., and Kowalczykowski, S.C. (2010). DNA end resection by Dna2-Sgs1-RPA and its stimulation by Top3-Rmi1 and Mre11-Rad50-Xrs2. Nature 467, 112–116.

Cejka, P., and Kowalczykowski, S.C. (2010). The full-length *Saccharomyces cerevisiae* Sgs1 protein is a vigorous DNA helicase that preferentially unwinds Holliday junctions. J Biol Chem 285, 8290–8301.

Cejka, P., and Symington, L.S. (2021). DNA End Resection: Mechanism and Control. In Annual Review of Genetics, Vol 55, N.M. Bonini, and T. Piotrowski, eds., pp. 285–307.

Chi, P., Van Komen, S., Sehorn, M.G., Sigurdsson, S., and Sung, P. (2006). Roles of ATP binding and ATP hydrolysis in human Rad51 recombinase function. DNA Repair 5, 381–391.

Ciccia, A., Nimonkar, A.V., Hu, Y.D., Hajdu, I., Achar, Y.J., Izhar, L., Petit, S.A., Adamson, B., Yoon, J.C., Kowalczykowski, S.C., et al. (2012). Polyubiquitinated PCNA Recruits the ZRANB3 Translocase to Maintain Genomic Integrity after Replication Stress. Mol Cell 47, 396–409.

Clemente-Ruiz, M., and Prado, F. (2009). Chromatin assembly controls replication fork stability. Embo Reports 10, 790–796.

Daigaku, Y., Davies, A.A., and Ulrich, H.D. (2010). Ubiquitin-dependent DNA damage bypass is separable from genome replication. Nature 465, 951–U913.

Daley, J.M., Gaines, W.A., Kwon, Y., and Sung, P. (2014). Regulation of DNA Pairing in Homologous Recombination. Cold Spring Harbor Perspectives in Biology 6.

Deem, A., Keszthelyi, A., Blackgrove, T., Vayl, A., Coffey, B., Mathur, R., Chabes, A., and Malkova, A. (2011). Break-induced replication is highly inaccurate. PLoS Biol 9, e1000594.

Erdeniz, N., Mortensen, U.H., and Rothstein, R. (1997). Cloning-free PCR-based allele replacement methods. Genome Research 7, 1174–1183.

Fan, H.Y., Cheng, K.K., and Klein, H.L. (1996). Mutations in the RNA polymerase II transcription machinery suppress the hyperrecombination mutant *hpr1 Delta* of *Saccharomyces cerevisiae*. Genetics 142, 749–759.

Fan, Q.F., Xu, X., Zhao, X., Wang, Q., Xiao, W., Guo, Y., and Fu, Y.V. (2018). Rad5 coordinates translesion DNA synthesis pathway by recognizing specific DNA structures in saccharomyces cerevisiae. Curr Genet 64, 889–899.

Galkin, V.E., Wu, Y., Zhang, X.P., Qian, X., He, Y., Yu, X., Heyer, W.D., Luo, Y., and Egelman, E.H. (2006). The Rad51/RadA N-terminal domain activates nucleoprotein filament ATPase activity. Structure 14, 983–992.

Gallo, D., Kim, T., Szakal, B., Saayman, X., Narula, A., Park, Y., Branzei, D., Zhang, Z.L., and Brown, G.W. (2019). Rad5 Recruits Error-Prone DNA Polymerases for Mutagenic Repair of ssDNA Gaps on Undamaged Templates. Mol Cell 73, 900-+.

Gangavarapu, V., Haracska, L., Unk, I., Johnson, R.E., Prakash, S., and Prakash, L. (2006). Mms2-Ubc13-dependent and -independent roles of Rad5 ubiquitin ligase in postreplication repair and translesion DNA synthesis in Saccharomyces cerevisiae. Mol Cell Biol 26, 7783–7790.

Giannattasio, M., Zwicky, K., Follonier, C., Foiani, M., Lopes, M., and Branzei, D. (2014). Visualization of recombination-mediated damage bypass by template switching. Nature Structural & Molecular Biology 21, 884–892.

Gonzalez-Prieto, R., Munoz-Cabello, A.M., Cabello-Lobato, M.J., and Prado, F. (2013). Rad51 replication fork recruitment is required for DNA damage tolerance. EMBO J 32, 1307–1321.

Goodman, M.F., and Woodgate, R. (2013). Translesion DNA polymerases. Cold Spring Harbor Perspect Biol 5, a010363.

Halder, S., Sanchez, A., Ranjha, L., Reginato, G., Ceppi, I., Acharya, A., Anand, R., and Cejka, P. (2022). Double-stranded DNA binding function of RAD51 in DNA protection and its regulation by BRCA2. Mol Cell 82, 3553-+.

Hashimoto, Y., Chaudhuri, A.R., Lopes, M., and Costanzo, V. (2010). Rad51 protects nascent DNA from Mre11-dependent degradation and promotes continuous DNA synthesis. Nature Structural & Molecular Biology 17, 1305–U1268.

Hoege, C., Pfander, B., Moldovan, G.L., Pyrowolakis, G., and Jentsch, S. (2002). RAD6-dependent DNA repair is linked to modification of PCNA by ubiquitin and SUMO. Nature 419, 135–141.

Hofmann, R.M., and Pickart, C.M. (1999). Noncanonical MMS2-encoded ubiquitin-conjugating enzyme functions in assembly of novel polyubiquitin chains for DNA repair. Cell 96, 645–653.

Johnson, R.E., Kovvali, G.K., Prakash, L., and Prakash, S. (1998). Role of yeast Rth1 nuclease and its homologs in mutation avoidance, DNA repair, and DNA replication. Curr Genet 34, 21–29.

Karras, G.I., and Jentsch, S. (2010). The RAD6 DNA Damage Tolerance Pathway Operates Uncoupled from the Replication Fork and Is Functional Beyond S Phase. Cell 141, 255–267.

Ke, R., Ingram, P.J., and Haynes, K. (2013). An integrative model of ion regulation in yeast. PLoS Comp Biol 9, e1002879.

Kockler, Z.W., Osia, B., Lee, R., Musmaker, K., and Malkova, A. (2021). Repair of DNA Breaks by Break-Induced Replication. In Annual Review of Biochemistry, Vol 90, 2021, R.D. Kornberg, ed., pp. 165–191.

Kolinjivadi, A.M., Sannino, V., De Antoni, A., Zadorozhny, K., Kilkenny, M., Techer, H., Baldi, G., Shen, R., Ciccia, A., Pellegrini, L., et al. (2017). Smarcal1-Mediated Fork Reversal Triggers Mre11-Dependent Degradation of Nascent DNA in the Absence of Brca2 and Stable Rad51 Nucleofilaments. Mol Cell 67, 867–881 e867.

Kowalczykowski, S.C. (1991). Biochemistry of genetic recombination: energetics and mechanism of DNA strand exchange. Annu Rev Biophys Biophys Chem 20, 539–575.

Kowalczykowski, S.C. (2015). An Overview of the Molecular Mechanisms of Recombinational DNA Repair. Cold Spring Harbor Perspectives in Biology 7, a016410.

Kowalczykowski, S.C., Clow, J., and Krupp, R.A. (1987). Properties of the duplex DNA-dependent ATPase activity of *Escherichia coli* RecA protein and its role in branch migration. Proc Natl Acad Sci U S A 84, 3127–3131.

Kowalczykowski, S.C., Hunter, N., and Heyer, W.-D., eds. (2016). DNA Recombination (Cold Spring Harbor Laboratory Press).

Krejci, L., Van Komen, S., Li, Y., Villemain, J., Reddy, M.S., Klein, H., Ellenberger, T., and Sung, P. (2003). DNA helicase Srs2 disrupts the Rad51 presynaptic filament. Nature 423, 305–309.

Laemmli, U.K. (1970). Cleavage of structural proteins during the assembly of the head of bacteriophage T4. Nature 227, 680–685.

Laughery, M.F., Hunter, T., Brown, A., Hoopes, J., Ostbye, T., Shumaker, T., and Wyrick, J.J. (2015). New vectors for simple and streamlined CRISPR-Cas9 genome editing in Saccharomyces cerevisiae.. Yeast 32, 711–720.

Lawrence, C.W., and Christensen, R.B. (1979). Metabolic suppressors of trimethoprim and ultraviolet light sensitivities of Saccharomyces cerevisiae rad6 mutants. J, Bacteriol 139, 866–876.

Lea, D.E., and Coulson, C.A. (1949). The distribution of the numbers of mutants in bacterial populations. J Genet 49, 399–406.

Lee, S.E., Pellicioli, A., Vaze, M.B., Sugawara, N., Malkova, A., Foiani, M., and Haber, J.E. (2003). Yeast Rad52 and Rad51 recombination proteins define a second pathway of DNA damage assessment in response to a single double-strand break. Mol Cell Biol 23, 8913–8923.

Lettier, G., Feng, Q., de Mayolo, A.A., Erdeniz, N., Reid, R.J.D., Lisby, M., Mortensen, U.H., and Rothstein, R. (2006). The role of DNA double-strand breaks in spontaneous homologous recombination in S. cerevisiae. Plos Genetics 2, 1773–1786.

Li, X., and Heyer, W.D. (2008). Homologous recombination in DNA repair and DNA damage tolerance. Cell Res 18, 99–113.

Li, X., Zhang, X.P., Solinger, J.A., Kiianitsa, K., Yu, X., Egelman, E.H., and Heyer, W.D. (2007). Rad51 and Rad54 ATPase activities are both required to modulate Rad51-dsDNA filament dynamics. Nucleic Acids Res 35, 4124–4140.

Liberi, G., Maffioletti, G., Lucca, C., Chiolo, I., Baryshnikova, A., Cotta-Ramusino, C., Lopes, M., Pellicioli, A., Haber, J.E., and Foiani, M. (2005). Rad51-dependent DNA structures accumulate at damaged replication forks in sgs1 mutants defective in the yeast ortholog of BLM RecQ helicase. Genes Dev 19, 339–350.

Liu, J., Gore, S.K., and Heyer, W.-D. (2024). Local structural dynamics of Rad51 protomers revealed by cryo-electron microscopy of Rad51-ssDNA filaments. bioRxiv 2024.05.06.592824; doi: 10.1101/2024.05.06.592824

Liu, J., Renault, L., Veaute, X., Fabre, F., Stahlberg, H., and Heyer, W.-D. (2011a). Rad51 paralogs Rad55-Rad57 balance the anti-recombinase function of Srs2 in Rad51 pre-synaptic filament formation. Nature 479, 245–248.

Liu, J., Sneeden, J., and Heyer, W.D. (2011b). In vitro assays for DNA pairing and recombination-associated DNA synthesis. Methods in Molecular Biology 745, 363–383.

Liu, W.P., Saito, Y., Jackson, J., Bhowmick, R., Kanemaki, M.T., Vindigni, A., and Cortez, D. (2023). RAD51 bypasses the CMG helicase to promote replication fork reversal. Science 380, 382-+.

Loeillet, S., Herzog, M., Puddu, F., Legoix, P., Baulande, S., Jackson, S.P., and Nicolas, A.G. (2020). Trajectory and uniqueness of mutational signatures in yeast mutators. Proceedings of the National Academy of Sciences of the United States of America 117, 24947–24956.

Makarova, A.V., and Burgers, P.M. (2015). Eukaryotic DNA polymerase zeta. DNA Repair 29, 47–55.

Maloisel, L., Ma, E., Phipps, J., Deshayes, A., Mattarocci, S., Marcand, S., Dubrana, K., and Coic, E. (2023). Rad51 filaments assembled in the absence of the complex formed by the Rad51 paralogs Rad55 and Rad57 are outcompeted by translesion DNA polymerases on UV-induced ssDNA gaps. Plos Genetics 19.

Maslowska, K.H., Laureti, L., and Pages, V. (2019). iDamage: a method to integrate modified DNA into the yeast genome. Nucleic Acids Res 47.

Maslowska, K.H., Makiela-Dzbenska, K., Fijalkowska, I.J. (2019). The SOS system: A complex and tightly regulated response to DNA damage Environ Mol Mutagen 60, 368–384.

Maslowska, K.H., and Pages, V. (2022). Rad5 participates in lesion bypass through its Rev1-binding and ubiquitin ligase domains, but not through its helicase function. Frontiers in Molecular Biosciences 9.

Maslowska, K.H., and Pages, V. (2023). Cells limit mutagenesis by dealing with DNA lesions behind the fork. bioRxiv 2023*.09.04.556208; doi:* 10.1101/2023.09.04.556208

Mason, J.M., Chan, Y.L., Weichselbaum, R.W., and Bishop, D.K. (2019). Non-enzymatic roles of human RAD51 at stalled replication forks. Nature Communications 10.

Mason, J.M., Dusad, K., Wright, W.D., Grubb, J., Budke, B., Heyer, W.-D., Connell, P.P., Weichselbaum, R.R., and Bishop, D.K. (2015). RAD54 family translocases counter genotoxic effects of RAD51 in human tumor cells.. Nucleic Acids Res 43, 3180–3196.

Mijic, S., Zellweger, R., Chappidi, N., Berti, M., Jacobs, K., Mutreja, K., Ursich, S., Chaudhuri, A.R., Nussenzweig, A., Janscak, P., et al. (2017). Replication fork reversal triggers fork degradation in BRCA2-defective cells. Nature Communications 8.

New, J.H., Sugiyama, T., Zaitseva, E., and Kowalczykowski, S.C. (1998). Rad52 protein stimulates DNA strand exchange by Rad51 and replication protein A. Nature 391, 407–410.

Nimonkar, A.V., Dombrowski, C.C., Siino, J.S., Stasiak, A.Z., Stasiak, A., and Kowalczykowski, S.C. (2012). Saccharomyces cerevisiae Dmc1 and Rad51 Proteins Preferentially Function with Tid1 and Rad54 Proteins, Respectively, to Promote DNA Strand Invasion during Genetic Recombination. J Biol Chem 287, 28727–28737.

Papouli, E., Chen, S.H., Davies, A.A., Huttner, D., Krejci, L., Sung, P., and Ulrich, H.D. (2005). Crosstalk between SUMO and ubiquitin on PCNA is mediated by recruitment of the helicase Srs2p. Mol Cell 19, 123–133.

Patel, M., Jiang, Q.F., Woodgate, R., Cox, M.M., and Goodman, M.F. (2010). A new model for SOS-induced mutagenesis: how RecA protein activates DNA polymerase V. Critical Reviews in Biochemistry and Molecular Biology 45, 171–184.

Petukhova, G., Stratton, S., and Sung, P. (1998). Catalysis of homologous DNA pairing by yeast Rad51 and Rad54 proteins. Nature 393, 91–94.

Pfander, B., Moldovan, G.L., Sacher, M., Hoege, C., and Jentsch, S. (2005). SUMO-modified PCNA recruits Srs2 to prevent recombination during S phase. Nature 436, 428–433.

Piazza, A., Shah, S.S., Wright, W.D., Gore, S.K., Koszul, R., and Heyer, W.D. (2019). Dynamic Processing of Displacement Loops during Recombinational DNA Repair. Mol Cell 73, 1255–1266.

Rattray, A.J., Shafer, B.K., McGill, C.B., and Strathern, J.N. (2002). The roles of REV3 and RAD57 in double-strand-break-repair-induced mutagenesis of Saccharomyces cerevisiae. Genetics 162, 1063–1077.

Schiestl, R.H., Prakash, S., and Prakash, L. (1990). The *SRS2* suppressor of *rad6* mutations of *Saccharomyces cerevisiae* acts by channeling DNA lesions into the *RAD52* DNA repair pathway. Genetics 124, 817–831.

Schlacher, K., Christ, N., Siaud, N., Egashira, A., Wu, H., and Jasin, M. (2011). Double-Strand Break Repair-Independent Role for BRCA2 in Blocking Stalled Replication Fork Degradation by MRE11. Cell 145, 529–542.

Segurado, M., and Diffley, J.F.X. (2008). Separate roles for the DNA damage checkpoint protein kinases in stabilizing DNA replication forks. Genes Dev 22, 1816–1827.

Shah, P.P., Zheng, X.Z., Epshtein, A., Carey, J.N., Bishop, D.K., and Klein, H.L. (2010). Swi2/Snf2-Related Translocases Prevent Accumulation of Toxic Rad51 Complexes during Mitotic Growth. Mol Cell 39, 862–872.

Shinohara, A., Ogawa, H., and Ogawa, T. (1992). Rad51 protein involved in repair and recombination in *S. cerevisiae* is a RecA-like protein. Cell 69, 457–470.

Smirnova, M., and Klein, H.L. (2003). Role of the error-free damage bypass postreplication repair pathway in the maintenance of genomic stability. Mutation Research - Fundamental & Molecular Mechanisms of Mutagenesis 532, 117–135.

Smith, J., and Rothstein, R. (1999). An allele of RFA1 suppresses RAD52-dependent double-strand break repair in Saccharomyces cerevisiae. Genetics 151, 447–458.

Sogo, J.M., Lopes, M., and Foiani, M. (2002). Fork reversal and ssDNA accumulation at stalled replication forks owing to checkpoint defects. Science 297, 599–602.

Solinger, J.A., Kiianitsa, K., and Heyer, W.-D. (2002). Rad54, a Swi2/Snf2-like recombinational repair protein, disassembles Rad51:dsDNA filaments. Mol Cell 10, 1175–1188.

Stelter, P., and Ulrich, H.D. (2003). Control of spontaneous and damage-induced mutagenesis by SUMO and ubiquitin conjugation. Nature 425, 188–191.

Sung, P. (1994). Catalysis of ATP-dependent homologous DNA pairing and strand exchange by yeast RAD51 protein. Science 265, 1241–1243.

Sung, P. (1997). Yeast Rad55 and Rad57 proteins form a heterodimer that functions with replication protein A to promote DNA strand exchange by Rad51 recombinase. Genes Dev 11, 1111–1121.

Sung, P., and Robberson, D.L. (1995). DNA strand exchange mediated by a RAD51-ssDNA nucleoprotein filament with polarity opposite to that of RecA. Cell 82, 453–461.

Taglialatela, A., Alvarez, S., Leuzzi, G., Sannino, V., Ranjha, L., Huang, J.W., Madubata, C., Anand, R., Levy, B., Rabadan, R., et al. (2017). Restoration of Replication Fork Stability in BRCA1-and BRCA2-Deficient Cells by Inactivation of SNF2-Family Fork Remodelers. Mol Cell 68, 414-+.

Tombline, G., Heinen, C.D., Shim, K.S., and Fishel, R. (2002). Biochemical characterization of the human RAD51 protein - III. - Modulation of DNA binding by adenosine nucleotides. J Biol Chem 277, 14434–14442.

Urulangodi, M., Sebesta, M., Menolfi, D., Szakal, B., Sollier, J., Sisakova, A., Krejci, L., and Branzei, D. (2015). Local regulation of the Srs2 helicase by the SUMO-like domain protein Esc2 promotes recombination at sites of stalled replication. Genes Dev 29, 2067–2080.

Veaute, X., Jeusset, J., Soustelle, C., Kowalczykowski, S.C., Le Cam, E., and Fabre, F. (2003). The Srs2 helicase prevents recombination by disrupting Rad51 nucleoprotein filaments. Nature 423, 309–312.

Wong, R.P., Garcia-Rodriguez, N., Zilio, N., Hanulova, M., and Ulrich, H.D. (2020). Processing of DNA Polymerase-Blocking Lesions during Genome Replication Is Spatially and Temporally Segregated from Replication Forks. Mol Cell 77, 3-+.

Xu, X., Ball, L., Chen, W.Y., Tian, X.L., Lambrecht, A., Hanna, M., and Xiao, W. (2013). The Yeast Shu Complex Utilizes Homologous Recombination Machinery for Error-free Lesion Bypass via Physical Interaction with a Rad51 Paralogue. PloS one 8.

Xu, X., Lin, A.Y., Zhou, C.Y., Blackwell, S.R., Zhang, Y.R., Wang, Z.H., Feng, Q.Q., Guan, R.F., Hanna, M.D., Chen, Z.C., et al. (2016). Involvement of budding yeast Rad5 in translesion DNA synthesis through physical interaction with Rev1. Nucleic Acids Res 44, 5231–5245.

Zaitseva, E.M., Zaitsev, E.N., and Kowalczykowski, S.C. (1999). The DNA binding properties of *Saccharomyces cerevisiae* Rad51 protein. J Biol Chem 274, 2907–2915.

Zellweger, R., Dalcher, D., Mutreja, K., Berti, M., Schmid, J.A., Herrador, R., Vindigni, A., and Lopes, M. (2015). Rad51-mediated replication fork reversal is a global response to genotoxic treatments in human cells. J Cell Biol 208, 563–579.

