## Supplement for "Rad51 determines pathway usage in post-replication repair"

^1^ Department of Microbiology & Molecular Genetics and ^5^ Department of Molecular & Cellular Biology, University of California, Davis, Davis CA 95616-8665, USA; ^2^ Institute for Research in Biomedicine, Università della Svizzera italiana (USI), Faculty of Biomedical Sciences, 6500 Bellinzona, Switzerland; ^3^ Centro Andaluz de Biología Molecular y Medicina Regenerativa –CABIMER; Consejo Superior de Investigaciones Científicas; Universidad de Sevilla; Universidad Pablo de Olavide; Seville, Spain; ^4^ Cancer Research Center of Marseille: Team DNA Damage and Genome Instability, CNRS, Aix Marseille Univ, Inserm, Institut Paoli-Calmettes, Marseille F-13009, France.

Address correspondence to:

Wolf-Dietrich Heyer, Department of Microbiology & Molecular Genetics, University of California, Davis, One Shields Ave., Davis, California 95616-8665, Tel. 530-752-3001; Fax. 530-752-3011;

Supplemental Tables S1-4

Supplemental Movie S1

Supplemental References

Supplemental Figures S1-8

**Supplemental Table S1: *Saccharomyces cerevisiae* strains**

**Number	Genotype	Source**

WDHY1275	*MATa ade2-1 can1-100 his3-11,15 leu2-3, 112 trp1-1 ura3-1*

*rad54::KANMX*	This study

WDHY1611	*MATa his3-∆1 leu2-3,112 trp1 ura3-52 pep4-3 rad51-∆::KanMX*	(1,2)

BJ strain for protein expression.

WDHY1636	*MATa ade2-1 can1-100 his3-11,15 leu2-3,112 trp1-1 ura3-1*	(3)

WDHY2217	*MATa ade2-1 can1-100 his3-11,15 leu2-3,112 trp1-1 ura3::loxP*	This study

WDHY2542	*MATa ade2-1 can1-100 his3-11,15 leu2-3, 112 trp1-1 ura3-1*

*rad51::KANMX*	This study

WDHY2543	*MATa ade2-1 can1-100 his3-11,15 leu2-3,112 trp1-1 ura3-1
rad51::KANMX*	This study

WDHY2544	*MATa ade2-1 can1-100 his3-11,15 leu2-3,112 trp1-1 ura3-1*

*rad51Δ::KANMX rad54Δ::KANMX*	This study

WDHY2546	*MATa ade2-1 can1-100 his3-11,15 leu2-3,112 trp1-1 or trp1-289*

*ura3-1 or ura 3-52 rad51Δ::KANMX rad54-3^ts^(-C692Y)*	This study

WDHY3162	*MATa ade2-1 CAN1 his3::(delta)5’his3(delta)3’:: URA3 leu2-3, 112*

*trp1-1 ura3-1 rad51::LEU2 rad54::KANMX*	This study

WDHY3169	*MATa ade2-1 CAN1 his3::(delta)5’his3(delta)3’:: URA3 leu2-3, 112*

*trp1-1 ura3-1 rad54::KANMX*	This study

WDHY3348	*MATa/MATα ade2-1/ade2-1 can1-100/can1-100 his3-11,15/his3-11,15*

*leu2(delta)EcoRI-URA3-HOcs(117)-leu2(delta) BstEII/leu2-3,112*

*trp1-1/trp1::KANMX-GALHO ura3-1/ura3-1 rad52::TRP1/RAD52*	This study

WDHY3349	*MATa/MATα ade2-1/ade2-1 can1-100/can1-100 HIS3/his3-11,15*

*leu2(delta)EcoRI-URA3-HOcs(117)-leu2(delta) BstEII/leu2-3,112*

*trp1-1/trp1::KANMX-GALHO ura3-1/ura3-1 rad54::KANMX/RAD54*	This study

WDHY3383	*MATa/MATα ade2-1/ade2-1 can1-100/can1-100 his3Δ200/his3-11,15*

*leu2(delta)EcoRI-URA3-HOcs(117)-leu2(delta) BstEII/leu2-3,112*

*trp1-1/trp1::KANMX-GALHO ura3-1/ura3-1*	This study

WDHY3385	*MATa/MATα ade2-1/ade2-1 can1-100/can1-100 his3-11,15/his3-11,15*

*leu2(delta)EcoRI-URA3-HOcs(117)-leu2(delta) BstEII/leu2-3,112*

*trp1-1/trp1::KANMX-GALHO ura3-1/ura3::TRP1 RAD51/rad51-K305N*	This study

WDHY3462	*MATa/MATα ade2-1/ade2-1 can1-100/can1-100 his3-11,15/his3-11,15*

*leu2(delta)EcoRI-URA3-HOcs(117)-leu2(delta) BstEII/leu2-3,112*

*trp1-1/trp1::KANMX-GALHO ura3-1/ura3::TRP1*

*RAD51/rad51-E135D RAD54/rad54::KANMX*	This study

WDHY3463	*MATa/MATα ade2-1/ade2-1 can1-100/can1-100 his3-11,15/his3-11,15*

*leu2(delta)EcoRI-URA3-HOcs(117)-leu2(delta)BstEII/leu2-3,112*

*trp1-1/trp1::KANMX-GALHO ura3-1/ura3::TRP1*

*RAD51/rad51-K305N RAD54/rad54::KANMX*	This study

WDHY3536	*MATa ade2-1 can1-100 HIS3 trp1-1 leu2-3,112 ura3-1*

*rad5-G535R rad51-E135D*	This study

WDHY3539	*MATa ade2-1 can1-100 his3-11,15 TRP1 leu2-3,112 ura3-1*

*rad5-G535R rad51-E135D rad54::KANMX*	This study

WDHY3546	*MATα ade2-1 can1-100 his3-11,15 trp1-1 leu2-3,112 ura3-1*

*rad51-E135D rad54::KANMX*	This study

WDHY3547	*MATα ade2-1 can1-100 his3-11,15 trp1-1 leu2-3,112 ura3-1*

*rad51-E135D rad54::KANMX*	This study

WDHY3548	*MATa ade2-1 can1-100 his3-11,15 trp1-1 leu2-3,112 ura3-1*

*rad51-E135D*	This study

WDHY3555	*MATa ade2-1 can1-100 leu2-3,112 ura3-1 pol30-K127R,K164R*	This study

WDHY3557	*MATa ade2-1 can1-100 his3-11,15 leu2-3,112 ura3-1 rad51-K305N*

*pol30-K127R,K164R*	This study

WDHY3561	*MATα ade2-1 can1-100 his3-11,15 leu2-3,112 ura3-1 rad51-E135D*

*pol30-K127R,K164R*	This study

WDHY3563	*MATα ade2-1 can1-100 leu2-3,112 ura3-1 rad54::KANMX*

*pol30-K127R,K164R*	This study

WDHY3568	*MATa ade2-1 can1-100 his3-11,15 leu2-3,112 ura3-1 rad51-K305N*

*rad54::KANMX pol30-K127R,K164R*	This study

WDHY3570	*MATa ade2-1 can1-100 his3-11,15 leu2-3,112 ura3-1 rad51-E135D*

*rad54::KANMX pol30-K127R,K164R*	This study

WDHY3572	*MATa ade2-1 can1-100 his3-11,15 trp1-1 leu2-3,112 ura3-1*

*rad51-K305N rad54::KANMX*	This study

WDHY3578	*MATa ade2-1 can1-100 his3-11,15 trp1-1 leu2-3,112 ura3-1*

*rad5-G535R rad51-K305N*	This study

WDHY3581	*MATα ade2-1 can1-100 his3-11,15 TRP1 leu2-3,112 ura3-1*

*rad5-G535R rad51-K305N rad54::KANMX*	This study

WDHY3584	*MATa ade2-1 can1-100 trp1-1 leu2-3,112 ura3-1 rev3::KANMX*	This study

WDHY3588	*MATα ade2-1 can1-100 trp1-1 leu2-3,112 ura3-1 his3-11,15*

*rad54::KANMX* *rev3::KANMX*	This study

WDHY3589	*MATa ade2-1 can1-100 trp1-1 leu2-3,112 ura3-1 rad51-K305N*

*rev3::KANMX*	This study

WDHY3777	*MATa/MATα ade2-1/ade2-1 can1-100/can1-100 HIS3/his3-11,15*

*leu2(delta)EcoRI-URA3-HOcs(117)-leu2(delta) BstEII/leu2-3,112*

*trp1-1/trp1::KANMX-GALHO ura3-1/ura3-1 rad51E135D/RAD51*	This study

WDHY3780	*MATα ade2-1 can1-100 trp1-1 leu2-3,112 ura3-1 his3-11,15*

*rad51::KANMX rad54::KANMX pol30-K127R,K164R*	This study

WDHY3783	*MATα ade2-1 can1-100 trp1::KANMX-GAL-HO leu2-3,112 ura3-1*

*his3-11,15 rad51::LEU2 pol30-K127R,K164R*	This study

WDHY3784	*MATa ade2-1 can1-100 leu2-3,112 ura3-1 his3-11,15 rad51-E135D*

*rev3::KANMX*	This study

WDHY3785	*MATα ade2-1 can1-100 trp1-1 leu2-3,112 his3-11,15 trp1-1*

*rad51-E135D rad54:KANMX rev3::KANMX*	This study

WDHY3787	*MATa ade2-1 can1-100 trp1-1 leu2-3,112 his3-11,15 trp1-1*

*rad51::LEU2 rev3::KANMX*	This study

WDHY3788	*MATα ade2-1 can1-100 trp1-1 leu2-3,112 ura3-1 his3-11,15*

*rad51::LEU2 rad54::KANMX rev3::KANMX*	This study

WDHY3789	*MATa ade2-1 can1-100 trp1-1 leu2-3,112 ura3-1 his3-11,15*

*rad51-K305N rad54::KANMX rev3::KANMX*	This study

WDHY3855	*MATα ade2-1 can1-100 HIS3 leu2-3,112 trip1-1 ura3-1*

*mms2::KANMX*	This study

WDHY3898	*MATa ade2-1 can1-100 his3-11,15 leu2-3,112 TRP1 ura3-1*

*rad51::HIS3*	This study

WDHY3915	*MATa/MATα ade2-1/ade2-1 can1-100/can1-100 his3-11,15/his3-11,15*

*leu2(delta)EcoRI-URA3-HOcs(117)-leu2(delta) BstEII/leu2-3,112*

*TRP1/trp1::KANMX-GALHO ura3-1/ura3-1 rad51::HIS3/RAD51*	This study

WDHY3952	*MATα ade2-1 can1-100 HIS3 leu2-3,112 trip1-1 ura3-1*

*mms2::KANMX	rad51::HIS3*	This study

WDHY3956	*MATα ade2-1 can1-100 HIS3 leu2-3,112 trip1-1 ura3-1*

*mms2::KANMX	rad54::KANMX*	This study

WDHY3958	*MATα ade2-1 can1-100 HIS3 leu2-3,112 trip1-1 ura3-1*

*mms2::KANMX	rad51-K305N*	This study

WDHY3959	*MATα ade2-1 can1-100 his3-11, 15 leu2-3,112 ura3-1 trp1-1*

*rad51::HIS3 rad54::KanMX*	This study

WDHY3960	*MATa ade2-1 can1-100 his3-11, 15 leu2-3,112 ura3-1 trp1-1*	This study

WDHY3961	*MATα* *ade2-1 can1-100 his3-11 15 leu2-3,112 ura3-1 trp1-1*

*rad54::kanMX*	This study

WDHY3962	*MATα ade2-1 can1-100 his3-11,15 trp1-1 leu2-3,112 ura3-1*

*rad51-K305N*	This study

WDHY3963	*MATa ade2-1 can1-100 his3-11,15 trp1-1 leu2-3,112 ura3-1*

*rad51-K305N rad54::KanMX*	This study

WDHY4204	*MATa/MATα ade2-1/ade2-1 CAN1/can1-100 HIS3/his3-11,15*

*leu2-3,112/leu2-3,112 trp1-1/trp1-1 ura3-1/ura3-1*

*hxt13::URA3/HXT13 rad51::HIS3/RAD51*	This study

WDHY4205	*MATa/MATα ade2-1/ade2-1 CAN1/can1-100 HIS3/his3-11,15*

*leu2-3,112/leu2-3,112 trp1-1/trp1-1 ura3-1/ura3-1*

*hxt13::URA3/HXT13 rad54::KANMX/RAD54*	This study

WDHY4207	*MATa/MATα ade2-1/ade2-1 CAN1/can1-100 HIS3/his3-11,15*

*leu2-3,112/leu2-3,112 trp1-1/trp1-1 ura3-1/ura3-1*

*hxt13::URA3/HXT13 rad51-E135D/RAD51*	This study

WDHY4208	*MATa/MATα ade2-1/ade2-1 CAN1/can1-100 HIS3/his3-11,15*

*leu2-3,112/leu2-3,112 trp1-1/trp1-1 ura3-1/ura3-1,*

*hxt13::URA3/HXT13 rad51-K305N/RAD51*	This study

WDHY4209	*MATa/MATα ade2-1/ade2-1 CAN1/can1-100 HIS3/his3-11,15*

*leu2-3,112/leu2-3,112 trp1-1/trp1-1 ura3-1/ura3-1*

*hxt13::URA3/HXT13 rad51-E135D/RAD51 rad54::KANMX/RAD54*	This study

WDHY4210	*MATa/MATα ade2-1/ade2-1 CAN1/can1-100 HIS3/his3-11,15*

*leu2-3,112/leu2-3,112 trp1-1/trp1-1, ura3-1/ura3-1*

*hxt13::URA3/HXT13 rad51-K305N/RAD51 rad54::KANMX/RAD54*	This study

WDHY4564	*MATa/MATα ade2-1/ade2-1 CAN1/can1-100 HIS3/his3-11,15*

*leu2-3,112/leu2-3,112 TRP1/trp1-1 ura3-1/ura3-1*

*mms2::KANMX/MMS2 rad51-E135D/RAD51 rad54::KANMX/RAD54*	This study

WDHY5169	*MATa ade2-1 can1-100 his3-11,15 leu2-3,112 trp1-1 ura3-1*

*rad51-E135D mms2::KANMX*	This study

WDHY6127	*MATa ade2-1 can1-100 his3-11,15 leu2-3,112 trp1-1 ura3-1*

*rad51-E135D rad54::KANMX mms2::KANMX*	This study

wR51MN-2	*MATa ade2-1 can1-100 his3-11,15 leu2-3,112 trp1-1 ura3-1*

*RAD51-MN::HIS3*	(4)

wR51-135MN	*MATa ade2-1 can1-100 leu2-3,112 trp1-1 ura3::loxP his3-11,15*

*rad51-E135D-MN::HIS3*	This study

wR51-305MN	*MATa ade2-1 can1-100 leu2-3,112 trp1-1 ura3::loxP his3-11,15*

*rad51-K305N-MN::HIS3*	This study

W303sgs1	*MATa ade2-1 can1-100 his3-11,15 leu2-3,112 trp1-1 ura3-1*

*sgs1::KANMX*	(5)

W303sr305	*MATa ade2-1 can1-100 his3-11,15 leu2-3,112 trp1-1 ura3*

*rad51-K305N* *sgs1::KANMX*	This study

W303sr135	*MATa ade2-1 can1-100 his3-11,15 leu2-3,112 trp1-1 ura3*

*rad51-E135D* *sgs1::KANMX*	This study

Unless otherwise noted, all strains have the W303 background and are wild type for *RAD5*.

SC53	*MATa his3-Δ1 leu2-3,112 trp1-Δ ura3-Δ met25-Δ rad14-Δ phr1-Δ*	(6) *msh2Δ::hisG VI(167260–167265):: (lox66-3′lacZ-MET25/lag)*

SC55	*MATa his3-Δ1 leu2-3,112 trp1-Δ ura3-Δ met25-Δ rad14-Δ phr1-Δ*	(6)

*msh2Δ::hisG VI(167260–167265):: (lox66-3′lacZ-MET25/lead)*

SC254	*MATa his3-Δ1 leu2-3,112 trp1-Δ ura3-Δ met25-Δ rad14-Δ phr1-Δ*	(6)

*msh2Δ::hisG rad51Δ::KAN*

*VI(167260–167265):: (lox66-3′lacZ-MET25/lag)*

SC255	*MATa his3-Δ1 leu2-3,112 trp1-Δ ura3-Δ met25-Δ rad14-Δ phr1-Δ*	(6)

*msh2Δ::hisG rad51Δ::KAN*

*VI(167260–167265):: (lox66-3′lacZ-MET25/lead)*

SC844	*MATa his3-Δ1 leu2-3,112 trp1-Δ ura3-Δ met25-Δ rad14-Δ phr1-Δ*	This study

*msh2Δ::hisG rad51-E135D*

*VI(167260–167265):: (lox66-3′lacZ-MET25/lag)*

SC845	*MATa his3-Δ1 leu2-3,112 trp1-Δ ura3-Δ met25-Δ rad14-Δ phr1-Δ*	This study

*msh2Δ::hisG rad51E135D*

*VI(167260–167265):: (lox66-3′lacZ-MET25/lead)*

SC868	*MATa his3-Δ1 leu2-3,112 trp1-Δ ura3-Δ met25-Δ rad14-Δ phr1-Δ*	This study *msh2Δ::hisG rad51-K305N*

*VI(167260–167265):: (lox66-3′lacZ-MET25/lag)*

SC869	*MATa his3-Δ1 leu2-3,112 trp1-Δ ura3-Δ met25-Δ rad14-Δ phr1-Δ*	This study

*msh2Δ::hisG rad51-K305N*

*VI(167260–167265):: (lox66-3′lacZ-MET25/lead)*

All SC strains share the EMY74.7 background (7).

WDHY668	*MATa/α ura3-52/ ura3-52 trp1/ trp1 leu2∆1/ leu2∆1 his3∆200/ his3∆200*	(8)

*pep4::HIS3/ pep4::HIS3 prb1∆1.6R/ prb1∆1.6R can1/ can1 GAL/ GAL*

Strain for protein purification with BJ background (2).

**Supplemental Table S2: Plasmids**

**Number	Description	Source**

WDH647	pR51.3 with wild type *RAD51* for protein expression	(9)

pWDH951	pR51.3 with *rad51-E135D* for protein expression	This study

pWDH952	pR51.3 with *rad51-K305N* for protein expression	This study

pWDH953	YEp351-*rad51-K305N +*1,000 bp up/downstream of ORF	This study

pWDH954	YEp351-*rad51-E135D +*1,000 bp up/downstream of ORF	This study

pWDH957	YEp351-*RAD51 +*1,000 bp up/downstream of ORF	This study

pWDH958	YEp351	*LEU2 amp^R^*	(10)

pFB-MBP-Sgs1-his	Expression of MBP- and His-tagged Sgs1 in insect cells	(11)

pFB-Exo1-FLAG	Expression of FLAG-tagged Exo1 in insect cells	(12)

pGAL:FLAG-DNA2-his	Expression of FLAG-, HA- and His-tagged Dna2 in yeast	(13)

p11d–tRPA	Expression of Rfa1, Rfa2 and Rfa3 in bacteria	(14)

**Supplemental Table S3: Oligonucleotides**

**Number	Sequence and use	Source**

olWDH566	5’- ATGGCAGCACTGCATAATTCTCTTACTGTCATGCCATCCGTAAGATG	Operon

CTTTTCTGTGACTGGTGAGTACTCAACCAAGTCATTCTGAGAATAGTG

D-loop assay

olWDH632	5’- AATGGACGGTAAATGTTGGA	Operon

Forward primer for random *RAD51* mutagenesis and

*in vivo* recombination, ~400 nt upstream

olWDH633	5’- AACGTCGAAACGAAGACAAG	Operon

Reverse primer for random *RAD51* mutagenesis and

*in vivo* recombination, ~ 400 nt downstream

olWDH830	5’- CCCGAGCTCCTCAGCGAAGTCGTGAAACTCGGA	Invitrogen

Reverse primer 1,000 downstream *RAD51* open reading

frame with *Sac*I site

olWDH1351	5’- AATTCCAGCTGACCACCATGATGTCTCAAGTTCAAGAACAA	Invitrogen

Forward *RAD51* adaptamer for mutant integration

olWDH1352	5’- GATCCCCGGGAATTGCCATGAGAATTGAAAGTAAACCTGTG	Invitrogen

Reverse *RAD51* adaptamer for mutant integration

olWDH1355	5’- CCCAAGCTTCCGCAATAAAGGGCTTCCCGGACT	Invitrogen

Forward primer 1,000 upstream of *RAD51* open reading
frame with *Hind*III site

olWDH1357	5’- TGCAATATCAACGGTACCCTTAGT	Invitrogen

Internal *K. lactis URA3* primer

olWDH1358	5’- GGTCCATACATTTGCCTTTTGAAA	Invitrogen

Internal *K. lactis URA3* primer

BIO100C	5’- GATGCAGGAGGCTGCTACGACCATGGCAGAAGATTATGAGGTGGAGT	Eurogentec

ACGCGCCCGGGGAGCCCAAGGGCACGCCCTGGCACCCGCACCGCGGCACTTAC	100 nt oligonucleotide for the 100 bp substrate used in helicase and

nuclease assays

BIO100	5’- G**T**AAGTGCCGCGGTGCGGGTGCCAGGGCGTGCCCTTGGGCTCCCCGG	Eurogentec

GCGCGTACTCCACCTCATAATCTTCTGCCATGGTCGTAGCAGCCTCCTGCATC

100 nt oligonucleotide for the 100 bp substrate used in helicase and

nuclease assays (the bold **T** represent the position of biotin-conjugated T)

**Supplemental Table 4:** **Confidence intervals for the median. Two sided Symmetric – 95% or better**

Nonparametric two-sided confidence intervals for the median of a continuous distribution based on order statistics. Shown are the narrowest symmetric intervals whose confidence coefficient is 95% or better.

Key

N	Sample size

L U	Order statistics defining the Lower and Upper endpoints

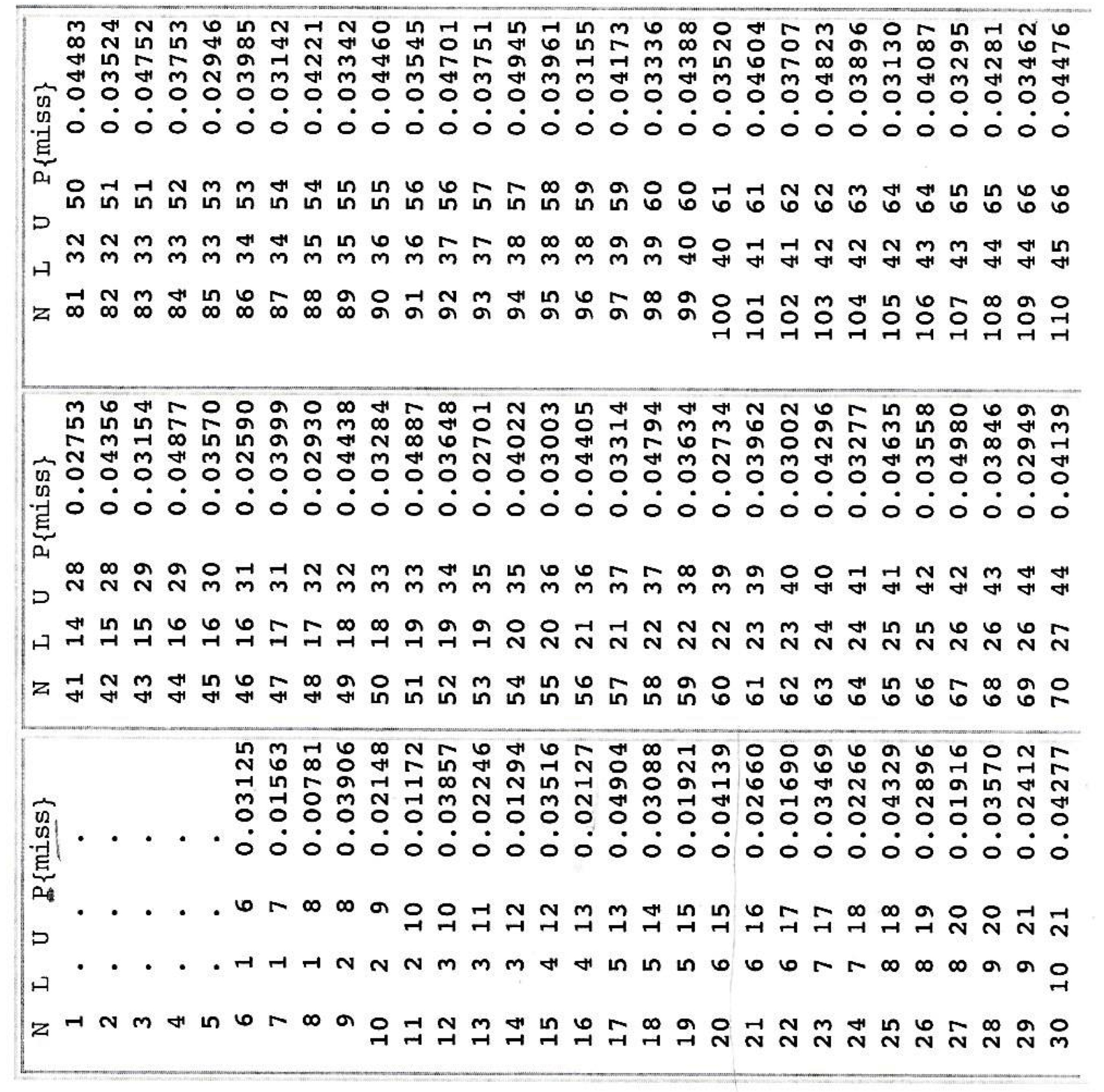
P (15)	Probability the interval does not cover the true median (never exceeds 0.05)

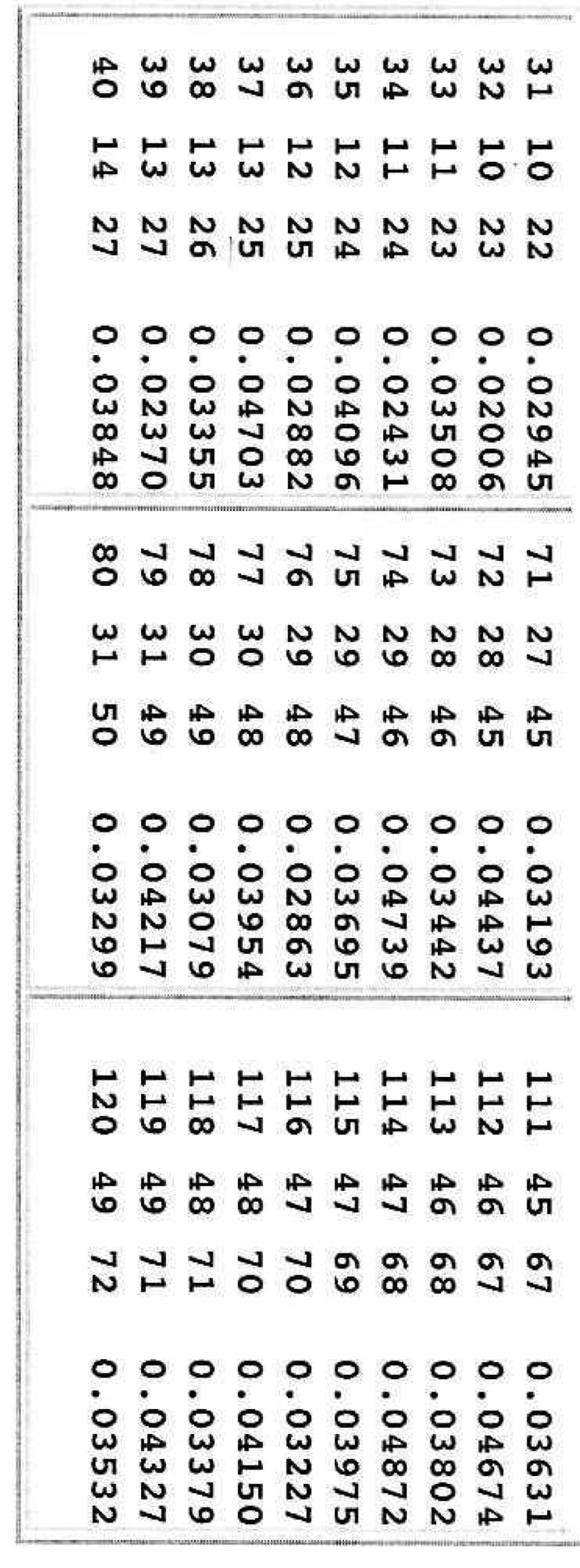

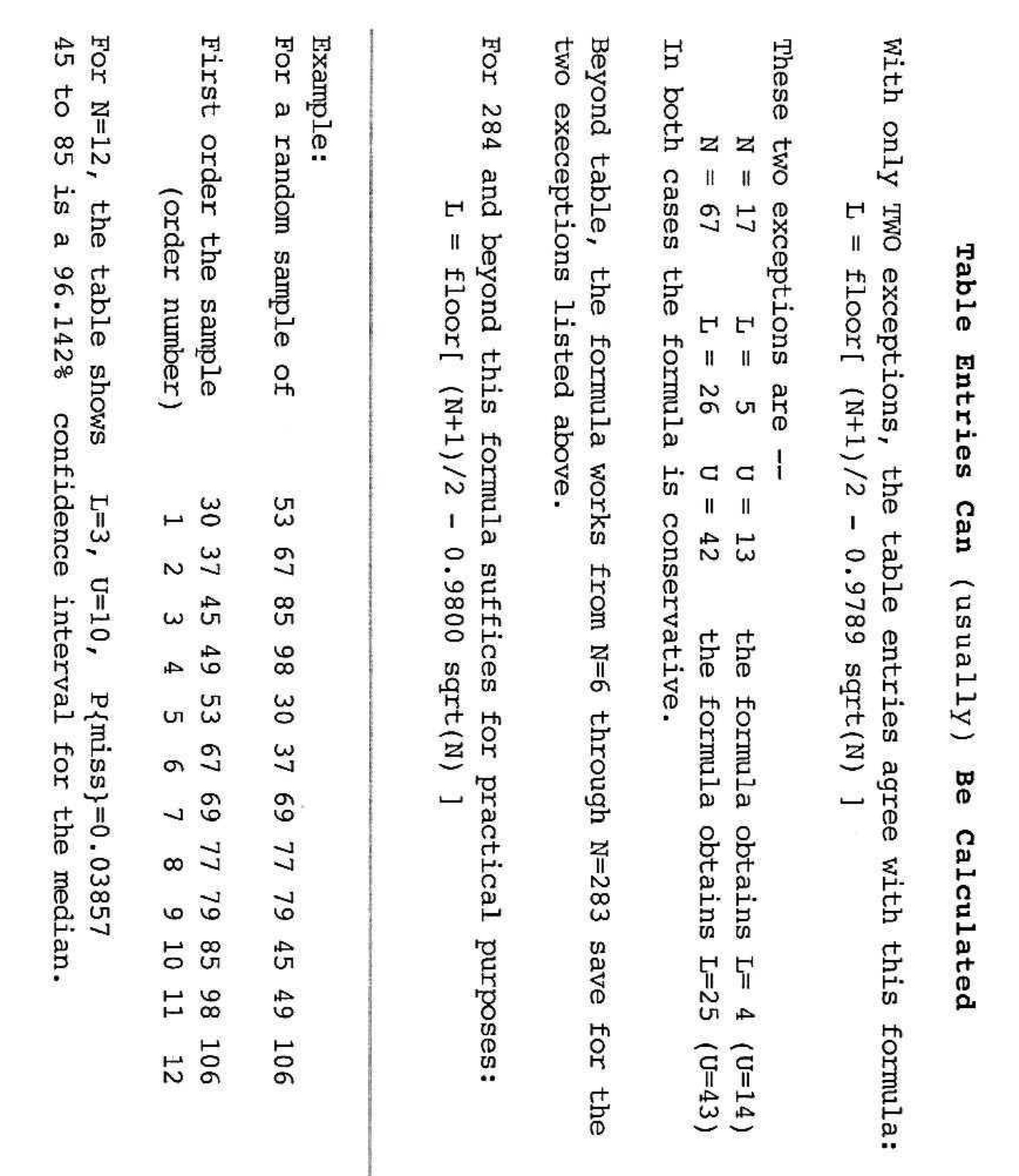

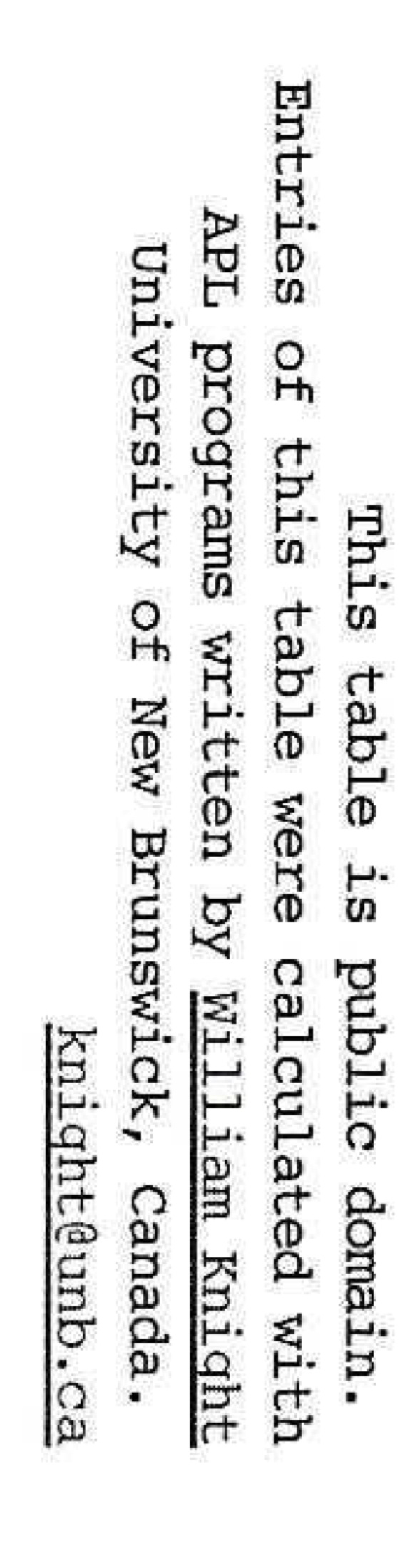

**Supplementary Movie 1: Locations of E135 and K305 in the Rad51-ssDNA filament structure with modeled dsDNA.** E135 and K305 are highlighted in red and labeled accordingly. The movie contains zoomed views to show the distance between dsDNA (in purple and green) and E135/K305 in rainbow-colored Rad51 protomers.

**Supplemental references**

1.	Zhang, X.P., Galkin, V.E., Yu, X., Egelman, E.H. and Heyer, W.D. (2009) Loop 2 in Saccharomyces cerevisiae Rad51 protein regulates filament formation and ATPase activity. *Nucleic Acids Res.*, **37**, 158-171.

2.	Jones, E.W. (1991) Tackling the protease problem in *Saccharomyces cerevisiae*. *Methods Enzymol.*, **194**, 428-453.

3.	Thomas, B.J. and Rothstein, R. (1989) The genetic control of direct-repeat recombination in Saccharomyces: the effect of rad52 and rad1 on mitotic recombination at GAL10, a transcriptionally regulated gene. *Genetics*, **123**, 725-738.

4.	Cabello-Lobato, M.J., Gonzalez-Garrido, C., Cano-Linares, M.I., Wong, R.P., Yanez-Vilchez, A., Morillo-Huesca, M., Roldan-Romero, J.M., Vicioso, M., Gonzalez-Prieto, R., Ulrich, H.D. *et al.* (2021) Physical interactions between MCM and Rad51 facilitate replication fork lesion bypass and ssDNA gap filling by non-recombinogenic functions. *Cell Rep*, **36**, 109440.

5.	Gonzalez-Prieto, R., Munoz-Cabello, A.M., Cabello-Lobato, M.J. and Prado, F. (2013) Rad51 replication fork recruitment is required for DNA damage tolerance. *EMBO J.*, **32**, 1307-1321.

6.	Maslowska, K.H., Laureti, L. and Pages, V. (2019) iDamage: a method to integrate modified DNA into the yeast genome. *Nucleic Acids Res.*, **47**.

7.	Johnson, R.E., Kovvali, G.K., Prakash, L. and Prakash, S. (1998) Role of yeast Rth1 nuclease and its homologs in mutation avoidance, DNA repair, and DNA replication. *Curr. Genet.*, **34**, 21-29.

8.	Solinger, J.A., Lutz, G., Sugiyama, T., Kowalczykowski, S.C. and Heyer, W.-D. (2001) Rad54 protein stimulates heteroduplex DNA formation in the synaptic phase of DNA strand exchange via specific interactions with the presynaptic Rad51 nucleoprotein filament. *J. Mol. Biol.*, **307**, 1207-1221.

9.	Sung, P. (1994) Catalysis of ATP-dependent homologous DNA pairing and strand exchange by yeast RAD51 protein. *Science*, **265**, 1241-1243.

10.	Hill, J.E., Myers, A.M., Koerner, T.J. and Tzagoloff, A. (1986) Yeast/E. coli shuttle vectors with multiple unique restriction sites. *Yeast*, **2**, 163-167.

11.	Cejka, P. and Kowalczykowski, S.C. (2010) The full-length *Saccharomyces cerevisiae* Sgs1 protein is a vigorous DNA helicase that preferentially unwinds Holliday junctions. *J Biol Chem*, **285**, 8290-8301.

12.	Cannavo, E., Cejka, P. and Kowalczykowski, S.C. (2013) Relationship of DNA degradation by *Saccharomyces* *cerevisiae* Exonuclease 1 and its stimulation by RPA and Mre11-Rad50-Xrs2 to DNA end resection. *Proc Natl Acad Sci USA*, **110**, E1661-E1668.

13.	Cejka, P., Cannavo, E., Polaczek, P., Masuda-Sasa, T., Pokharel, S., Campbell, J.L. and Kowalczykowski, S.C. (2010) DNA end resection by Dna2-Sgs1-RPA and its stimulation by Top3-Rmi1 and Mre11-Rad50-Xrs2. *Nature*, **467**, 112-116.

14.	Binz, S.K., Dickson, A.M., Haring, S.J. and Wold, M.S. (2006) Functional assays for replication protein A (RPA). *Methods Enzymol*, **409**, 11-38.

15.	Meddows, T.R., Savory, A.P., Grove, J.I., Moore, T. and Lloyd, R.G. (2005) RecN protein and transcription factor DksA combine to promote faithful recombinational repair of DNA double-strand breaks. **57**, 97-110.

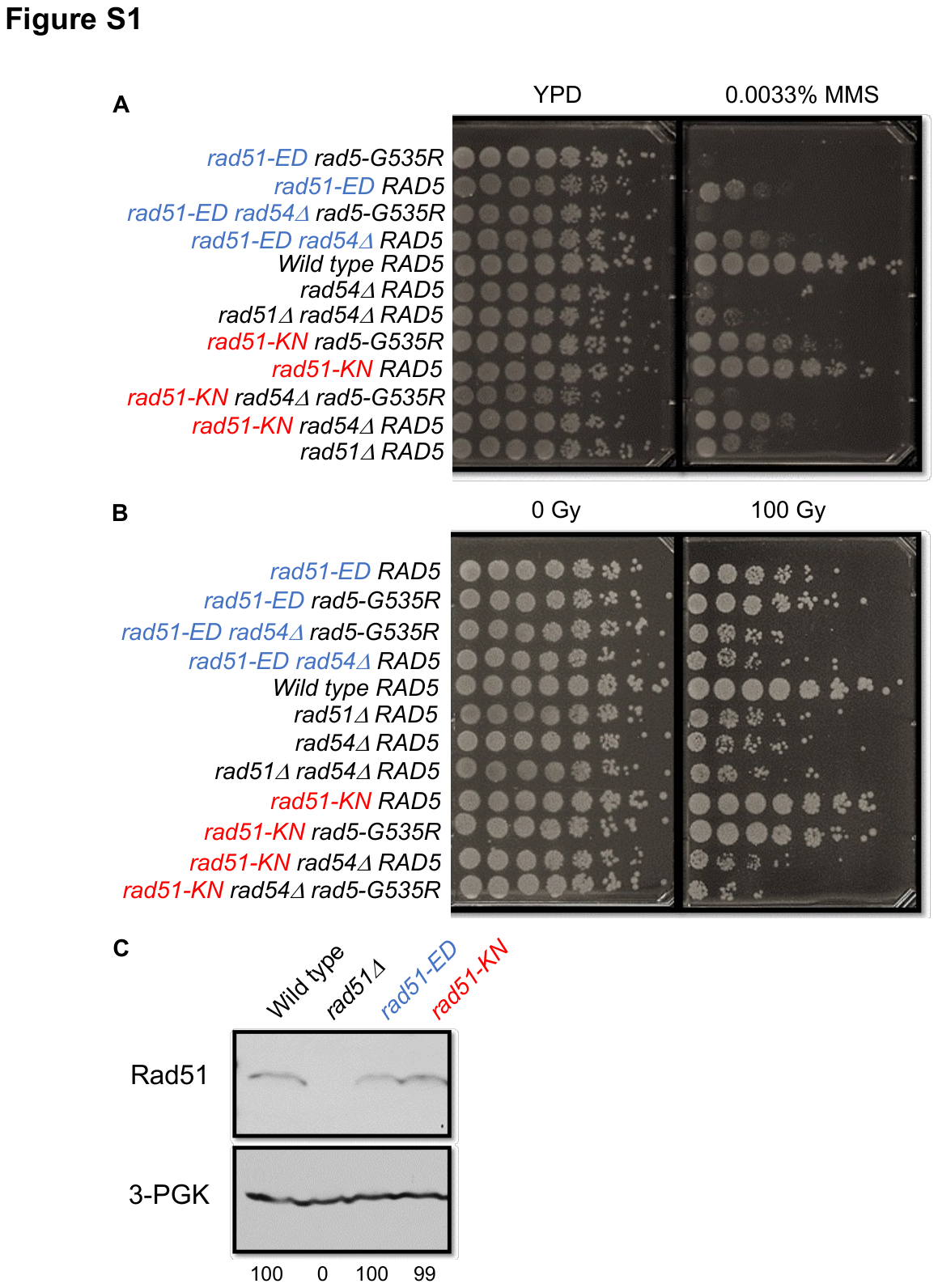

**Figure S1: MMS sensitivity and protein levels of *rad51* mutants. A, B.** Serial dilutions of strains with chromosomally integrated mutations: Wild-type *RAD5* (WDHY1636), *rad54Δ* *RAD5* (WDHY1275), *rad51Δ* *RAD5* (WDHY2542), *rad51Δ rad54Δ* *RAD5* (WDHY2544), *rad51-KN RAD5* (WDHY3962), *rad51-KN rad54Δ RAD5* (WDHY3572), *rad51-KN rad5-G535R* (WDHY3578), *rad51-KN* *rad54Δ rad5-G535R* (WDHY3581), *rad51-ED RAD5* (WDHY3548), *rad51-ED rad54Δ RAD5* (WDHY3546), *rad51-ED rad5-G535R* (WDHY3536), *rad51-ED rad54Δ rad5-G535R* (WDHY3539) to monitor sensitivity to 0.0033% MMS (**B**) or 100 Gy ionizing radiation (**B**). **C.** Rad51-E135E and Rad51-KN show wild type steady state protein levels. Immunoblot analysis of Rad51 protein levels in extracts from wild type *RAD51* (WDHY1636), *rad51Δ* (WDHY2543), *rad51-ED* (WDHY3548), and *rad51-KN* (WDHY3962) strains. Values below the lanes are the amount of Rad51 protein normalized to the 3-phosphoglycerate kinase (PGK) loading control expressed as a percentage of the wild-type.

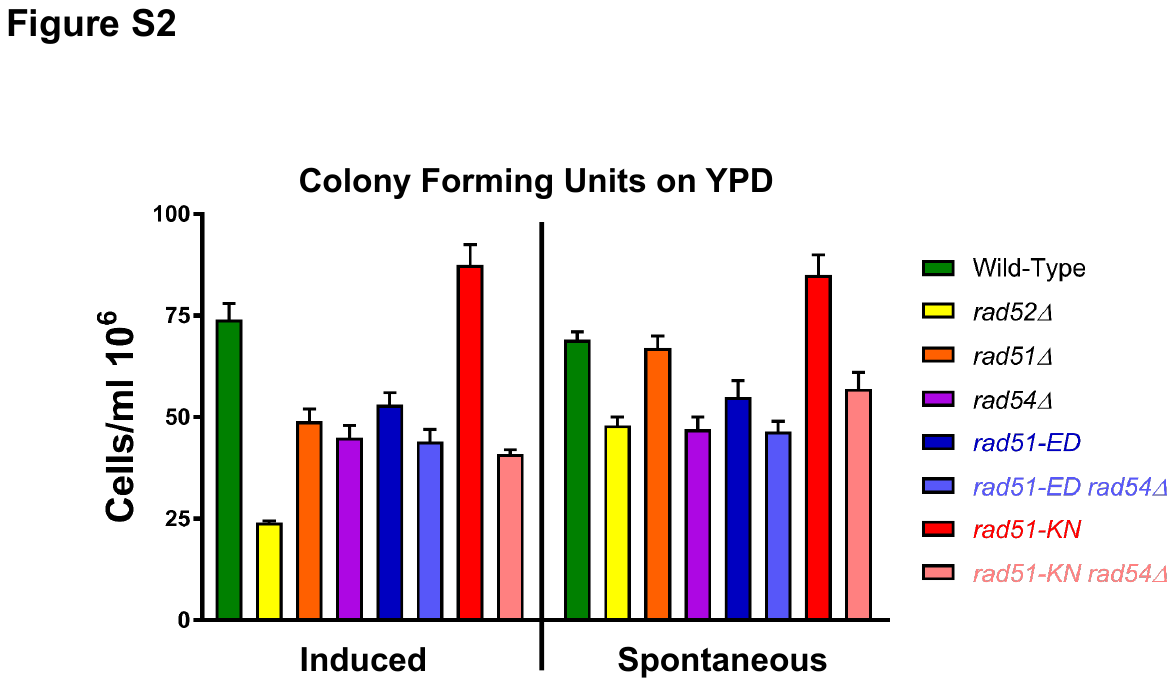

**Figure S2: Viability data for Figure 2 after DSB induction (induced) or without DSB induction (spontaneous).** The average number of colonies that grew on YPD was plotted to ascertain efficiency of plating and viability after DSB induction. Strains were freshly dissected from diploid strains and single spore clones were assayed, a minimum of 12 single clones were assayed per genotype. Diploids allowed for multiple strains to be created by dissection. Wildtype (WDHY3383), *rad52Δ* (WDHY3348), *rad51Δ* (WDHY3915)*, rad51-KN* (WDHY3385), *rad51-KN rad54Δ* (WDHY3463), *rad51-ED* (WDHY3777), *rad51-ED rad54Δ* (WDHY3462), *rad54Δ* (WDHY3349). Shown are means and error bars represent 1 standard deviation.

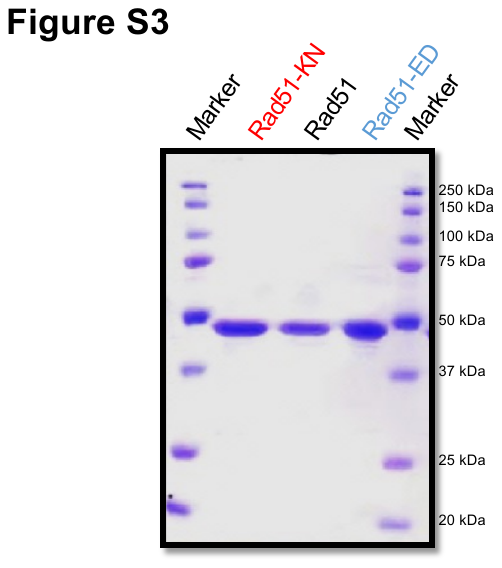

**Figure S3: Purification of Rad51-ED and Rad51-KN mutant proteins.** *Saccharomyces cerevisiae* wild type Rad51, Rad51-KN, and Rad51-ED proteins were purified and 1 µg each was analyzed by 10 % SDS-PAGE and stained with Coomassie brilliant blue. Molecular markers are in the outer lanes.

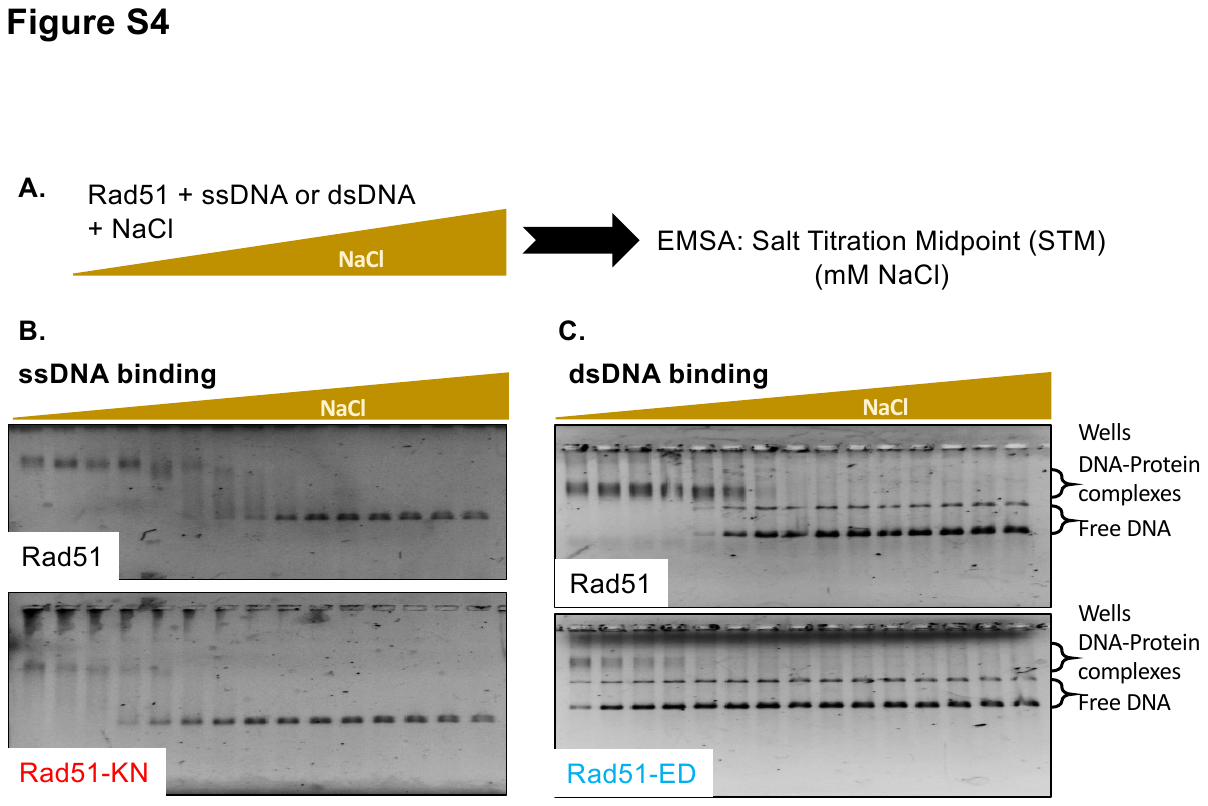

**Figure S4: DNA binding defects of Rad51-ED and Rad51-KN. A.** Reaction scheme of EMSA assays. **B, C.** Representative gels with ssDNA (B) and dsDNA (C).

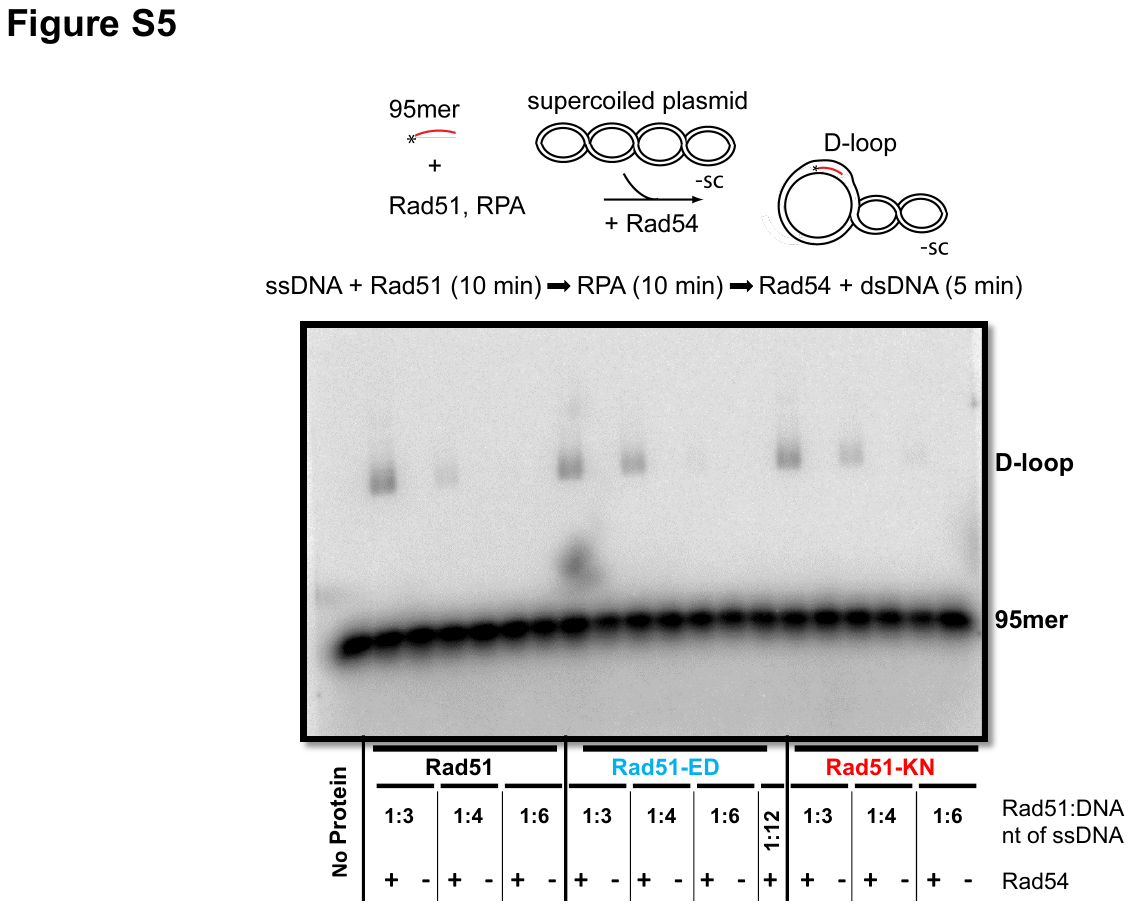

**Figure S5: D-loop formation by Rad51-ED and Rad51-KN depends on Rad54.** Qualitative D-loop assays with 5’-end labelled 95mer (olWDH566, see Table S3) and pUC19 dsDNA conducted for 5 min. Products were analyzed by gel electrophoresis and a representative gel is shown. The reactions were conducted at the Rad51 to nucleotide (nt) ssDNA ratios indicated in the presence and absence of Rad54.

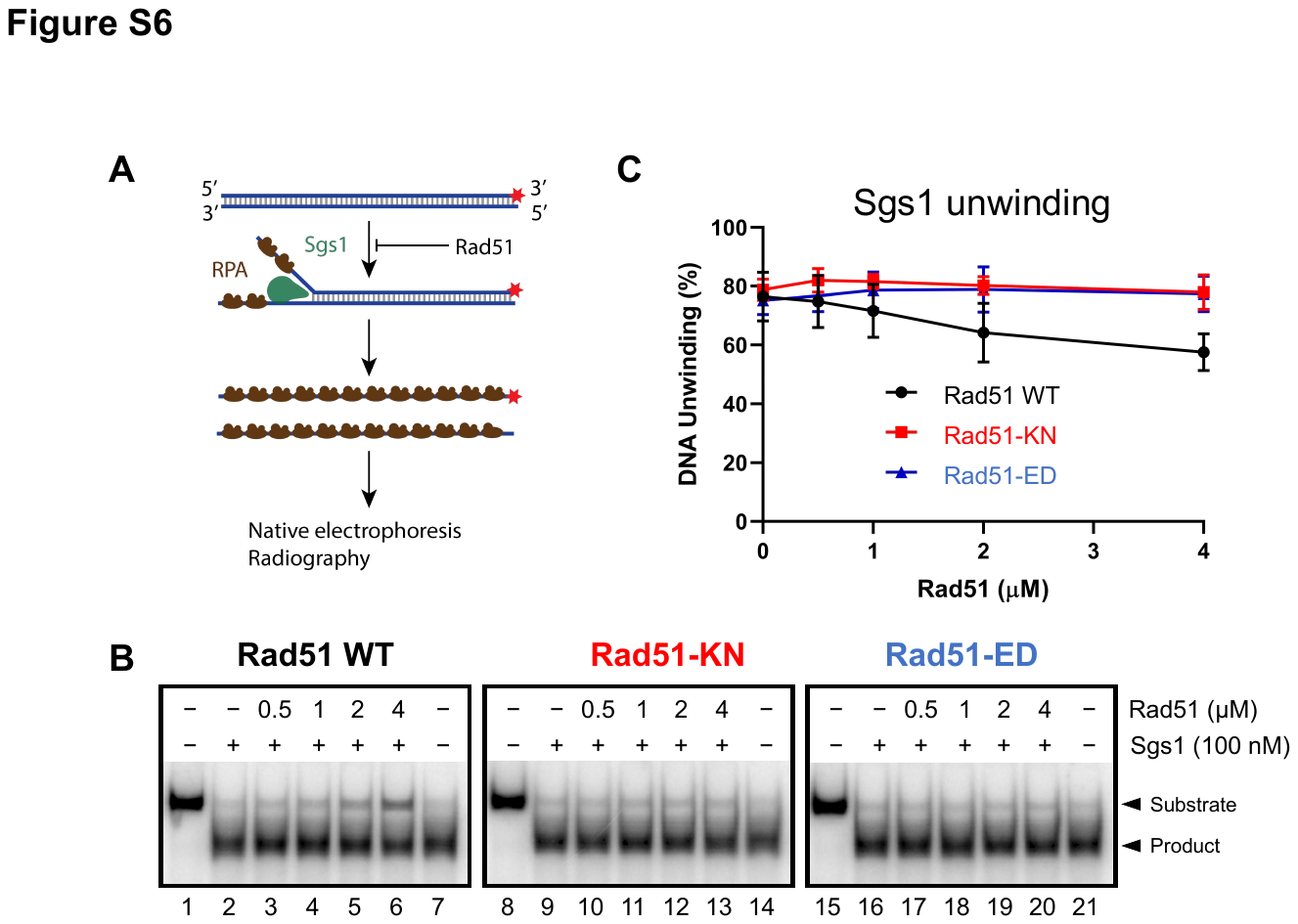

**Figure S6: Rad51-KN and Rad51-ED are defective in protecting dsDNA from unwinding by Sgs1. A.** Cartoon depicting the helicase assay with Sgs1 performed in panels B-C. The red asterisks represent the position of the radioactive label. **B.** Representative helicase assays with Sgs1 in the presence of increasing concentrations of Rad51 wild type and variants. In lanes 7, 14 and 21, the substrate was boiled to indicate the position of the ssDNA. **C.** Quantitation of experiments such as shown in B. Averages shown; error bars, range; n=2.

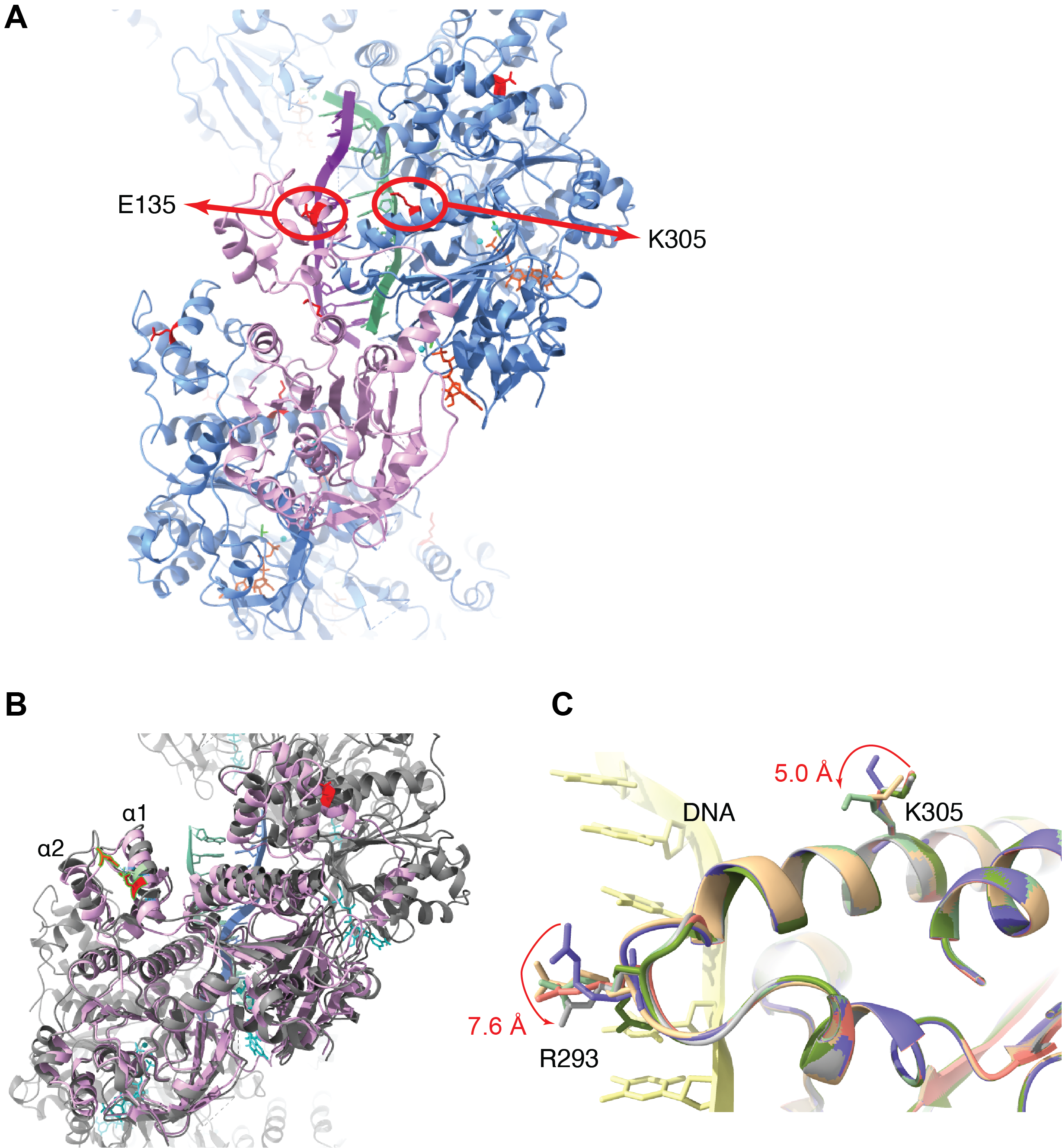

**Fig. S7. Superimposed wild type (experimental) and mutant (predicted) Rad51 structures. A**. Positions of E135 (red) and K305 (red) in the Rad51 filament structure (PDB: 9B2D) with modeled dsDNA. Rad51 protomers, alternating blue and purple. ADP, orange. AlF_3_, green. Magnesium, cyan. Modeled dsDNA, purple and green. **B**. Positions of locked α1 and α2 helices in the predicted Rad51-ED (purple) structure, in comparison to the wild type (gray) structure. ADP, AlF_3_, and magnesium are shown in cyan. Modeled dsDNA, blue and green. **C**. Dynamic movements of K305 and R293 in the Rad51-ssDNA cryo-EM structure.

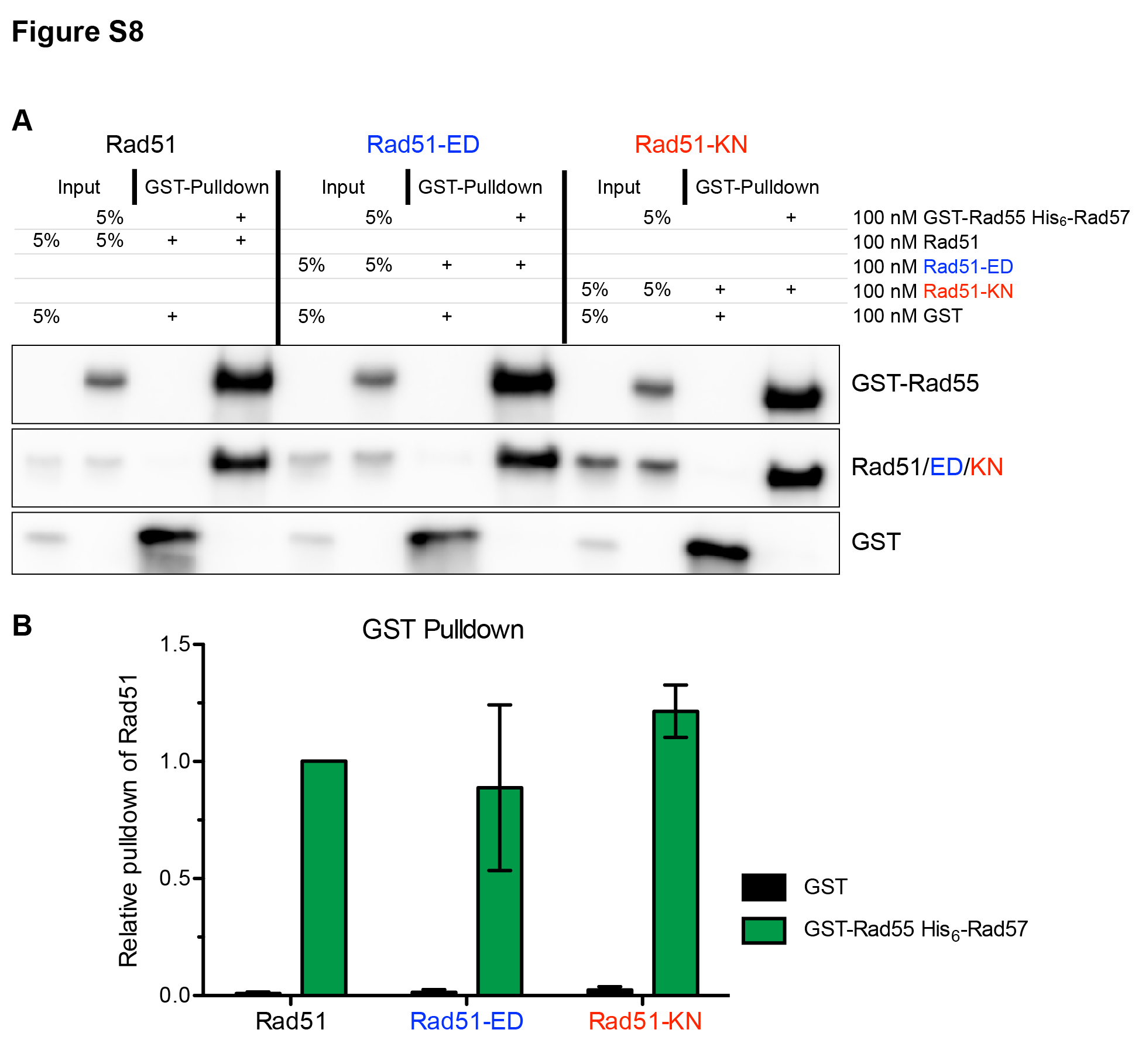

**Figure S8. Rad51-ED and Rad51-KN are proficient in their interaction with Rad55-Rad57. A.** Representative immunoblot of pulldown of Rad51 by GST-tagged Rad55 in complex with Rad57. **B.** Quantitation of pulldowns from n=3, shown are means with error bars representing 1 standard deviation.

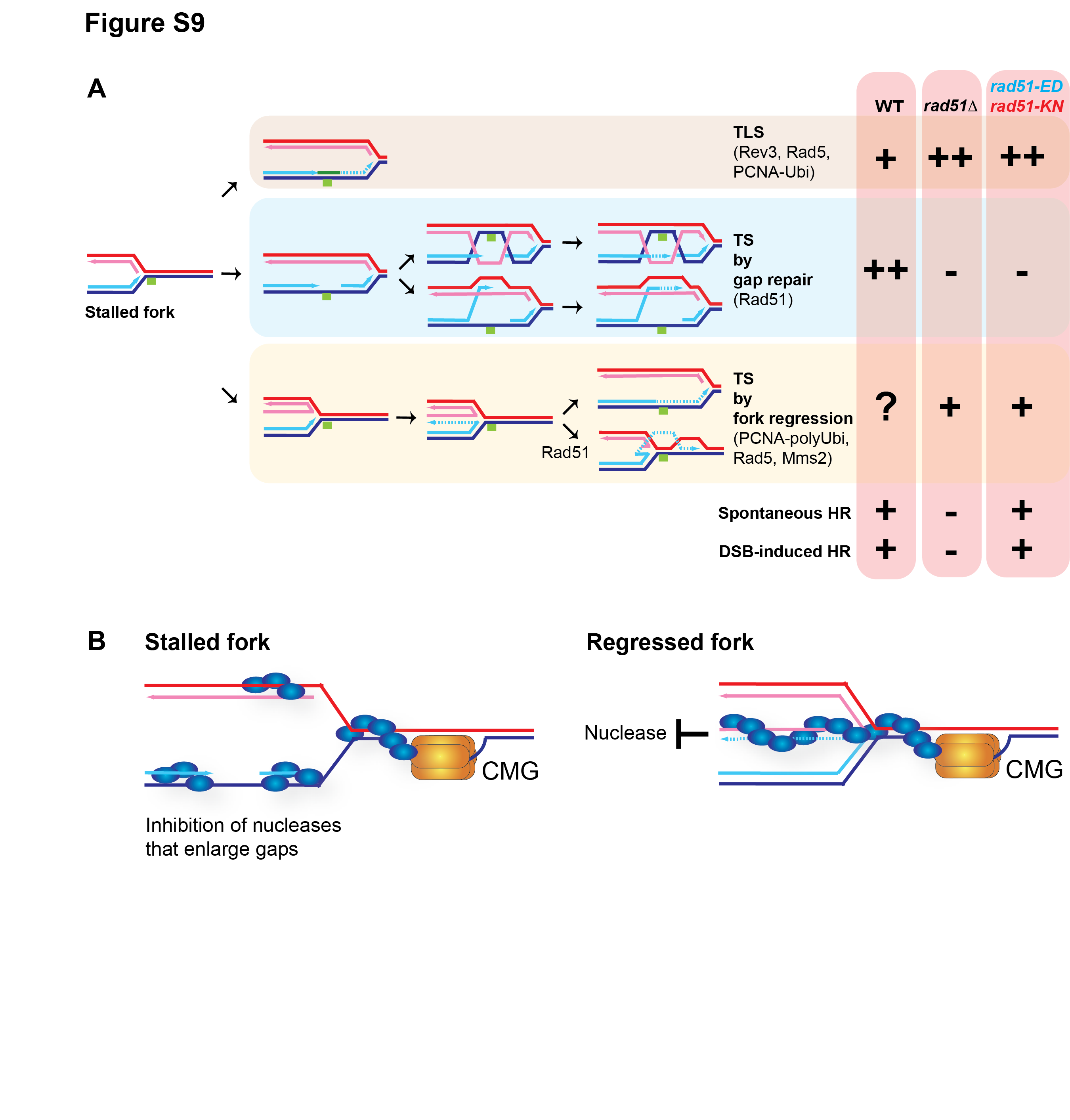

**Figure S9. Summary, interpretation, and model. A.** Representation of postreplication repair pathways and genes analyzed in this study, including translesion synthesis (TLS) which in budding yeast depends on PCNA ubiquitylation, Rad5, and Rev3. There are two different template switching (TS) pathways. First, an HR-dependent pathway of gap repair that can proceed either by gap invasion (top) or end invasion (bottom). Both pathways are expected to be dependent on Rad51. Second, fork regression which in budding yeast is expected to depend on PCNA poly-ubiquitylation, Rad5, and Mms2 in analogy to mammalian cells. **B.** Potential sites of dsDNA binding by Rad51 to protect from exonucleolytic degradation of gaps at stalled forks and the dsDNA end of the regressed fork. For more discussion see text.
